# Cytochrome P450 Enzyme Design by Constraining Catalytic Pocket in Diffusion model

**DOI:** 10.1101/2024.01.08.574609

**Authors:** Qian Wang, Xiaonan Liu, Hejian Zhang, Huanyu Chu, Chao Shi, Lei Zhang, Pi Liu, Jing Li, Xiaoxi Zhu, Yuwan Liu, Zhangxin Chen, Rong Huang, Jie Bai, Hong Chang, Tian Liu, Zhenzhan Chang, Jian Cheng, Huifeng Jiang

## Abstract

Although cytochrome P450 enzymes are the most versatile biocatalysts in nature, there is insufficient comprehension of the molecular mechanism underlying their functional innovation process. Here, by combining ancestral sequence reconstruction, reverse mutation assay and structure analysis, we identified five founder residues in the catalytic pocket of flavone 6-hydroxylase (F6H) and proposed a “three-point fixation” model to elucidate the functional innovation mechanisms of P450s in nature. According to this design principle of catalytic pocket, we further developed a de novo diffusion model (P450Diffusion) to generate artificial P450s. Ultimately, among the 17 non-natural P450s we generated, ten designs exhibited significant F6H activity and six exhibited a 1.3- to 3.5-fold increase in catalytic capacity compared to the natural CYP706X1. This work not only explores the design principle of catalytic pockets of P450s, but also provides an insight into the artificial design of P450 enzymes with desired functions.

## Introduction

Cytochrome P450 enzymes (P450s) are ubiquitous in nearly all living organisms, playing pivotal roles in various metabolic processes and pathways crucial for life, growth, and development^1^. As the most versatile biocatalysts in nature, P450s not only catalyzed more than 95% of the reported oxidation and reduction reactions^2, 3, 4^, but also are known as “Universal catalyst” in industrial applications due to the ability of selective oxidation of inert carbon-hydrogen bonds under mild conditions^5, 6^. Therefore, obtaining new P450s with better properties has become an important goal in the field of bioengineering^7, 8^. In spite of huge functional diversity, most P450s share the same catalytic mechanism^9, 10^ and similar structural scaffolds^4^. However, the catalytic pockets exhibited significant variability in P450s with different functions (Fig. S1)^11^. Moreover, the nonpolar composition and unique conformational flexibility of the substrate binding pockets are likely to enhance the capacity of these enzymes to modify their active sites and adapt to new substrates and selectivities^4^. Considering the high evolvability of P450s, directed evolution has been extensively employed in engineering P450s with better traits^12, 13, 14, 15^. However, this method often necessitates multiple rounds of random mutagenesis and high-throughput screening, making it challenging to exhaustively explore the potential protein space, whether in the laboratory or computationally^16^.

The rapid development of deep learning has opened up a new method to acquire novel P450s with desired characteristics. Even though impressive achievements have been witnessed in protein structure prediction^17, 18^, the desired functional design still is a big challenge^19, 20^. Recent developments in protein design leveraged by deep learning methods encompass a broad spectrum. These include designing sequences for fixed backbones^21^, variable backbone design^22^, as well as the direct generation of novel sequences and backbones within the natural protein space^23^. These models employ various architectures, including Convolutional Neural Networks (CNN), Graph Neural Networks (GNN), and Transformers, which are all instrumental in capturing the complex interactions between amino acids within a protein sequence^19^. The abundance of sequence and structure data contributes to these deep learning models surpassing the performance of traditional physical or statistical models^24, 25^. However, when considering functional design, it’s impossible to collect sufficient high-quality functional data to train a sophisticated model to create sequences with a desired function^26, 27^. Considering the current shortage, an approach that fuses knowledge-based techniques to scrutinize the design principles of natural P450s with powerful deep learning models to expand the natural protein sequence space, may be appropriate for designing new P450s. As our comprehension of the fundamental mechanisms that govern the evolution of the catalytic pocket for functional innovation in natural P450s remains limited, elucidating the process by which a particular P450 adopts a new function becomes crucial in designing a new one.

In this work, we used a flavone 6-hydroxylase (CYP706X1) from *Erigeron breviscapus* as an example, which belongs to the CYP706X subfamily and converts apigenin into scutellarein in the biosynthetic pathway of scutellarin (Fig. S2)^28^. Firstly, we determined the founder residues constituting the catalytic pocket responsible for the functional innovation of the P450 gene through ancestral sequence reconstruction, reverse mutation assay and crystallographic analysis. Then, we elucidated the design principle of catalytic pocket for the functional innovation by an in-depth structural analysis. Finally, we devised the P450Diffusion, an artificial P450 generative model, by integrating the catalytic pocket design principle with a denoising diffusion probabilistic model which has demonstrated outstanding performance in image generation^29^. With the P450Diffusion model, we successfully designed 10 artificial P450s with F6H activity, and one design outperforms the naturally best-performing gene about 3.5-fold, indicating the potential of P450Diffusion in the design of new P450 enzymes.

## Results

### Functional innovation of F6H in CYP706 family

Among the characterized P450s in CYP706 family, only the P450s in CYP706X subfamily could catalyze the flavonoid substrates, indicating that the F6H function may be de novo innovated in the ancestor of CYP706X subfamily (Fig. 1a and Fig. S3). Moreover, we found that the catalytic pocket’s configuration of CYP706X1 (i.e., EbF6H from *Erigeron breviscapus*) is totally different from other P450s in CYP706 family. The substrate apigenin even could not be properly positioned in other P450s with a C6-prone reactive state, which refers to the molecular configuration that is best suited for binding to the catalytic pocket of the enzyme and undergoing a reaction (Fig. S4). Therefore, it provides us an opportunity to decipher the constructive mechanisms for the formation of F6H’s catalytic pocket by comparing the neighboring genes in CYP706 family.

**Figure 1.**
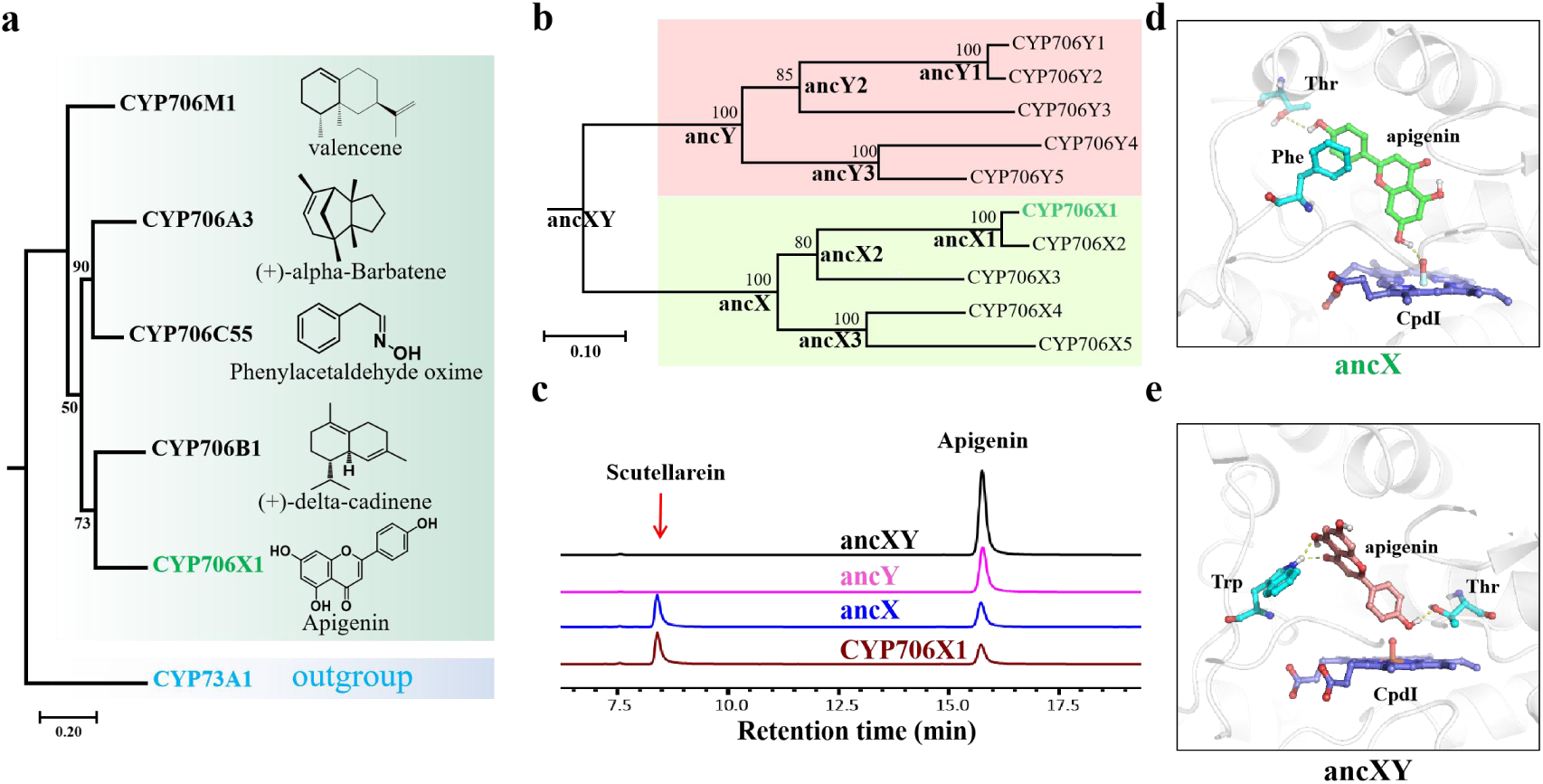
De novo innovation of F6H function in CYP706X subfamily. **(a)** Phylogenetic relationship of 5 characterized genes in CYP706 family. The CYP73A1 was set as an out-group. The maximum likelihood tree was constructed and all nodes received bootstrap support values from 100 replicates. **(b)** Phylogenetic tree of CYP706X and CYP706Y subfamilies. The inferred ancestral nodes are annotated with bold representations. CYP706X1 referred to F6H in *E. breviscapus*. **(c)** HPLC analysis of the fermented products of ancXY, ancY, ancX and CYP706X1. **(d and e)** Substrate-binding models of apigenin in catalytic pocket of ancX **(d)** and ancXY **(e)**. The dash lines represented the hydrogen bond interactions.

We compared the evolutionary trajectory between the CYP706X subfamily and the most closely non-functional CYP706Y subfamily using ancestral sequence reconstruction (Methods). By testing the function of the inferred ancestral P450s for all key nodes in the phylogenetic tree (Fig. 1b), most ancestral sub-nodes in CYP706X subfamily displayed significant F6H activity (Fig. 1c and Fig. S5). Conversely, the F6H function disappeared in both the common ancestor ancXY and the ancestor of CYP706Y subfamily (Fig. 1c). Thus, the F6H’s catalytic pocket should be originated when the CYP706X subfamily diverged from the common ancestor of CYP706X and CYP706Y (ancXY). To gain insight into the evolution of the catalytic pocket underlying functional innovation, we determined the crystal structure of ancX3, which was found to crystallize more readily after screening for crystallization conditions (Fig. S5, Fig. S6 and Table S1). Indeed, the binding mode of apigenin in the common ancestor of CYP706X subfamily (ancX) was obviously different from the non-functional ancXY, though they possessed very similar structural arrangement (RMSD < 1.0Å, sequence identity = 83%) (Fig. 1d and Fig. 1e). A strong pai-pai stacking and an obvious hydrogen bond are found to stabilize the substrate in a C6-prone reactive state in ancX’s catalytic pocket (Fig. 1d). However, the substrate in the non-functional ancXY is held in a non-C6-prone reactive state with the hydrogen bonds only by the surrounding residues like Trp and Thr (Fig. 1e).

### Founder residues for functional innovation of F6H

In order to clarify the molecular mechanism of forming the catalytic pocket with F6H function, we proposed to analyze the changes of amino acid compositions between catalytic pockets of non-functional ancXY and functional ancX. Within 8 Å range of the active center, 16 out of 48 residues are different (Fig. 2a). Interesting, when we replaced all of the 16 residues with the corresponding residues in ancX, the mutant (referred to as the ancXY-16) obtained F6H function (Fig. 2b). Given that not all residues in the catalytic pocket contributed significantly to substrate recognition and binding due to different locations of residues in three-dimensional space^30^, we attempt to find out the founder residues of the catalytic pocket in ancXY-16 by the reverse mutation assay (RMA) to eliminate non-essential residues (Fig. 2b).

**Figure 2.**
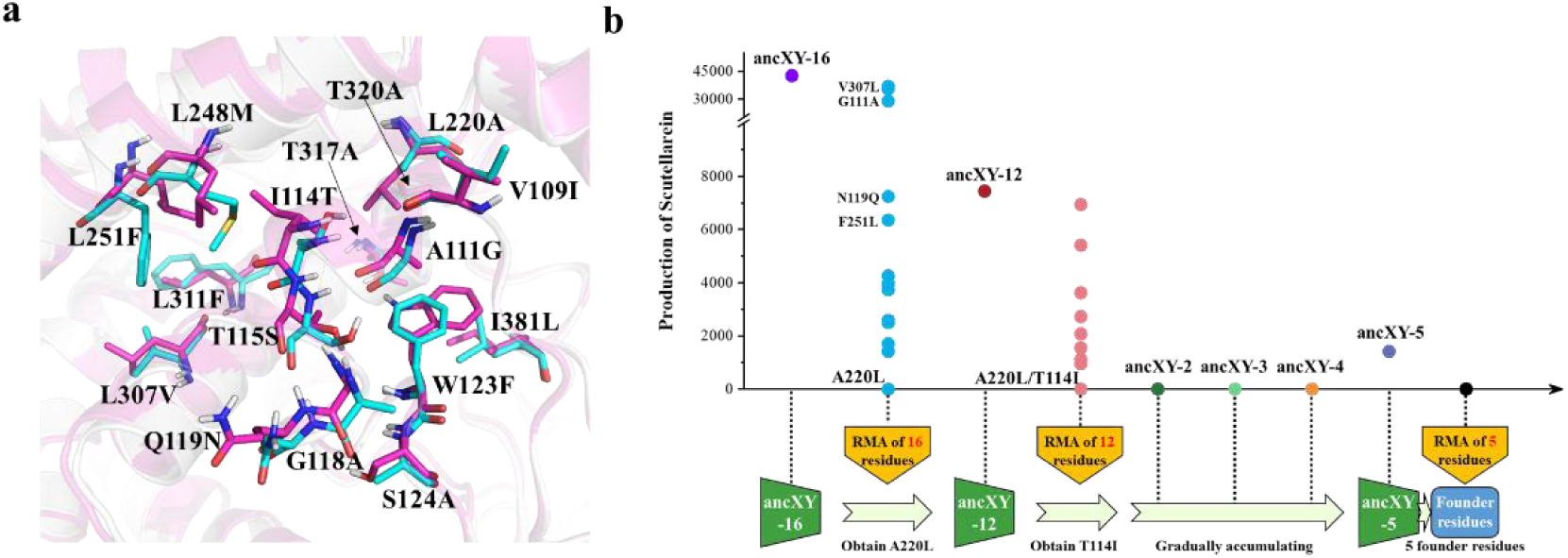
Reverse mutation assay for the identification of founder residues. **(a)** 16 different residues within the 8 Å range of active center of non-functional ancXY and functional ancX. All residues were represented as ball-and-stick model, and the residues of ancX and ancXY were color by cyan and magenta, respectively. **(b)** Process of RMA for the identification of founder residues. In the first round of RMA, one founder residue A220L was identified and four non-essential residues (V307L, G111A, N119Q and F251L) were eliminated; In the second round of RMA, the other founder residue T141I was identified; At last, three founder residues (i.e., W123F, L248M and T317A) were identified. ancXY-2, ancXY-3 and ancXY-4 referred to ancXY-L220A/I141T, ancXY-L220A/I141T/F123W and ancXY-L220A/I141T/F123W/M248L, respectively.

Firstly, RMA was respectively carried out on the 16 residues of ancXY-16 to clarify the effect of each residue on the catalytic activity. We found one of them (A220L) inactivated the ancXY-16, and 12 mutations significantly decreased the catalytic activity, but four mutations (i.e., G111A, N119Q, F251L and V307L) had less impact on the activity. Structural analysis showed that these four mutations were distant from the P450 catalytic center and did not involve in the changes in the residue’s intrinsic hydrophilicity/hydrophobicity (Fig. S7). Subsequently, we excluded these four mutations to construct the ancXY-12. RMA against the 12 residues of ancXY-12 showed one extra mutation (T114I) could destroy the function of F6H. We combined the two inactivating mutations (L220A and I114T) in ancXY to construct ancXY-2, however, it didn’t show F6H activity. Furthermore, we gradually added single mutation to ancXY-2 according to the order of the RMA mutational effect in ancXY-12. And finally, the constructed ancXY-5 (i.e., L220A, I114T, W123F, L248M, and T317A) displayed F6H activity, and each of the five reverse mutations in ancXY-5 deactivated the enzyme (Fig. 2b). The results showed that the mutations of the five amino acids play a founder role (referred to as founder residues in the following) in the F6H functional innovation process from ancXY to ancX. As to other 11 residues, the structural analysis showed that these mutations decreasing the catalytic activity might play auxiliary roles in the enzyme catalysis due to no direct interactions with the substrate apigenin (Fig. S7 and Fig. S8).

### The principle of catalytic pocket for functional innovation of F6H

We further interpreted the underlying mechanism of five founder residues for functional innovation through an in-depth analysis of the apigenin-binding model in ancXY-5 (Fig. 3a). The five founder residues could be divided into two parts according to their roles in protein structure. The first part included I114T, W123F and L248M which mainly contributed to fix or bind the apigenin. For example, the I114T introduced a hydrogen bond with 7’ hydroxyl of apigenin with an energy contribution of 0.66±0.10 kcal/mol (Methods, Fig. 3b). A null mutation of T114V in ancXY-5 also ascertained the indispensability of this hydrogen bond for the F6H function (Fig. S9). The W123F contributed to the apigenin binding (-3.14±0.37 kcal/mol) with an aromatic pai-pai stacking interaction to the phenyl ring of the apigenin and alleviated the spatial conflicts caused by ancestral tryptophan in the ancXY (Fig. 3c). The L248M, located in the substrate access gate, was not only involved in the substrate tunneling process (Fig. 3d, Video S1), but also contributed to the apigenin binding with a pai stacking to the phenyl ring of apigenin. The second part included L220A and T317A contributed to alleviate inappropriate interactions and space conflicts. The L220A alleviated the space conflict conducted by ancestral leucine and provided sufficient space for the placement of the B ring of substrate apigenin through the introduction of a small side chain (Fig. 3e). The T317A not only provided sufficient space for the placement of the A ring of apigenin but also avoided the wrong-orientation apigenin-binding mode shown in nonfunctional ancXY caused by a hydrogen bond between the hydroxyl group of threonine and the substrate (Fig. 3f).

**Figure 3.**
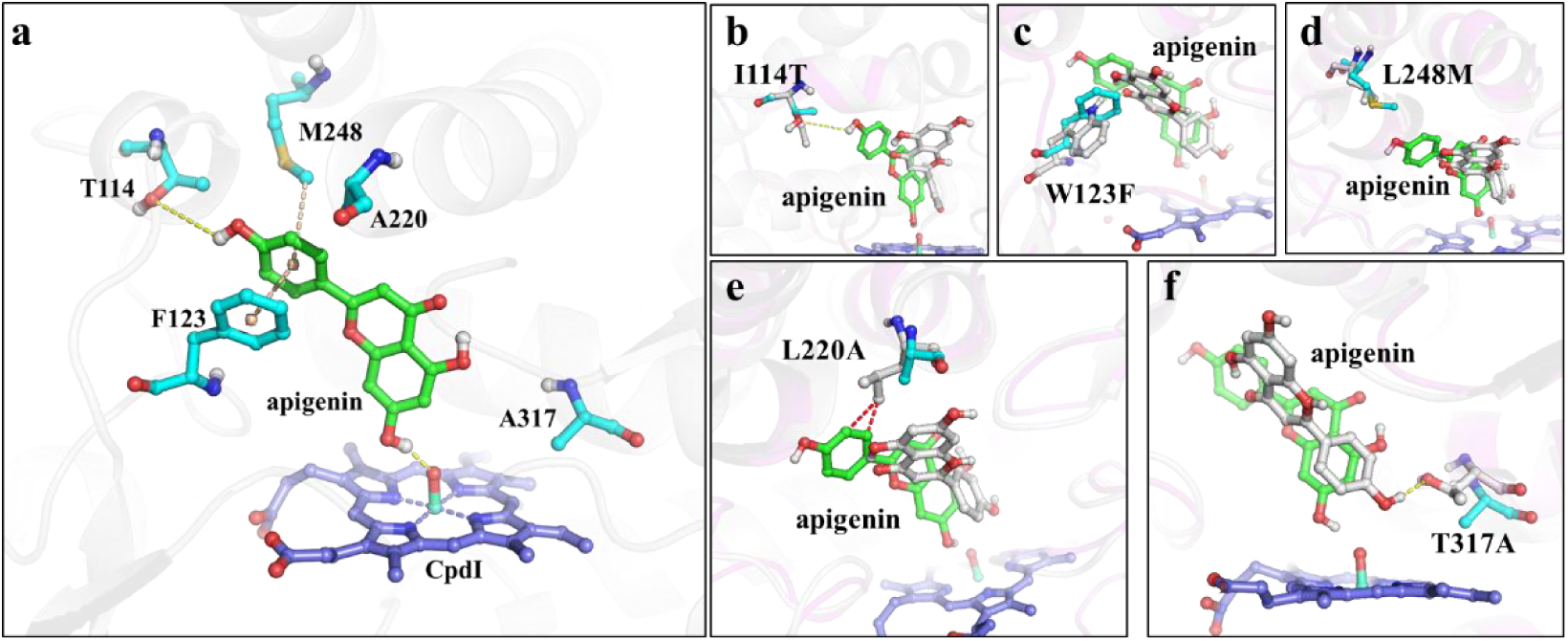
Contribution of five founder residues for forming the reactive near-attack conformation. **(a)** Spatial conformation of five founder residues (cyan), substrate apigenin (green) and CpdI (lightblue). **(b-f)** Comparison of each founder residue interacting with substrate in ancX and ancXY. The substrate apigenin in ancX and ancXY referred green and white, respectively. The founder residue in ancX and ancXY referred cyan and white, respectively.

Based on the mutations of five founder residues, it appears that, with an appropriate spatial capacity (provided by small side chain residues A220 and A317), the catalytic pocket evolved following a “three-point fixation” model. The “three-point fixation” refers to essential interactions with three pivots in apigenin including: 4’-OH of apigenin molecule (the first pivot) was fixed by the hydrogen bond from T114, the “B” ring of apigenin (the second pivot) was fixed by the pai stacking interactions from F123 and M248, and 7-OH of apigenin (the third pivot) was fixed by the hydrogen bond with CpdI iron-oxo moiety (Fig. S10). The model held the substrate apigenin in a reactive near-attack conformation (NAC), which maintained the relative orientation between the reaction site of apigenin and CpdI iron-oxo moiety at a favorable distance and angle (3.6 Å and 155°), thus serving to initiate the 6-hydroxylation reaction of apigenin in the catalytic process (Fig. S11). We propose that the “three-point fixation” model could serve as the design principle for the catalytic pocket responsible for the natural functional innovation of F6H, which also offers us the potential to de novo design P450s with the desired functions.

### Diffusion model-based designing of P450 with the specific function

Hundreds of thousands of P450 protein sequences collected in public databases offer us an opportunity to learn natural P450 sequence diversity and design new functional P450s^31^. Recent advancements in diffusion models have shown significant potential in enhancing the design of P450 enzymes with specific functions^29, 32^. Here, we proposed a P450 Sequences Diffusion Model (P450Diffusion) to de novo design P450s with a desired function by combining the diffusion model with the design principle of F6H catalytic pocket (Fig. 4a). P450Diffusion mainly consists of two models (i.e., pre-trained and fine-tuning diffusion models). Firstly, 226,509 natural P450 sequences were collected to train a pre-trained P450 sequence diffusion model. This pre-trained model consists of two subprocesses: a forward diffusion subprocess, which gradually adds Gaussian noise to the representation of P450 sequence until it becomes random noise, and a reverse generation subprocess, which starts from random noise and gradually de-noises the representation of P450 sequence to generate a new P450 sequence. After 150,547 training rounds, the pre-trained diffusion model could generate a wide variety of sequences, with similarities to natural sequences ranging from 20% to 50%. Secondly, 19,202 P450 sequences with appreciable similarity to CYP706X subfamily were used to fine-tune the pre-trained diffusion model for ensuring that the generated sequences have a similar structural backbone to the F6H. Besides, the five founder residues including T114, F123, A220, M248 and A317 were constrained to ensure the reproduction of the “three-point fixation” design principle in de novo generated sequences. The model integrating training set fine-tuning with constrained generation was referred to as the fine-tuning diffusion model.

**Figure 4.**
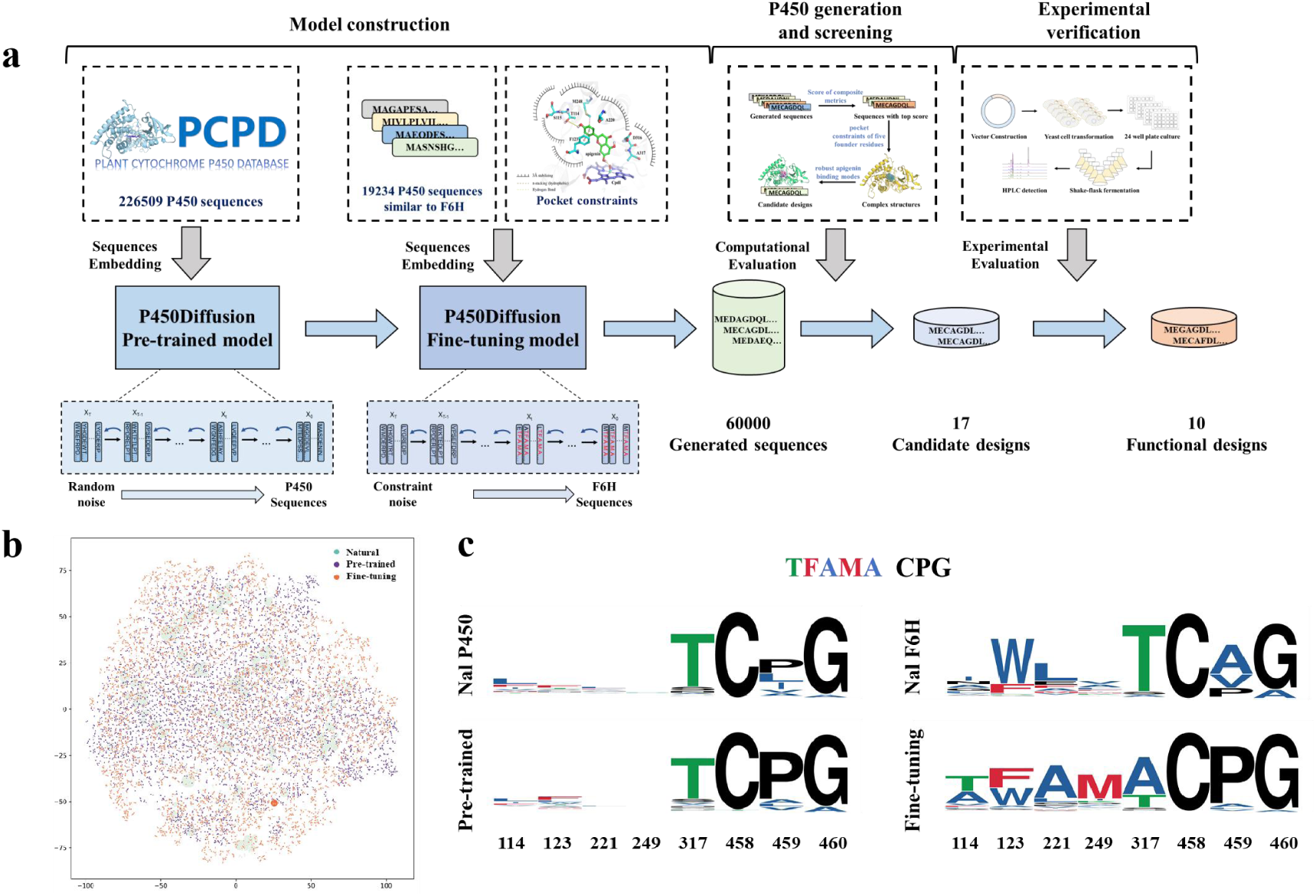
P450Diffusion de novo design new P450 Processes from Scratch. **(a)** The design process for the new P450 includes P450Diffusion model construction (a pre-trained model and a fine-tuning model), sequence generation and screening and experimental verification. The generated sequences were screened and evaluated to obtain the candidate sequences for experimental verification. **(b)** t-SNE embedding of natural, pre-trained model and fine-tuning model generated sequences. The protein sequence space was visualized by transforming a distance matrix derived from k-tuple measures of protein sequence alignment into a t-SNE embedding. Dot sizes represent the 50% identity cluster size for each representative. **(c)** The distribution of five founder residues and control residues (CPG) among natural and generated P450s is illustrated in the WebLogo using multiple sequence alignment (MSA)^58^. This visualization incorporates data from four distinct sources: the P450Diffusion pre-trained model dataset (Nal P450), sequences generated by the P450Diffusion pre-trained model (Pre-trained), the P450Diffusion fine-tuning model dataset (Nal F6H), and sequences generated by the P450Diffusion fine-tuning model (Fine-tuning).

Furthermore, we used the fine-tuning diffusion model to generate a total of 60,000 non-natural P450 sequences, which share about 50% average amino acid identity to that of the natural sequences. In comparison with natural P450s, the generated sequences not only have a highly similar distribution of Shannon entropies for each position in multiple sequence alignments, but also display very consistent residue-residue co-evolution patterns and physicochemical properties (Fig. S12 and Fig. S13). However, the generated sequences can be grouped into smaller clusters and interpolated between the natural sequence clusters, indicating that the generated sequences have higher diversity than natural P450s (Fig. 4b). It is noteworthy that the sequences generated by the fine-tuning P450Diffusion model form a larger cluster, exhibiting greater similarity to the CYP706X subfamily, thereby demonstrating the effectiveness of the fine-tuning model. Besides, we compared the distribution of five founder residues among natural and generated P450s (Fig. 4c). It is found that except the threonine (T) in position 317, other positions are highly variable in natural and generated P450s from pre-trained model, even in natural P450s from CYP706 family. However, all of five founder residues are relatively conserved in the generated P450s from fine-tuning model, indicating that the P450Diffusion possessed the capability of generating sequences with an amino-acid distribution similar to that of natural F6H on the basis of constrained five founder residues.

### Experimental verification and structural insights of de novo generated P450s

Finally, we experimentally tested whether the generated sequences from P450Diffusion were true P450 enzymes, and performed F6H function. In order to accurately obtain functional sequences from numerous designs, we conducted virtual screening on 60,000 generated sequences based on three specific criteria: the computational scores of composite metrics for assessing the quality of generated sequences, the 3-dimensional pocket constraints of the five founder residues, and the robustness of the apigenin binding modes (details in Methods, Fig. 4a). 17 designs with sequence identities ranging from 70% to 87% to CYP706X1, were retained by the virtual screening, then synthesized and expressed in yeast expression systems (Table S2). The recombinant yeasts were cultivated for four days by feeding apigenin as substrate and HPLC analysis revealed ten designs with significant F6H activity (Fig. 5a). Surprisingly, there are six designs exhibited a 1.3- to 3.5-fold increase in scutellarein production compared to CYP706X1 (Fig. 5b). The four remaining active designs also displayed comparable activities with other natural F6H enzymes (i.e., Cnan706X and Lsal706X). Therefore, the results indicated that the P450Diffusion could not only capture the fundamental design principle of F6H catalytic pocket and effectively generate P450s sequences with F6H activity, but also selected out the better P450 enzymes compared to natural sequences from the P450 sequence space.

**Figure 5.**
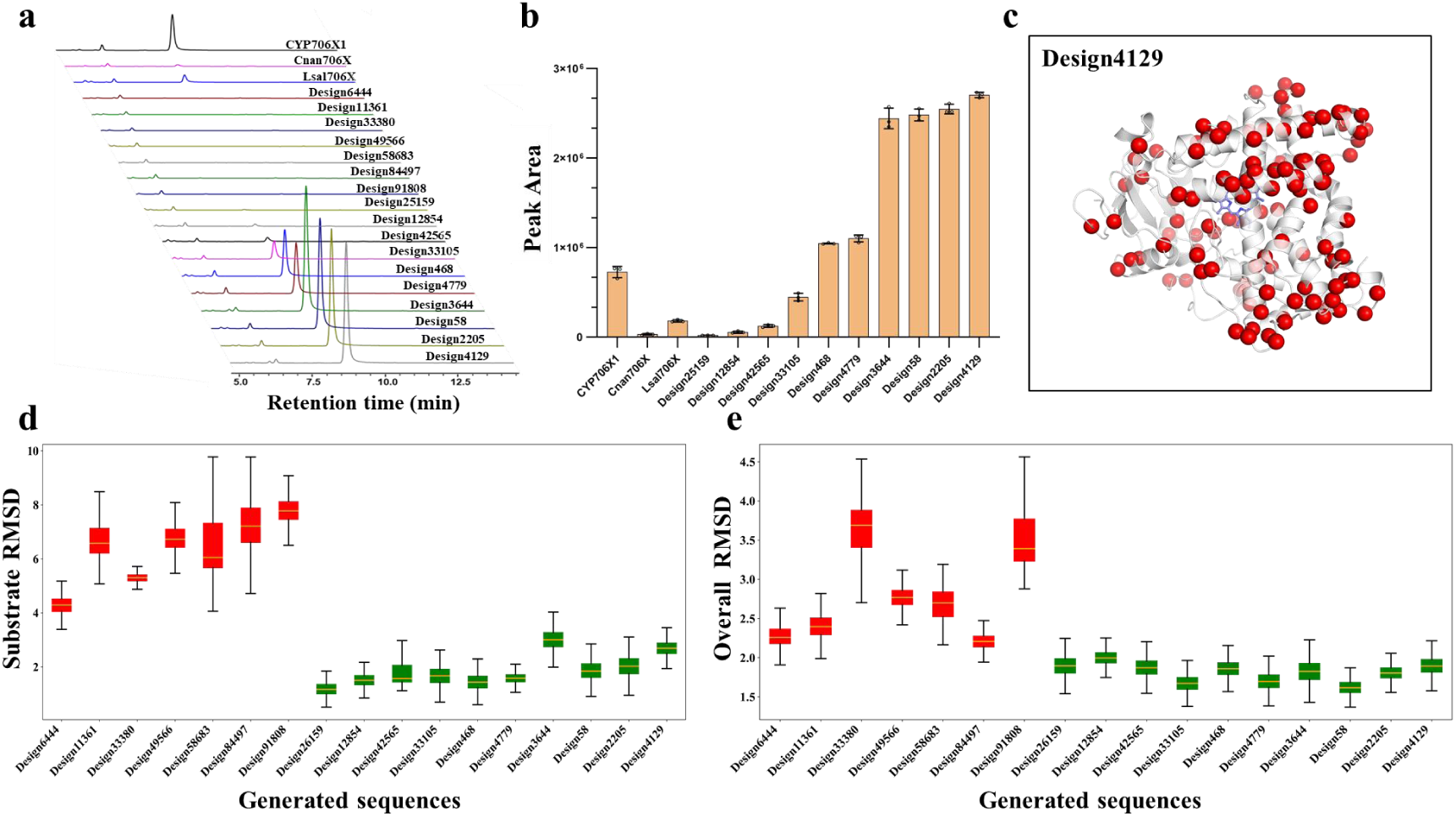
Experimental verification and structural insights of de novo generated P450s. **(a)** The product scutellarein peak area of 17 designs, compared with natural Cnan706X, Lsal706X and CYP706X1. Different colors were assigned to different proteins. **(b)** The histogram displays the peak areas of products associated with functional designs, with CYP706X1 used as the control group. **(c)** The structural distribution of mutations in Design4129 was compared to that in CYP706X1, with mutations represented as red spheres. **(d)** The boxplot illustrates the substrate RMSD values across long-term MD simulations, with active designs depicted in green and inactive designs in red. **(e)** The boxplot represents the RMSD values for the overall protein structure across long-term MD simulations, with active designs shown in green and inactive designs in red.

Meanwhile, in order to further analyze the other seven designs without F6H activity, we first test whether the seven designs can be soluble expressed in yeast expression systems by integrating green fluorescent protein at the C-terminal. All recombinant proteins successfully showed green fluorescence, demonstrating that seven designs folded correctly in the yeast expression systems (Fig. S14).

Furthermore, we presented a structural perspective on the active designs as well as the distinctions between the active and inactive ones. The structural analysis reveals that no substantial mutations in the protein-substrate binding pockets between active and inactive designs except the surface of the protein structure (Fig. 5c), and substrates bind to catalytic pockets of all designs in a manner highly similar to natural CYP706X1 (Fig. S15). However, long-term Molecular Dynamics (MD) simulations have demonstrated significantly weaker binding stability of the substrate apigenin in the inactive designs when compared to the active ones (Fig. 5d). This discrepancy likely serves as the primary reason for the inactivity observed in these seven designs. Besides, we observed that the overall protein structures of the active designs appear to exhibit greater stability than the inactive ones following extensive MD simulations (Fig. 5e). Notably, significant structural fluctuations are observed, particularly within the sequence ranges of 220-230 and 390-410, as illustrated in the inactive designs (Fig. S16). For instance, in Design33380, the R229K mutation disrupts the salt bridge with E251, while the S230P mutation causes a break in the alpha-helix structure (Fig. S17). And in Design91808, the S407L mutation break the hydrogen bond with the backbone of A51, resulting in a less stable protein backbone than observed in active designs (Fig. S18). These results imply that the amino acid mutations on the surface of the protein could lead to a reduction in the global stability of the protein, which further leads to substrate binding instability and ultimately to the loss of activity of the designs. This analysis provided us with valuable insights for future improvements of the P450 generative model.

## Discussion

Nature has evolved an amazing array of enzymes to catalyze biological functions and enabled living systems to face diverse environmental challenges^33^. Gene duplication contributes most to the generation of new enzymes^34^, especially for cytochrome P450s, which evolve to the largest enzyme family for plant metabolism by widespread whole-genome and tandem duplications^7, 35, 36^. Although most duplicates are lost or subfunctionalized by purifying or neutral selection, neofunctionalization often happened in P450 evolution due to high plasticity and variability of catalytic pockets^37, 38^. The evolutionary trajectory of P450’s functional innovation have attracted researchers’ attention for a long time^39, 40, 41^ and the previous researches were mainly focused at the gene level^42, 43, 44, 45, 46^ or residue level^47, 48^. In this study, based on ancestral sequence reconstruction, RMA and structural analysis, we suggested the “three-point fixation” model as the design principle of catalytic pocket which played a pivotal role in the functional innovations of F6H function.

The “three-point fixation” model seems to be a general principle for the substrate binding in P450’s catalytic pocket, such as the camphor binding in P450cam^49^ and N-palmitoyl glycine binding in P450BM3^50^. Similar fixation rules could also be found in the general enzymatic catalysis where the substrates or catalytic residues are held in the catalytic pockets^51^, even as a term commonly used in medicine and architecture^52^. It is worth mentioning that besides the “three-point fixation” model, nature also evolved other catalytic pocket design principles for functional innovations in P450s. For example, the SbaiCYP82D4, as the isoenzyme of CYP706X1, have evolved to a completely different catalytic pocket configuration for flavone 6-hydroxylation^53, 54^. The catalytic pocket of SbaiCYP82D4 consisted of more residues with strong hydrophobicity, and no obvious hydrogen bond was found between surrounding residues and substrate apigenin, making the substrate binds in an “oblique binding” orientation (Fig. S19), which is distinguished with the “vertical binding” orientation in CYP706X1. Although a different substrate binding model was found in SbaiCYP82D4, the substrate apigenin also formed a reactive conformation in a NAC model to enable the initiation of the catalytic reaction. This fact indicated that substrates in P450s could be held in favorable orientations with different fixing rules under the premise of sufficient space and suitable shape for the placement of the substrate.

The rapid development of deep learning has witnessed many impressive achievements in protein structure design, while the desired functional design still is a big challenge^55, 56, 57^. Our research provides a novel strategy for the de novo design of P450s with specific function by coupling the design principle of catalytic packet with deep learning model. In this study, non-natural P450s with F6H function were successfully designed by integrating the “three-point fixation” model with a denoising diffusion probabilistic model. The structural analysis of active designs suggested that the design principle of F6H catalytic pocket has been fully incorporated into the deep learn model. Furthermore, the structural insights between active and inactive designs suggest that mutations on protein surface may be the fundamental factors contributing to the inactivity or reduced activity of designed sequences, providing us with valuable insights for future improvements of the P450 generative model. There are more structure or sequence-based features should be considered, like the substrate-tunneling feature, the overall stability of protein, and so on.

In general, the current work provides insights into the principle of pocket design in the P450 functional innovations and offers a potential research paradigm for the de novo design of P450 enzymes with desired functions. With the increasing of in-depth investigated P450s, more catalytic pocket design principles would be deciphered and facilitated the design of P450s with novel and desired functions.

## Methods

### Phylogenetic analysis and ancestral sequence reconstruction

The P450 sequences of CYP706 subfamilies were selected from the previous study^28^, including ten P450s of CYP706X/Y subfamilies for ancestral sequence reconstruction, and a P450 of CYP706W subfamily as an out-group. The transmembrane domains of P450 sequences were annotated with the TMHMM package^59^. Using the crystal structures of CYP76AH1 (PDB ID: 5YLW), a structural information-based sequence alignment of the P450s deprived of N-transmembrane region were generated by Expresso^60^. Poorly aligned regions (N- and C termini) were trimmed. Then a phylogenetic ML tree was created with the RAxML^61^. All protein sequences of ancestral nodes were deduced using FastML^62, 63^. The N- and C-terminal amino acids include transmembrane domain derived from CYP706X1 were added to each ancestor. Ultimately, we obtained the most probable ancestor of CYP706Y subfamily (ancY) and CYP706X subfamily (ancX), the common ancestor of two subfamilies (ancXY), and all sub-ancestors of CYP706Y subfamily (ancY1, ancY2 and ancY3) and CYP706X subfamily (ancX1, ancX2 and ancX3) in the sub-nodes of the phylogenetic tree (Fig. 1b). The ancestral sequences are available in Supplementary information.

### Crystallization and Structure Solution

Initial crystallization screening was performed using the sitting-drop vapor-diffusion method with commercial crystal screen kits at 16 ℃. The ancX3 protein at concentration 10 mg/mL in buffer (2 mM KH_2_PO_4_, 8 mM K_2_HPO_4_, 500 mM NaCl, 0.2 mM EDTA, 1 mM DTT, 10% (v/v) glycerol and pH 7.4) was used in the initial crystallization screening to determine the crystallization condition. The ancX3 protein was mixed with precipitant solution at a drop size of 0.6+0.6 μL against the reservoir containing 50 μL precipitant solution. The crystals grew from the mixture with the precipitant solution consisting of 1.34 M NaCl, 13.4% (w/v) PEG3350, 0.1 M MgCl_2_, 0.1 M imidazole and pH 6.5. The crystal optimization was performed using the hanging-drop vapor-diffusion method at 16 ℃ against the reservoir containing 0.5 mL of the precipitant solution. The drops contained 2 μL precipitant solution, 2 μL ancX3 protein and 0.2 μL of additive solution (40% v/v Polypropylene glycol P400) from Hampton additive screen kit.

Crystals of ancX3 were mounted from the crystallization drops in nylon loops and flash-frozen in liquid nitrogen using cryoprotectant consisting of 1.34 M NaCl, 13.4% (w/v) PEG3350, 0.1 M MgCl_2_, 0.1 M imidazole, 25% (v/v) glycerol, pH 6.5. Diffraction data (λ = 0.97918 Å) were collected on beamlines 17U1 at Shanghai Synchrotron Radiation Facility for IFS crystals. Diffraction images were indexed, integrated and scaled using the XDS program. Details of the data-collection statistics are summarized in Table S1.

The structure of ancX3 was solved by molecular replacement with the structure of CYP76AH1 (PDB code: 5YLW) as search model^64^. Iterative model building and refinement were performed using COOT and PHENIX, respectively. Coordinates and structure factors have been deposited with the PDB under accession id 8JC2.

### Structural modelling and molecular docking

The 3D models of all P450s and ancestral proteins are predicted by the local ColabFold algorithm through inputting the crystal structure of ancX3 as one of templates^65^. The Cartesian coordinates and atom charges of CpdI was obtained from a published data^66^. The structure of substrate apigenin was obtained from PubChem^67^, and assigned with AM1-BCC charges^68^. An ensemble of different conformations of the substrate were generated by enumerating these under OpenBabel^69^. Substrate rotamers were extensively sampled around the C2-C1’ axis with 5° intervals. The mol2 formatted CpdI and apigenin were parameterized with molfile_to_params.py script. Before molecular docking, the protein structure complex with CpdI species was firstly sampled and minimized by the RosettaRelax protocol without constraints^70,71^. Then the apigenin was docked into relaxed structures using RosettaLigand^72, 73, 74^.

Distance restraints were added between the Fe ion and ligated cysteine (2.3 Å +/– 0.1 Å), between carboxylate groups of heme and arginines (2.2 Å +/– 0.4 Å) in Rosetta-Scripts^75^. Each run of 100,000 models were generated with the MPI^76^ version of RosettaLigand and the top 100 models with lowest REU were clustered with Calibur^77^, and the structures with the lowest binding free energy (interface_delta) were selected as our final docking models. The Rosetta scripts and option files for RosettaLigand and are available in Supplementary information.

### MD simulations and MM-PB/GBSA

Our target models with CpdI and substrate molecules were set as the initial structures for MD simulation. The protein structures were prepared with the pdb4amber application in Amber20 package^78^. The force field for the CpdI species was taken from a published data^66^. The partial atomic charges and missing parameters for substrate apigenin were generated by Antechamber with AM1-BCC charge model^79, 80^. A few Na^+^ ions were added to the protein surface to neutralize the total charge of the system. Finally, the resulting system was solvated in a rectangular box of TIP3P waters extending up to minimum cutoff of 12 Å from the protein boundary. The Amber ff14SB force field was employed for all the proteins in MD simulations.

After proper parameterizations and setup, the resulting systems were minimized with two steps (the first step with 5,000 steps of steepest descent and 10,000 steps of conjugate gradient, the second step with 10,000 steps of steepest descent and 30,000 steps of conjugate gradient) to remove the poor contacts and relax the systems. The systems were then gently annealed from 0 to 300 K under the NVT ensemble for 50 ps with a restraint of 5 kcal mol^−1^ Å^−2^. Subsequently, the systems were maintained for a total of five rounds of density equilibration of 20 ps in the NPT ensemble at a target temperature of 300 K and a target pressure of 1.0 atm using the Langevin thermostat^81^ with a restraint of 1 kcal mol^−1^ Å^−2^. Totally five rounds of density equilibration relaxed the system to achieve a uniform density after heating dynamics under periodic boundary conditions. Thereafter, we removed all of the restraints applied during heating and density dynamics and further equilibrated the systems for ∼2 ns to get a well-settled pressure and temperature for conformational and chemical analyses. This was followed by a MD production run for 100 ns for each of the systems. During all of the MD simulations, the covalent bonds containing hydrogen were constrained using SHAKE^82^ and particle-mesh Ewald^83^ was used to treat long-range electrostatic interactions. All of the MD simulations were performed with the GPU version of the Amber 20 package.

The python script mmpbsa.py^84^ in Amber20 package was used in this research to analyze the binding free energy of apigenin. According to the systematic research of Hou et al., the inclusion of the conformational entropy may be crucial for the prediction of absolute binding free energies but not for ranking the binding affinities of similar ligands^85^. The binding free energy analysis implemented here just for analyzing the interaction energy contribution of each key residue. Therefore, the change of conformational entropy upon ligand binding has been ignored in our calculation because of expensive computational cost and low prediction accuracy. The calculation procedure mainly referred the MMPBSA protocol in AMBER tutorial websites (http://ambermd.org/tutorials/advanced/tutorial3/section1.htm).

### Building and training the P450 Sequences Diffusion Model (P450Diffusion)

Denoising diffusion probability models (or diffusion models, for short) work by applying a Markov process to corrupt the training data by successively adding Gaussian noise, then learning to recover the data by reversing this denoising process^86^. We adapt this framework to generate protein sequences, introducing necessary modifications to encode the discrete protein sequences into a vector of a specific length. We used physicochemical character-based schemes, the principal components score Vectors of Hydrophobic, Steric, and Electronic properties (VHSE8)^87^, to encode protein sequences. The P450 Sequences Diffusion model (P450diffusion) is composed of a U-Net with self-attention layers and features a classical U-shaped structure with down-sampling and up-sampling blocks.

To build the P450Diffusion, we screened and analyzed all potential P450s from a published P450 database^31^ and public databases, filtering out sequences with a length greater than 560 and resulting in 226,509 sequences to form the training dataset. Then we encode the training dataset, where each amino acid in the protein sequence is encoded as an 8-dimensional vector, and each batch protein sequence is encoded as a 64×1×560×8 vector. Here 64 is the batch size equal to the number of samples in the training data; 1 represents the channel size; 560 represents the maximum length of the protein sequence; 8 represents the VHSE8 encode vector for each amino acid in the protein sequence. If the protein sequence is shorter than 560, we add gaps until it reaches a length of 560. In this case, we assign a vector of eight zeroes as the encoding for gaps. Then we started to train the pre-trained P450 sequence diffusion model. After 150,547 training steps, the loss functions of the pre-trained diffusion model converged and the model was obtained. (Fig. S20a).

In order to generate sequences with F6H function more effectively, we fine-tune the pre-trained diffusion model with the filtered dataset by selecting sequences with more than 30% amino acid identity to the CYP706X subfamily and clustering them with 90% sequence similarity. Finally, a total of 19234 sequences formed a fine-tuning dataset. Meanwhile, we assigned different sample weights to 30 sequences from the CYP706X subfamily and other sequences in the fine-tuning dataset. The sampling weight ratio between the 30 sequences from the CYP706X subfamily and other sequences was 600:1. The P450Diffusion was obtained after 150,500 training steps (Fig. S20b).

The P450Diffusion architecture to generate P450 sequences was based on the diffusion model. The diffusion model is composed of a U-Net with self-attention layers. The main difference with traditional U-Net is that the up-sampling and down-sampling blocks support an extra timestep argument on their forward pass. This is done by embedding the timestep linearly into the convolutions. In the training process, the network takes a batch of noisy protein sequences of shape (batch size, channels, height, width) and a batch of noise levels of shape (batch size, 1) as input, and returns a tensor of shape (batch size, channels, height, width). In this model, we used a mean squared error loss (MSELoss) function and optimized the networks with the AdamW algorithm, setting the learning rate to 2e-4. Our model was implemented in PyTorch and trained on 6 GeForce RTX 3090 systems for about 150,000 steps, which took approximately 63 hours.

### Computational evaluation and structure-based virtual screening for generated sequences

Three criteria were used to screen the generated sequences in silico to improved experimental validation success rates: the computational scores of composite metrics for assessing the quality of generated sequences, the 3-dimensional pocket constraints of the five founder residues, and the robustness of the apigenin binding modes. Details are as follows.

We used random protein sequences of length 560 with the five founder residues as the starting sequence for the diffusion model sample. In the reverse diffusion process, we perform 600 steps of denoising the 60,000 starting sequences to obtain 60,000 generated sequences. In order to increase the likelihood that the generated sequences would function as F6H, we evaluated the generated protein sequences using a variety of computational metrics, including esm-1v^88^, Alphafold2^18^, ProteinMPNN^89^, and others^90^. Firstly, the 60,000 generated sequences were screened by the sequence motif constructed by the five founder residues, and 77 sequences were filtered out. Secondly, both the 77 generated sequences and the F6H sequences were scored for esm-1v, and then the top 33 sequences in the esm-1v results were selected for alphafold2 structure modeling. Thirdly, the constructed structures and sequences were evaluated using ProteinMPNN, and the top 19 designs were selected based on their ProteinMPNN scores, which were higher than that of CYP706X1 (-1.63). Fourthly, substrate apigenin and CpdI were docked into constructed structures using RosettaLigand and the substrate-binding models were obtained based on binding affinity (interface_delta_X); Subsequently, MD simulations were performed to evaluate the overall structure stability and binding pocket stability for each designed sequences; Finally, the substrate-binding structures that meet catalytic pocket constraints constituted by founder residues and maintain stable substrate binding modes were chosen as candidate sequences for experimental verifications (Fig. S21).

### Cloning construction and products detection

Chemicals and media used in this study were exhibited in supplementary materials. All primers used in this study are listed in Table S3. All strains and plasmids are listed in Table S4. The protein sequences and DNA sequences can be found in supplementary information. Nucleotide sequences of ancXY, ancX, ancX1, ancX2 and ancX3, ancX-16 were codon optimized for *Saccharomyces cerevisiae* and synthesis by Genscript, China. Subsequently, the gene fragments, ATR2 (P450 reductase from *Arabidopsis thaliana*) and the head-to-head promoters (pPGK1-pTDH3) were cloned into the vector Y22-TC using the Minerva Super Fusion Cloning Kit (US Everbright Inc., China). The assembly system was transformed into DMT competent cells and the sequences assembled successfully were verified by further sequencing. For mutants constructing, mutation sites were introduced by the mutant primers which listed in Table S3 and used the same method for recombinant vectors assembly. The nucleotide sequences of P450 designs were codon optimized for *S. cerevisiae* and subcloned between PGK1 promoter and CYC1 terminator of Y22-PE by Genscript, China.

Due to the functional expression of P450 enzyme needed an auxiliary reductase partner (CPR), the ATR2 from *Arabidopsis thaliana* was cloned into expression vector YCplac33-TP which contained a TDH3 promoter and a PDC1 terminator and named Y33-ATR2. The plasmid Y33-ATR2 was preserved in our laboratory. The recombinant vectors containing P450 enzymes designed by deep learning were separately co-transformed with Y33-ATR2 into W303-1B, and transformants were selected on a tryptophan and uracil minus plate (CM-Trp-Ura). Three colonies were picked for each genotype, and used to inoculate 3 ml of CM-Trp-Ura medium in a 24-well-plate. The recombinant vectors containing ATR2 and P450 (ancXY, ancX, ancX1, ancX2 or ancX3) were directly transformed into W303-1B without extra Y33-ATR2 and cultured in tryptophan minus medium (CM-Trp). The cells were grown at 30 ℃ and 550 rpm for 48 hours, after which the resulting seed cultures were transferred into fresh medium at a ratio of 1:50. The new cultivation was fermented under the same condition for 4 days after feeding 1mM apigenin. For the mutants, flasks containing 30 ml of medium were then inoculated at a ratio of 1:50 using the resulting seed cultures by feeding 1mM apigenin. The main cultures were grown at 30 ℃ and 220 rpm for 4 days. The products extraction method and HPLC detection method was based on our previous study^28^ and was described in detail in the supplementary methods.

## Authorship contribution statement

Qian Wang performed computational analysis and enzyme design, and wrote the manuscript. Xiaonan Liu and Qian Wang designed experiments and interpreted experimental results. Hejian Zhang and Huanyu Chu conducted deep learning work. Chao Shi performed the crystallization of ancX3. Other authors contributed to collating experimental results. Zhenzhan Chang and Jian Cheng revised the paper. Huifeng Jiang conceived and directed the project.

## Declaration of competing interests

The authors declare no competing financial interests.

## Acknowledgements

We thank the staff of beamline BL17U1 at Shanghai Synchrotron Radiation Facility (SSRF), Shanghai, People’s Republic of China, for assistance during data collection.

This project has received funding from the National Key R&D Program of China (Grant No. 2021YFC2103500); National Natural Science Foundation of China (No. 32371499); China Postdoctoral Science Foundation (Grant No. 2019M661032); National Natural Science Foundation of China (NSFC; Grant No. 31901026 and No. 32171418); Tianjin Synthetic Biotechnology Innovation Capacity Improvement Project (No. TSBICIP-KJGG-002-02 and No. TSBICIP-CXRC-015); the Tianjin Science Fund for Distinguished Young Scholars (No.18JCJQJC48300).

## Appendix A. Supplementary data

Supplementary data for this article can be found online.

## Notes

### Competing Interest Statement

The authors have declared no competing interest.

