## Supplementary information for "Cytochrome P450 Enzyme Design by Constraining Catalytic Pocket in Diffusion model"

### Supplementary Methods

#### Chemicals and media

Yeast nitrogen base without amino acids and ammonium sulfate (YNB), Bacto peptone, Bacto yeast extract, Luria Broth (LB), agar, lithium acetate, ssDNA and glucose were obtained from Solarbio, China. Kanamycin, ampicillin, amino acids, adenine, histidine, leucine, tryptophan and uracil were obtained from Sigma-Aldrich, USA. Chromatography-grade methanol and acetonitrile were obtained from EMD chemicals, USA. Chromatography-grade formic acid and isopropanol were obtained from Thermo Fisher Scientific, USA. Authentic reference standard of scutellarein and apigenin was obtained from Solarbio, China. The Hi-Fusion Cloning Mix was purchased from CWBIO, China and used for recombinant plasmid construction. *S. cerevisiae* W303-1B (*MATα leu2-3112 ura3-1 trp1-92 his3-11,15 ade2-1 can1-100*)[1] was used as the parent strain for all engineered strains. Competent cells of *E. coli* DMT (TransGen Biotech, China) were used for recombinant vectors construction. Competent cells of *E. coli* BL21(*DE3*) (TransGen Biotech, China) was used for recombinant protein expression. *E. coli* was grown in LB medium with appropriate antibiotics. Yeast strains were grown in CM medium minus tryptophan or uracil and was used for 24-well plates and shake-flask fermentation of yeast. The corresponding solid plate was added 15 g/L of agar powder.

#### Bacterial expression and purification of the P450 enzyme

Nucleotide sequence of ancX3 was codon optimized for *Escherichia coli* and codons for the N-terminal transmembrane domain was replaced with codons for an optimized sequence of a short hydrophilic peptide “AKKTSSKGK”, and codons for 6 × His-tag were inserted before the stop codon to facilitate purification. The gene was subcloned into $pCW_{ori+}$ expression vector between restriction sites *Nde*I and *Xba*I. The gene of recombinant construct was synthesized by Genscript, China. Expression and purification of recombinant protein were conducted as the methods reported by Gu *et al*[2]. The purity and subunit molecular masses of the recombinant ancX3 was verified by SDS-PAGE analysis, and the protein concentration was determined using a BCA Protein Assay Kit (Pierce, USA) with 2 mg/mL BSA as the standard. Purified enzymes were stored at -80 °C before use.

##### Fluorescence microscopic analysis

The nucleotide sequences of Design6444, Design11361, Design33380, Design49566, Design58683, Design84497, Design91808 and CYP706X1 were fused with green fluorescent protein (GFP) at C-terminal and subcloned between GAL1 promoter and CYC1 terminator of pYES2.0. The plasmids were verified by sequencing and transformed into W303-1B, and transformants were selected on a uracil minus plate (CM-Ura). The cells were grown at 30 °C and 550 rpm for 24 hours, after which the resulting seed cultures were transferred into fresh medium and induced by galactose for 12 hours. The cultivation was diluted twice and used for microscopic analysis. Images were acquired using a Leica DM5000B microscope (Leica, Germany) with a 100 × objective and Leica filter GFP for fluorescent microscopy. Leica LAS AF software (Leica, Germany) was used for image acquisition.

##### HPLC detection

The culture samples were diluted with an equal volume of 100% methanol. After vigorous mixing and ultrasonic breaking for 30 min, the lysates were spun down at 13,000 × g for 10 min. The supernatant compounds were measured at 335 nm using a Kinetex 5 μm Biphenyl 100 Å LC Column (250 × 4.6 mm; Phenomenex, USA) operating at 30 °C. The mobile phase consisted of 0.1% formic acid and acetonitrile with methanol at a flow rate of 1 mL·min-1 using the following gradients: 0-20 min, 22% acetonitrile, 5% methanol; 20 min to 22 min, acetonitrile increased from 22% to 90%; 25 min, 22% acetonitrile. Subsequently, the column was washed and equilibrated for 5 min before the next injection. 30 μL of the sample were injected into the HPLC system and each run was stopped at 30 min after the injection.

### Supplementary Figures

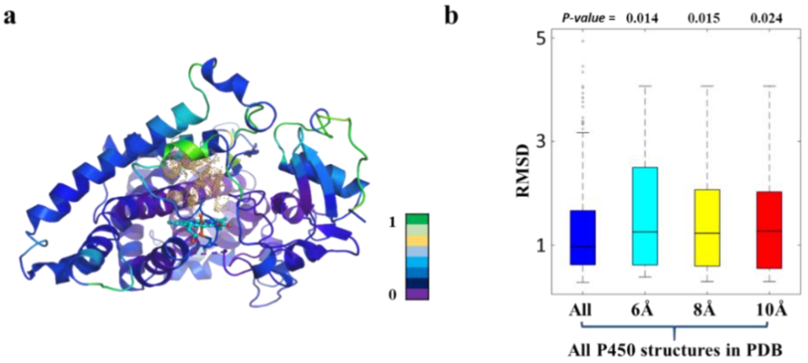


**Figure S1. Structural variability of the catalytic pocket between different P450s in PDB database. (a)** The structural alignment of non-redundant P450 structures in the PDB database reveals varying levels of structural variability among different regions. The color gradient in the legend, from green to purple, signifies decreasing structural variability, with the green region corresponding to the binding pocket area exhibiting the highest variability. **(b)** A box plot illustrates the structural variability (RMSD) at different distances (all, 6Å, 8Å, and 10Å) from the active site. The region within a 6Å distance from the active site displays the most significant variability.

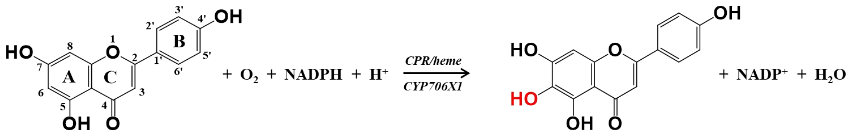


Apigenin Scutellarein

**Figure S2. The CYP706X1 in E. breviscapus catalyzes the 6-hydroxylation of apigenin to yield scutellarein.** The figure depicts the reaction scheme catalyzed by CYP706X1 in conjunction with CPR (cytochrome P450 reductase) and heme.

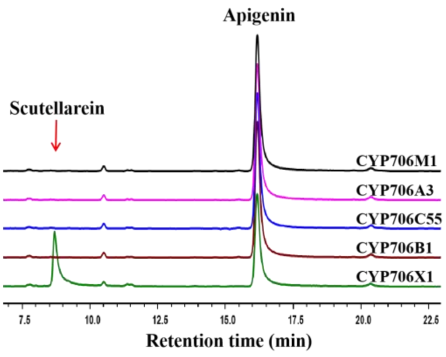


**Figure S3. HPLC analysis of the fermented products catalyzed by various cytochrome P450 enzymes (CYP706X1: green, CYP706B1: brown, CYP706C55: blue, CYP706A3: purple, CYP706M1: black).** Among these enzymes, only CYP706X1 exhibits reactivity towards the substrate apigenin, leading to the formation of the product scutellarein. The substrate apigenin and the product scutellarein are clearly labeled in the figure.

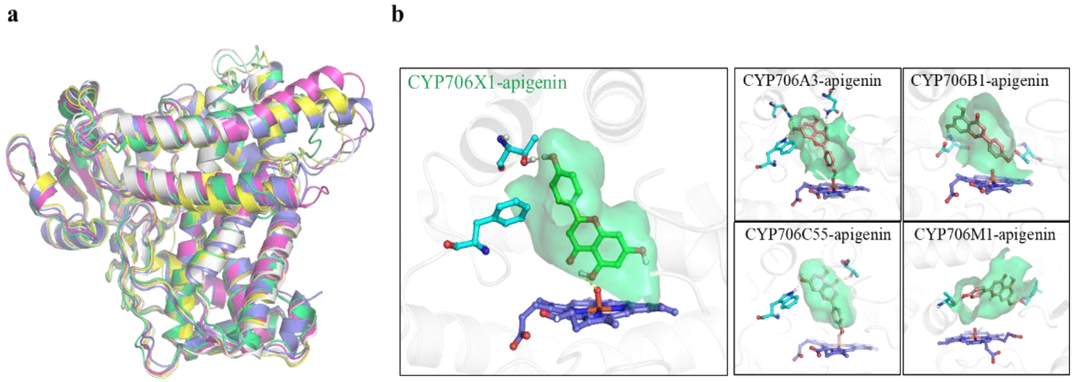


**Figure S4. The structural alignment and apigenin-binding models of CYP706X1, CYP706B1, CYP706C55, CYP706A3 and CYP706M1. (a)** The structural alignment of the five P450 enzymes reveals a unique structural arrangement, with each enzyme represented in a different color. **(b)** The structural models depict the binding of apigenin within these five P450 enzymes. The substrate apigenin and critical residues are illustrated as ball-and-stick models, while the green regions delineate the shapes of the substrate-binding domains. Dashed lines signify hydrogen bond interactions. All residues in the figures are colored cyan, with the apigenin in CYP706X1 shown in green, and brown in the other four proteins.

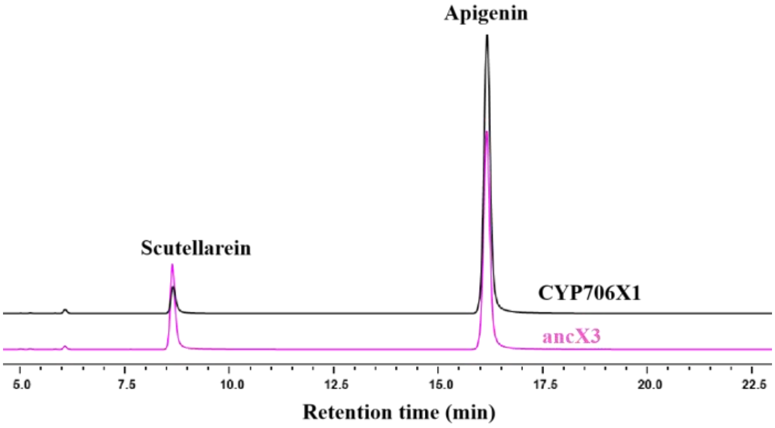


**Figure S5. HPLC analysis of the fermented products catalyzed by CYP706X1 (black) and ancX3 (magenta).** CYP706X1 and ancX3 exhibit reactivity towards the substrate apigenin, leading to the formation of the product scutellarein. The ancX3 shows a higher level of catalytic capability compared to CYP706X1. The substrate apigenin and the product scutellarein are labeled in the figure.

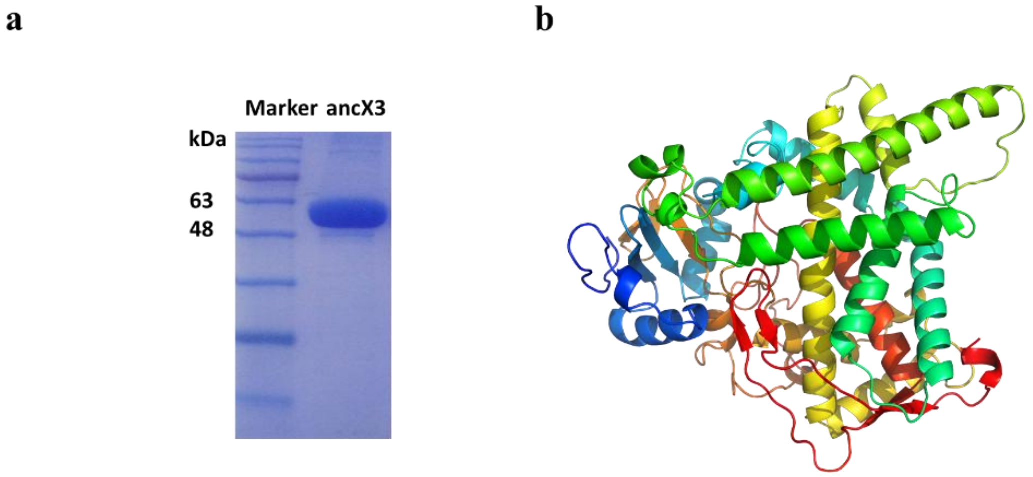


**Figure S6. SDS-PAGE analysis of the purified ancX3 and the crystal structure of ancX3. (a)** SDS-PAGE analysis of the purified ancX3 and showed in the second gel lane. The molecular weight of protein ancX3 is 53.90 kDa. **(b)** The structural representation of the ancX3 crystal structure is shown as a cartoon model and the crystal resolution is 2.3Å.

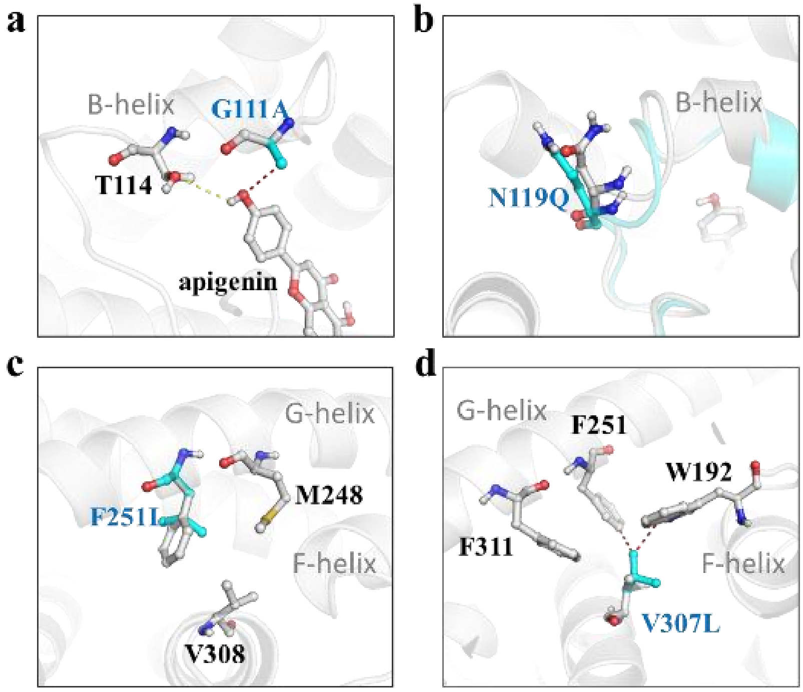


**Figure S7. Structural analysis of four mutations (i.e., G111A, N119Q, F251L and V307L) in ancXY-16. (a)** The G111A mutation introduced a collision between the side chain of alanine and substrate apigenin thus impairing the activity of ancXY-16. **(b)** The N119Q was far from the active center, and the mutation might influence the hydrophilicity of the protein surface. **(c and d)** The F251L and V307L changed the local hydrophobic property of ancXY-16. Ball-and-stick models are used to represent the substrates and residues. Wild-type residues are colored white and labeled in black, while mutations are colored cyan and labeled in blue.

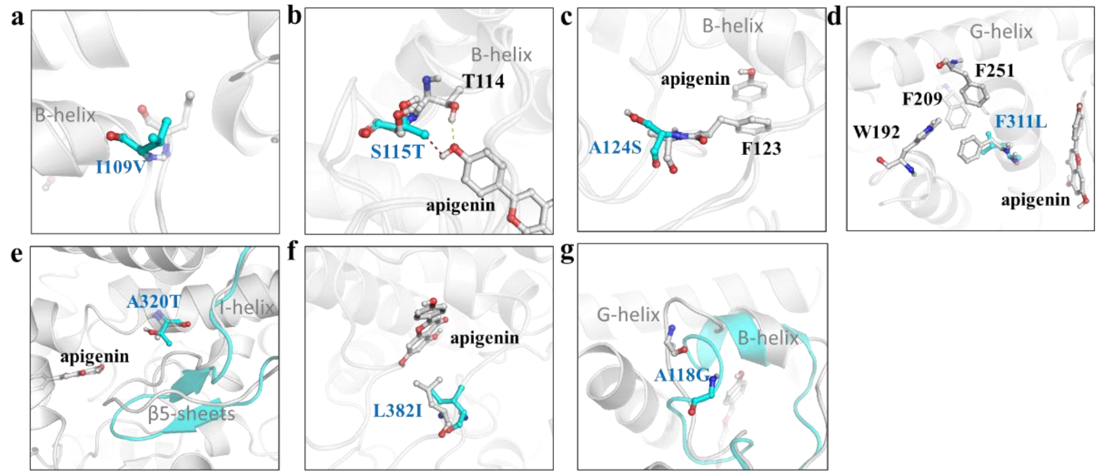


**Figure S8. Structural analysis of seven mutations (i.e., I109V, S115T, A124S, F311L, A320T, L382I and A118G) in ancXY-12. (a, d and f)** The I09V, F311L and L382I changed the local hydrophobic properties of ancXY-12 to influence the catalytic activity. **(b)** The S115T introduced space collision between the side chain of threonine and substrate apigenin, which impaired the activity of ancXY-12 severely. **(c and e)** The A124S and A320T are far from substrate apigenin, and may influence the hydrophilic environment of ancXY-12. **(g)** The A118G affected the main-chain conformation of the B-helix by introducing a turn in front of the B-helix. Ball-and-stick models are used to represent the substrates and residues. Wild-type residues are colored white and labeled in black, while mutations are colored cyan and labeled in blue.

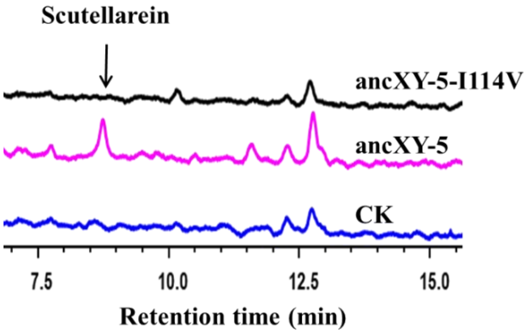


**Figure S9. HPLC analysis of the fermented products catalyzed by ancXY-5 (magenta) and ancXY-5-T114V mutation (black).** The T114V mutation deactivated the catalytic activity of ancXY-5. The control check is colored blue and the scutellarein is indicated with a black arrow.

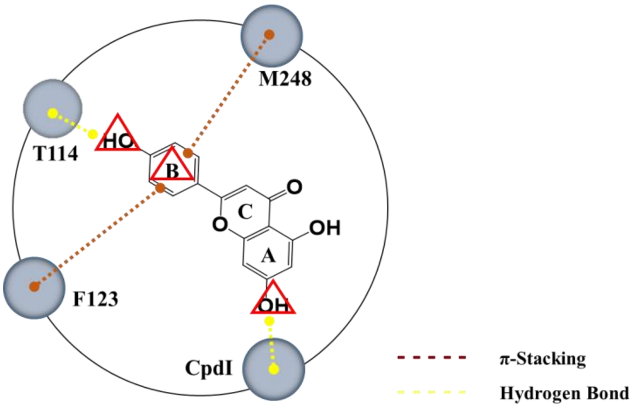


**Figure S10. The representative diagram of "three-point fixation" model.** Three pivots in apigenin are marked with red triangles, while yellow and brown dash lines represent hydrogen bond and π-stacking interactions, respectively. Key residues and CpdI in this model are represented by gray circles.

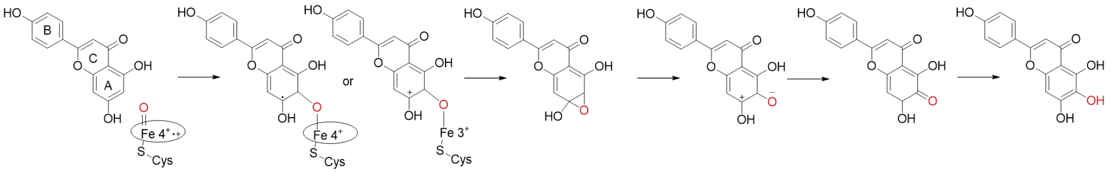


apigenin

scutellarein

**Figure S11. Proposed possible flavone 6-hydroxylation reaction mechanism.** The oxyferryl species of CpdI attacks the π-system of the "A" ring of apigenin to produce cationic σ-complex or radical σ-complex and then occurs ring closure to produce epoxide. The epoxide undergoes a hydride shift and enol isomerization to gain scutellarein. The mechanism speculation is based on a previous study[3].

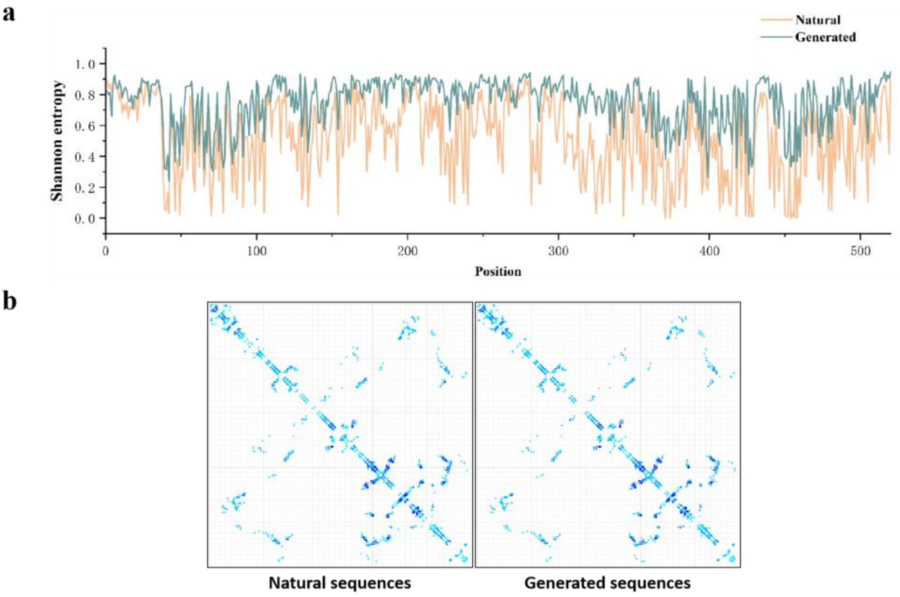


**Figure S12. The evaluation of P450Diffusion fine-tuning model generated sequences. (a)** The amino acids variations of generated sequences (orange) and natural sequences (green) at each position are expressed as Shannon entropy. Low Shannon entropy values represent highly conserved positions and high entropy values indicates high amino-acid diversity at a given position. **(b)** The coevolution feature of natural sequences (left) and generated sequences (right). The coevolution analysis was performed using GREMLIN software4, and the multiple sequence alignments (MSAs) for natural sequences and generated sequences were used as input files, respectively.

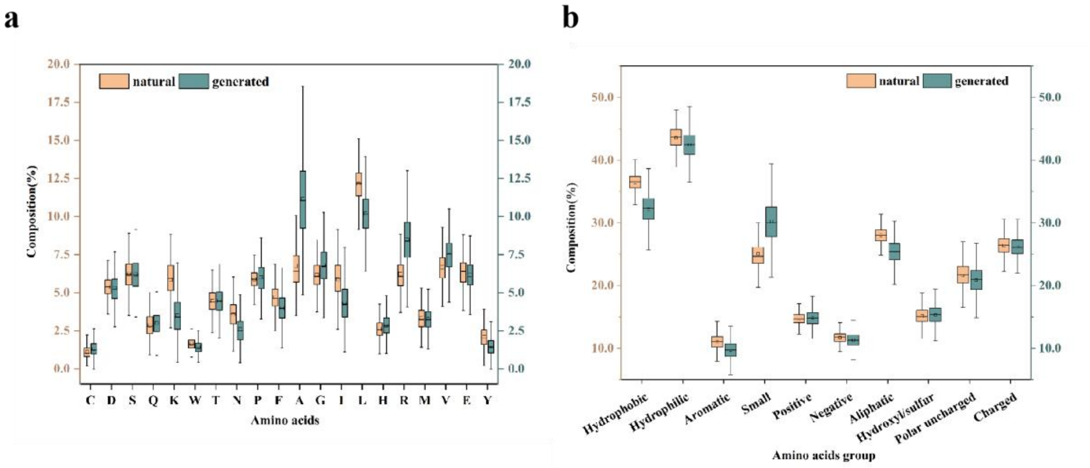


**Figure S13. The amino acid composition of raw sequences and generated sequences. (a)** A boxplot displays the percental amino acid composition of raw sequences, grouped by their physicochemical properties. **(b)** The amino acid distribution and compositional variability of sequences generated by P450Diffusion are highly similar to those of the natural sequences. P450Diffusion is able to replicate the specific physicochemical properties found in natural sequences.

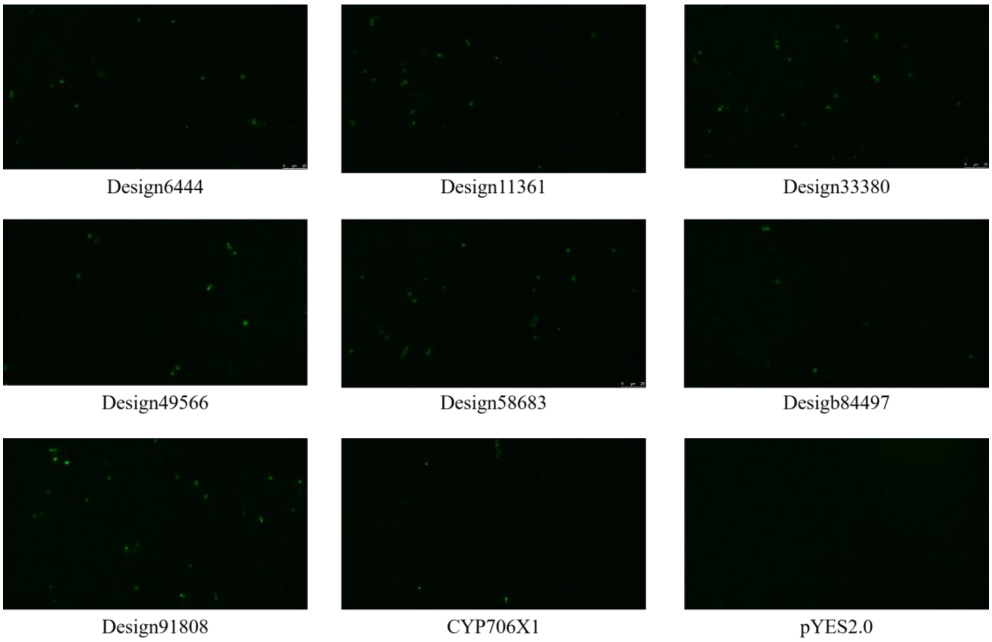


**Figure S14. Fluorescence microscopic analysis of inactive designs generated by deep learning.** Design6444, Design11361, Design33380, Design49566, Design58683, Design84497 and Design91808 were fused with GFP and then visualized by fluorescence microscopy. The pYES2.0-CYP706X1(EbF6H) was used as positive control. The pYES2.0 was used as negative control. The green dots represent correctly expressed and folded P450s tagged with green fluorescent proteins.

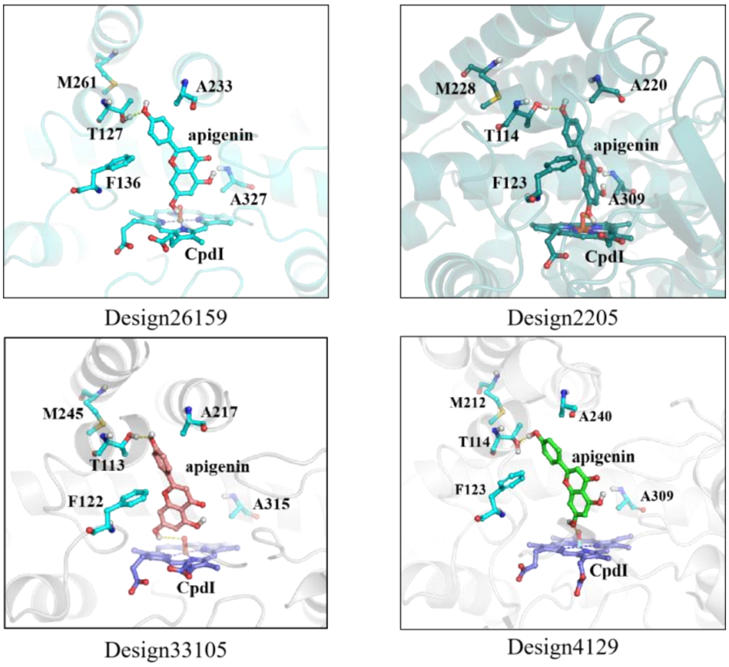


**Figure S15. The apigenin binding model of active designs.** Substrates in active designs bind to catalytic pockets in a manner highly similar to natural CYP706X1. Design26159, Design2205, Design33105 and Design4129 were selected as representative. The carbon of four designs is colored cyan, marine, brown and green, respectively. Key residues were labeled in figures.

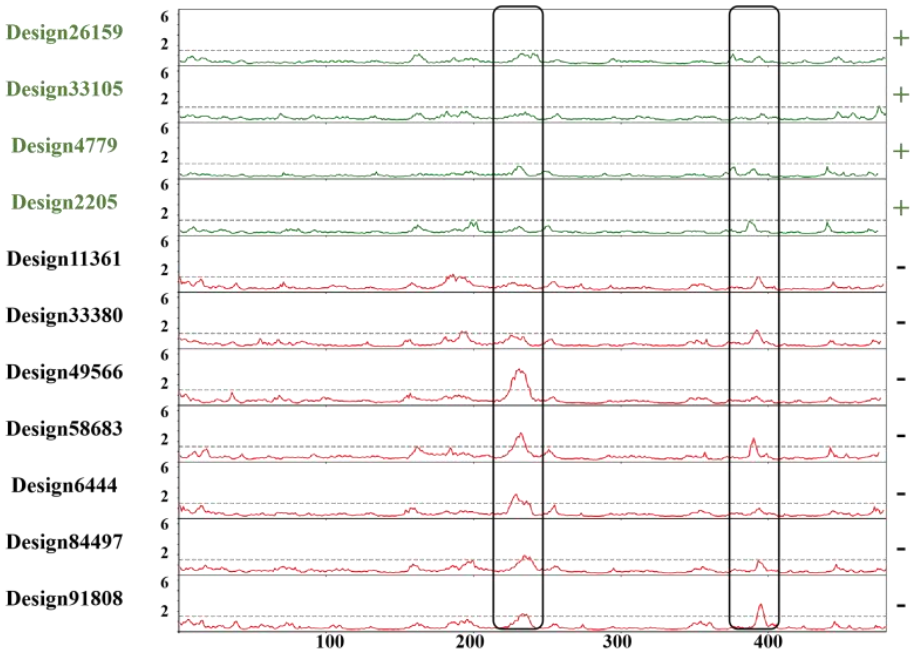


**Figure S16. Root Mean Square Fluctuation (RMSF) curve of the designed P450s (N-terminal trimmed) in molecular simulations.** This curve illustrates the average fluctuations of each residue within the designed P450s, reflecting their flexibility and stability during the simulation. Higher peaks typically correspond to flexible regions, while lower peaks indicate more stable regions. Four active designs, including Design26159, Design33105, Design4779, and Design2205 were selected as active controls and highlighted in green. Additionally, seven inactive designs, consisting of Design11361, Design33380, Design49566, Design58683, Design6444, Design84497 and Design91808, have been marked in red.

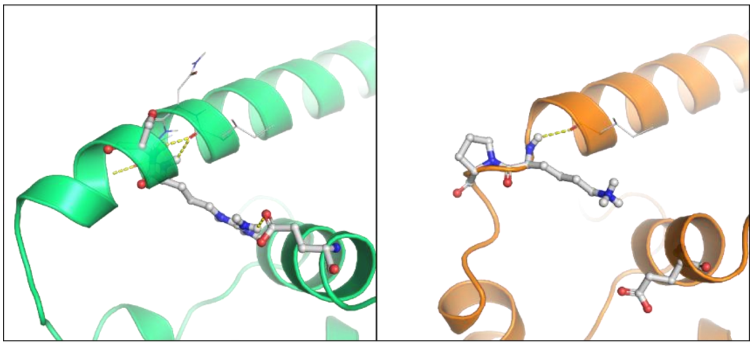

**Design33105**
**R227_S228_E254**

**Design33380**
**K229_P230_E251**

**Figure S17. The local fluctuations in region 220-230 (N-terminal trimmed) of active Design33105 (green) compared to inactive Design33380 (brown).** The mutations were represented as stick-and-ball model at both designs. The salt-bridge between R227 and E254 in active Design33105 are broken in inactive Design33380.

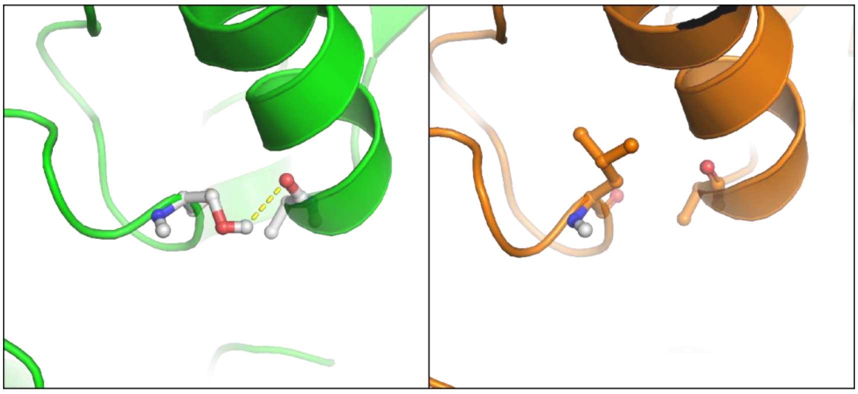

**Design33105** **Design91808**
**S407-A51** **L407-A51**

**Figure S18. The local fluctuations in region 390-410 (N-terminal trimmed) of active Design33105 (green) compared to inactive Design91808 (brown).** The mutations were represented as stick-and-ball model at both designs. The hydrogen bond between S407 and A51 in active Design33105 are broken in inactive Design91808.

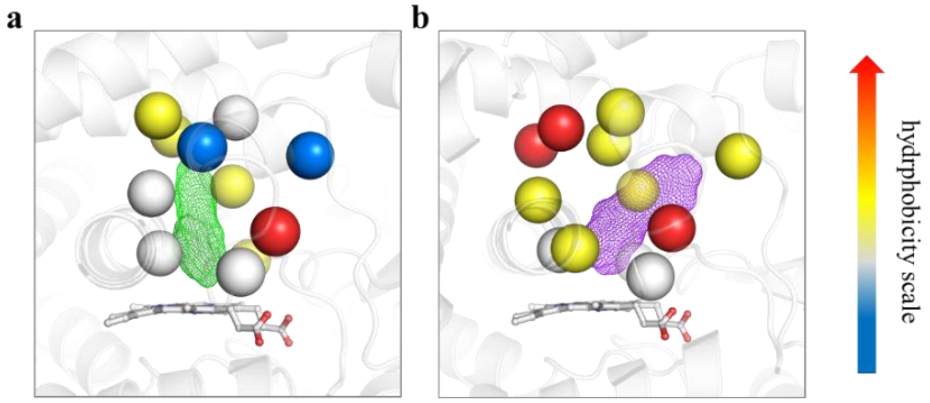


CYP706X1 SbaiCYP82D4

**Figure S19. Comparative analyses of the substrate-binding modes between CYP706X1 and SbaiCYP82D4.** A "vertical binding mode" and an "oblique binding mode" were found in CYP706X1 **(a)** and SbaiCYP82D4 **(b)**, respectively. Residues in the catalytic pockets and heme molecules are represented as spheres and ball-and-sticks, respectively. The gradient from blue to red represents increasing hydrophobicity. The substrates in CYP706X1 and SbaiCYP82D4 are represented as mesh, and colored green and purple, respectively. The cartoon representations of both proteins were colored white.

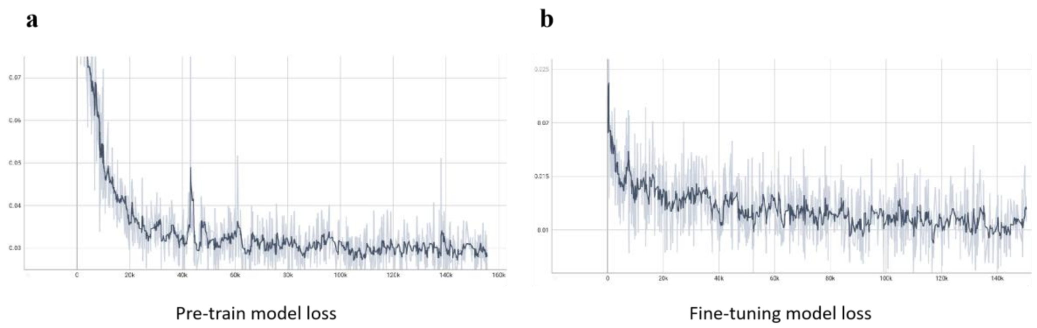


**Figure S20. The losses of the P450diffusion Pre-trained model (a) and Fine-tuning model (b) in the training procedure.** The loss of the training process is smoothed in the figure.

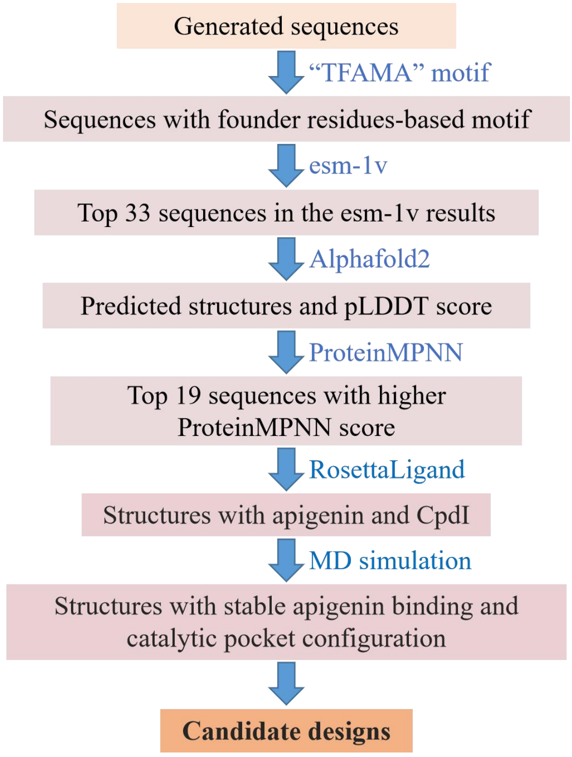


**Figure S21. The computational evaluation and structure-based virtual screening for generated sequences.** The virtual screening protocol is shown as a flowchart.

### Supplementary Tables

**Table S1. Data collection and refinement statistics of ancX3**

| **Data collection** | |
|---|---|
| Beamline | SSRF17U1 |
| Integration Package | XDS |
| Wavelength (Å) | 0.97918 |
| Space group | I432 |
| Unit cell: a, b, c (Å), α, β, γ (°) | 197.61 197.61 197.61 90.00 90.00 90.00 |
| Resolution (Å) | 46.58-2.30 (2.38-2.30) |
| Rmerge | 0.134 (3.349) |
| Rmean | 0.136 (3.374) |
| Rpim | 0.015 (0.406) |
| Mean((I)/sd(I)) | 30.8 (2.0) |
| CC(1/2) | 1.000 (0.705) |
| Completeness (%) | 100 (100) |
| Total number of measured reflections | 2283864 |
| Total number of unique reflections | 29446 |
| Multiplicity | 77.6 (68.6) |
| Mean(Chi^2) | 1 (0.96) |
| Molecules in an asymmetric unit | 1 |

*Numbers in parentheses are for the highest resolution shell.

**Table S2. Sequence similarity of 17 designs compared to CYP706X1.**

| **Design** | **Sequence similarity** | **Activity** |
|---|---|---|
| Design33380 | 83.8 | No |
| Design91808 | 83.6 | No |
| Design84497 | 84.3 | No |
| Design49566 | 86.2 | No |
| Design42565 | 80.8 | Yes |
| Design11361 | 77.5 | No |
| Design6444 | 77.5 | No |
| Design58683 | 76.9 | No |
| Design4779 | 77.1 | Yes |
| Design3644 | 77.8 | Yes |
| Design4129 | 76.7 | Yes |
| Design58 | 77.3 | Yes |

| | | |
|---|---|---|
| Design26159 | 71.3 | Yes |
| Design2205 | 75.9 | Yes |
| Design33105 | 75.9 | Yes |
| Design468 | 75.5 | Yes |
| Design12854 | 72.8 | Yes |

**Table S3 and Table S4 could be found in Supplementary Excel file.**

### The protein sequences and DNA sequences used in this study

#### Protein sequences used in this study

##### Natural sequences

**>CYP706X1**

MASNELAFSALLVTLVLVLISWYKREISNSRKAGTPPLPPGPKGLPLVGSLPFLGPNIHQELTKISHQYGPIFKLYLGSKLHIVVNSAELAKVITAEQDESF
ANRAPHIAGLATSYGGNDIAFAPNNANRRNLRKVLVQEVLSNVNLEASHAYRRHEVRKAVKYVYDRVGMDVDINEISFSTVLNVFTNIIWRKGFVDD
GANYANLSENIQKVICRIVEIAEGLNISDFFPMLARFDLQGVERKMKDQMKQFDKIIEPTIKERMNSRSTNVEETVEHKGRKDFLQILLEHTDQKNGTS
ITMTQLKALVADIFLGGTDATSAMVEWAMTEIFRDQKVMKRVQDELEEIVGLNNIVEESHIPKLKYLEAVCKETFRLHPPIPFLLPRAPIKSCTVGGYTV
PEGATIFVNVWAIQRDPRHWENPSEFNPDRFLNRNGSTEKWDYSGTNLTFLPFGSGRRRCPGIPLGEKMMMHILASLMHSFDWKLPNGEELDLSERF
GIALKKKKPLVAVPTKRLSDLSLYM*

**>Cnan706X**

MAASSHSSLWEEATNNKHEVALAVFVGTLVILAISWYKRSNSRTGKGTPPLPPGPKGLPIVGYLPFLSPNLHHEFAKIANQYGPIFKLYLGSKLHIVVNS
ADLAKVVTGEQDESFANRDPHIAGLTASYGASDVAWQNNNSNRRNLRKVLVHEVLSNKNLEASHAYRRSEVRKTIKNVHDMIGTAVDINEVSFSTVL
NILTHIVWGNSFVEGAKYPNLAADIRKVVLDIVEIAEGLNLSDFFPMLARFDFQGVERRVKAQVKKFDHIFETTIEERTNSKSKVSEEAVKQEGRKDFL
QILLELLDQNTATSINMTQLKALVVDIFLGGTDATAAMTEWAMAEILRNPKVMKKVQDELAEVVGLNNIVEESHLPKLKYLDAVFKETFRLHTPLPFL
LPRTPDKSCVVGGYTVPKGATVFLNVWAIQRDPQNWENPSEFNPERFLNNKGSEKWDYSGTNSTYFPFGSGRRRCPGILLGEKMMMHILASLMHSF
DWSLPKGEELDLSDKFGIAMKKKMPLVVIPSLRLSDLSLYS*

**>Lsal706X**

MASNELAFSALLVTLVLVLISWYKREISNSRKAGTPPLPPGPYGLPLVGYLPFLGPSLHHELTKMAHRYGPIFKLYLGSKLHIVVNSADLAKVITSEQDE
SFANRAPHIAGLATSYGANDIAFADNNANRRNLRKILVHEILSNVNLEASHAYRRREVRKTIKSVHDMIGMPVDINEMSFSTVVNVLTSIVWGNSMVE
GTKHSNLGEEIRKVVSEIVDIAEGLNISDFFPKLARFDLQGVEQKMKRKMKQFDWIFETTIEERINLKSTHGEDALKHEGRKDFLQILLELKDKKSITM
TQLKALVVDIFLGGTDATSAMVEWAMAEILKNQKVMKKVQDELAEIVGLKNMVEESHLPKLKYLNATFKETFRLHTPLPVLLPRTPSKSCMVGGYLI
PRDSTVFLNVWAIQRDPQHWENPSEFNPERFLNYEGSGKWDYSGTNSKYFPFGSGRRRCPGIPLAEKMMLHILASLLHSFDWSLPKGEDHDLFEKFGI
ALKKKKPLVAVPSPRLIDLSLYM*

##### Ancestral protein sequences

**>ancXY**

MASNELAFSALLVTLVLVLISWYKREISNSRKAGTPPLPPGPRGLPVVGYLPFLGPNLHQEFTKMAHRYGPIFKLHLGSKLHIVVNSADLAKVVAREQ
DETFANRNPPVAALAITYGGQDIAWSNNNSNWRNLRKVLVHEVLSNKNLEASRSFRRREVRKTIKNVYEKIGTEIDINEIAFSTELNVLTSMVWGKSL
VEGEKYSNLGDEFREVVSKIVEILGAPNISDFFPILAWFDLQGVEREMKRQLKQLDRIFESIIEERINSNSTKSEEAVEHEGRKDFLQILLELKDQKDATSI
NITQIKALLVDILLGGTDTTSTMVEWAMAEILQNQKVMKKVQDELAEIVGLNNIVEESHLPKLKYLDAVIKETFRLHPPLPLLIPRSPNQSCTVGGYTIP
KGSTVFLNVWAIHRDPQYWDNPLEFNPERFLNREGTDKWDYNGNNLKFLPFGSGRRRCPGIPLGEKMLMYILASLLHSFDWSLPKGEEHDLSDKFGI
ALKKRKPLIAIPSQRLPDASLYM*

**>ancX**

MASNELAFSALLVTLVLVLISWYKREISNSRKAGTPPLPPGPRGLPLVGYLPFLGPNLHQELTKMAHRYGPIFKLYLGSKLHIVVNSADLAKVVTGEQD
ESFANRAPHIAGLATSYGANDIAFADNNANRRNLRKVLVHEVLSNVNLEASHAYRRREVRKTIKNVHEMIGNEVDINEIAFSTVLNVLTSIVWGKSMV
EGAKYSNLGEEIRKVVSGIVEIAGGLNISDFFPMLARFDLQGVERKMKRQMKQFDKIFESTIEERINSKSTNVEEAVKHEGRKDFLQILLELKDQKNET
SITMTQLKALVVDIFLGGTDATSAMVEWAMTEILRNQKVMKKVQDELAEIVGLNNIVEESHLPKLKYLDAVFKETFRLHPPLPFLLPRAPNKSCTVGG
YTIPKGSTIFLNVWAIQRDPQYWENPSEFNPERFLNYEGSEKWDYSGTNSKFFPFGSGRRRCPGIPLGEKMMMHILASLLHSFDWSLPKGEEHDLSDK
FGIALKKRKPLIAVPSPRLSDASLYM*

**>ancX1**

MASNELAFSALLVTLVLVLISWYKREISNSRKAGTPPLPPGPKGLPLVGSLPFLGPNIHQELTKISHQYGPIFKLYLGSKLHIVVNSAELAKVITAEQDESF
ANRAPHIAGLATSYGGNDIAFAPNNANRRNLRKVLVQEVLSNVNLEASHAYRRHEVRKAVKYVYDRVGMDVDINEISFSTVLNVLTNIIWRKGFVDD
GANYANLSEKIQKVICRIVEIAEGLNISDFFPMLARFDLQGVERKMKDQMKQFDKIIETTIKERMNSKSTNVEETVEHKGRKDFLQILLEHTDQKNGTS
ITMTQLKALVADIFLGGTDATSAMVEWAMTEIFRDQKVMKRVQDELEEIVGLNNIVEESHIPKLKYLEAVCKETFRLHPPIPFLLPRAPNKSCTVGGYT
VPEGATIFVNVWAIQRDPRHWENPSEFNPDRFLNCNGSTEKWDYSGTNLTFLPFGSGRRRCPGIPLGEKMMMHILASLMHSFDWKLPNGEELDLSER
FGIALKKKKPLVAVPTKRLSDLSLYM*

**>ancX2**

MASNELAFSALLVTLVLVLISWYKREISNSRKAGTPPLPPGPRGLPLVGYLPFLGPNLHQELTKIAHRYGPIFKLYLGSKLHIVVNSADLAKVITGEQDE
SFANRAPHIAGLATSYGANDIAFADNNANRRNLRKVLVHEVLSNVNLEASHAYRRREVRKTIKNVHDMIGMEVDINEISFSTVLNVLTSIVWGKGMV
EGAKYSNLGEEIRKVVSGIVEIAEGLNISDFFPMLARFDLQGVERKMKRQMKQFDKIFETTIEERINSKSTNVEEAVKHEGRKDFLQILLELKDQKNGT
SITMTQLKALVVDIFLGGTDATSAMVEWAMTEILRNQKVMKKVQDELAEIVGLNNIVEESHLPKLKYLDAVFKETFRLHPPLPFLLPRTPNKSCTVGG
YTIPKGSTIFLNVWAIQRDPQHWENPSEFNPERFLNYEGSEKWDYSGTNSKFFPFGSGRRRCPGIPLGEKMMMHILASLLHSFDWSLPKGEEHDLSEK
FGIALKKKKPLVAVPSPRLSDLSLYM*

**>ancX3**

MASNELAFSALLVTLVLVLISWYKREISNSRKAGTPPLPPGPRGLPLVGYLPFLGPNLHQELTKMAHRYGPIFKLYLGSKLHIVVNSADLAKVVTGEQD
ESFANRAPHIAGLATSYNASDIAFADNNANRRKLRKVLVHEVLSNVNLEASHAYRRREVRKTIKNVHEIIGNEVDINEIAFSTVLSVLTSIVWGKSMVK
GAKYSNLVAEMRKFVSGVVEIAGELNISDFFPMLARFDFQGVERRMKKQMKLFDKIFESTVEERINSRSAIKEEAVKEEGRKDFLQILLELQEQNNETS
ITMTQMKALVVDIFLGGTDATSAMIEWAMTEILRNRQVMKKVQDELAEIVGLNNIVEESHLPKLKYLDAVFKETFRLHPPLPFLLPRAPNKSCTVGGY
TIPKGSTIFLNVWAIQRDPQYWENPSEFNPERFLNYKGSEKWDYAGTNSKFFPLGSGRRRCPGVSLGEKMMMHILASLLHSFDWSLPTGQKLDLSDK
FGIALKKRKPLIAVPSPRLNDASLYM*

**>ancX-16**

MASNELAFSALLVTLVLVLISWYKREISNSRKAGTPPLPPGPRGLPVVGYLPFLGPNLHQEFTKMAHRYGPIFKLHLGSKLHIVVNSADLAKVVAREQ
DETFANRNPPIAGLATSYGANDIAFANNNSNWRNLRKVLVHEVLSNKNLEASRSFRRREVRKTIKNVYEKIGTEIDINEIAFSTELNVLTSMVWGKSLV
EGEKYSNLGDEFREVVSKIVEIAGAPNISDFFPILAWFDLQGVEREMKRQMKQFDRIFESIIEERINSNSTKSEEAVEHEGRKDFLQILLELKDQKDATSI
NITQIKALVVDIFLGGTDATSAMVEWAMAEILQNQKVMKKVQDELAEIVGLNNIVEESHLPKLKYLDAVIKETFRLHPPLPLLLPRSPNQSCTVGGYTI
PKGSTVFLNVWAIHRDPQYWDNPLEFNPERFLNREGTDKWDYNGNNLKFLPFGSGRRRCPGIPLGEKMLMYILASLLHSFDWSLPKGEEHDLSDKF
GIALKKRKPLIAIPSQRLPDASLYM*

#### Other subfamily protein sequences

>CYP73A1  (NCBI Number: Q04468)

MDLLLIEKTLVALFAAIIGAILISKLRGKKFKLPPGPIPVPIFGNWLQVGDDLNHRNLTDLAKRFGEILLLRMGQRNLVVVSSPELAKEVLHTQGVEFGS
RTRNVVFDIFTGKGQDMVFTVYGEHWRKMRRIMTVPFFTNKVVQQYRYGWEAEAAAVVDDVKKNPAAATEGIVIRRRLQLMMYNNMFRIMFDRR
FESEDDPLFLKLKALNGERSRLAQSFEYNYGDFIPILRPFLRNYLKLCKEVKDKRIQLFKDYFVDERKKIGSTKKMDNNQLKCAIDHILEAKEKGEINE
DNVLYIVENINVAAIETTLWSIEWGIAELVNHPEIQAKLRHELDTKLGPGVQITEPDVQNLPYLQAVVKETLRLRMAIPLLVPHMNLHDAKLGGFDIPA
ESKILVNAWWLANNPDQWKKPEEFRPERFLEEEAKVEANGNDFRYLPFGVGRRSCPGIILALPILGITIGRLVQNFELLPPPGQSKIDTDEKGGQFSLHI
LKHSTIVAKPRSF*

>CYP706A3  (NCBI Number: KAG7604930)

MTDISSLFRNRSRKDQLDYGLTVIVISTLCWCLWLYAKCKRRSPPLPPGPWGLPIIGNLPFLQPELHTYFQGLAKKHGPIFKLWLGAKLTIVVTSSEVAQ

EILKTNDIIFANHDVPAVGLVNTYGGTEIIWSPYGPKWRMLRKLCVNRILRNAMLDSSTDLRRRETRQTVRYLADQARVGSPVNLGEQIFLMMLNVV
TQMLWGTTVKEEEREVVGAEFLEVIREMNDLLLVPNISDFFPVLSRFDLQGLAKRMRRPAQRMDQMFDRIINQRLGMDRDSSDGRAVDFLDVLLKV
KDEEAEKTKLTMNDVKAVLMDMVLGGTDTSLHVIEFAMAELLHNPDIMKRAQQEVDKVVGKEKVVEESHISKLPYILAIMKETLRLHTVAPLLVPR
RPSQTTVVGGFTIPKDSKIFINAWAIHRNPNVWENPLKFDPDRFLDMSYDFKGNDFNYLPFGSGRRICVGMAMGERVVLYNLATFLHSFDWKIPQGER
VEVEEKFGIVLELKNPLVATPVLRLSDPNLYL*

>CYP706B1  (NCBI Number: XP_016705189)

MLQIAFSSYSWLLTASHQKDGMLFPVALSFLVAILGISLWHVWTIRKPKKDIAPLPPGPRGLPIVGYLPYLGTDNLHLVFTDLAAAYGPIYKLWLGNKL
CVVISSAPLAKEVVRDNDITFSERDPPVCAKIITFGLNDIVFDSYSSPDWRMKRKVLVREMLSHSSIKACYGLRREQVLKGVQNVAQSAGKPIDFGETA
FLTSINAMMSMLWGGKQGGEQKGADVWGQFRDLITELMVILGKPNVSDIFPVLARFDIQGLEKEMTKIVNSFDKLFNSMIEERENFSNKLSKEDGNT
EAKDFLQLLLELKQKNDSGISITMNQVKALLMDIVVGGTDTTSTMMEWTMAELIANPEAMKKVKQEIDDVVGSDAAVDETHLPKLRYLDAAVKET
FRLHPPMPLLVPRCPGDLSNVGGYSVPKGTRVFLNIWCIQRDPQLWENPLEFKPERFLTDHQKLDYLGNDSRYMPFGSGRRMCAGVSLGEKMLYSSL
AAMIHAYDWNLADGEENDLIGLFGIIMKKKKPLILVPTPRPSNLQHYMK*

>CYP706C55 (NCBI Number: AYN73068)

MSPSISLSLLHNLESSLSSNDFSLLSLLFLLSGALAIGFWAWRRSAEKKNSLPLPPGPAGLPLVGNLPFLDPELHTYFATLAMTYGPILKLQLGKKLGIVV
TSPATAREVLKDNDVTFANRDVPIAGRVAFYGGSDIVWNSYGPEWRMFRKVCGLKMLSNHALDSVYELRRREVRRTVGYFLLQAGSPVNVGEQMFL
TVLNVITSMLWGGTAQGEEKESLGADFRQAVSSLTKLLGKPNISDFYPSLARFDLQGIERQMKGLAKRFDGIFQKMIEQRLKMQRENGSESLDGGEEN
KDFLQFLLNLKDEEDAQTQLTTTGLKALLMDMLVGGTDSSSNTIEFAMAEIINKPKVLQNIQQELETVVGRGNVVEESHIPKLPYLQAVMKESLRLHP
PVPLLIPHCPSATCTVGGYTVPKGSRVFINVWAIHRDPLIWRDPLEFDPERFLHSEGNYNEHNFNYFPFGSGRRMCVGILMAERMVLYSLATLLHSFDW
KLPKGEKMDLTEQFGIVMKKKKPLMAVPSPRFSNSRLYE*

>CYP706M1 (NCBI Number: AGJ03150)

MDMSTIWYYWVSIILGVFIFLIVGIQKWRSKKLPPGPFALPLLGHLHLLEPNVHECLSKISEKFGPLMSFKFGMKTSIIVSSPAMAKEILRENDQIFANRS
IPVVARCIAYDASDILWSPNGPRWRLLRKICVKELFSPKSTEALQPLRREEVRRTMGNIYKDSINGVSVDVGAKAFITSLNLITNMMWSTSTETGERGG
EFKDLVGELVHVLGVPNASDLFPFLERFDVQGLYRRMEKVFVRFDKMFDGIIEDKLSGKSKEKDFLQSLLDLVERGVDEQDPDSVQLTMKDVKVLL
MDMVTGSTDTTSNTVEWAMAELLQQPEIMKRAQKELEEVVGLDNMVEECHLSQLPYLDIIVKEVLRLHPALPLLAPHRPERECEIGGYIIPKDTQVLI
NVWSIQRNPKVWKEPLLFDPERFSDSKWDYNGRDFDYFPFGSGRRICAGLSMAKIMVHYSLASLLHSFDWSLPVAEKLNMDEKYGIVLRKAVPLVAL
PKPRLLYPNLYE*

#### Deep learning protein sequences

**>Design11361**

MASNELAFSALLVTLVLVLISWYKREISNSRKAGTPPLPPGPKGLPLVGFLPFLGPNLHLDLLKMANNYGPIFKLYLGSNLHIVVNSADLAKVVTGEQD
ESFANRAQHIAGLATSYNASDIAFADNNANRRKLRKVLVHEVLSNVNLEASNAYRRREVRKTIKNVHEIIGNEVDINEIAFSTVLSVLTSIVFGKSMVK
GAKYSNLVADMRKFVSGVVEIAGGLNISDFFPVLARFDFQGVKRKMADQMKMFDKIFETSVEERINSRSAIIEETVKQEGRKDFLQILLELLDQNTET
SITMTQLKALVVDIFLGGTDATSAMVEWAMTEIFRDKKVMKRVQDELAEVVGLNNIVEESHLPKLKYLDAVFKETFRLHPPLPFLLPRAPNKSCTVG
GYTVPKGSTIFLNVWAIQRDPQYWENPSDFNPERFLNYKGSNKWDYAGTNLKFFPFGSGRRRCPGVSLGEKMLMHILASLLHSFDWSLPTGQKLDLS
DKFGITLKKRKPLIAVPSPRLSDASLYL*

**>Design33380**

MASNELAFSALLVTLVLVLISWYKREISNSRKAGTPPLPPSPKSLPIVGHLPFLGTDIHHELTEISHQYGPIFKFHLGSKLHIIINSAELAKVITVEQDESFA
NRWPHIAGIATSYGGNDIAFAPNNANWRNLRKVLVQEVLSNVNLEASHAYRRHEVRKAVKYVYDRVGMDVDINEISFSTVLNVFSNIIWRKGFVDDG
TNYANLSEKIQKVICRIVEIAEGLNISDFFPMLARFDLQGVERKMKTQMKQFDKIFETTVDERINSKPAISEEAVKEEGRKDFLQILLELLDQNTATSITM
TQMKALVVDVFLGGTDATSAMTEWAMTEILRNRQVMKKVQDELAEVVGLNNIVEESHLPKLKYLDAVFKETFRLHPPLPFLLPRAPNKSCTVGGYT
VPKGSTIFLNVWAIQRDPQHWTNPSEFNPERFLNKGSEKWDYNGTNSKYFPFGSGRRRCPGIPLGEKMMMHILASLMHSFDWSLPRGEEHDLSDKFG
IAMKKKMPLVLIPSQRLSDHNLYM*

**>Design58683**

MASNELAFSALLVTLVLVLISWYKREISNSRKAGTPPLPPGPKGLPVVGFLPFLGPNLHLDFLTLVHKYGPIFKLYLGSNLHIVVNSADLAKVVTGEQD
ESFANRAQHIAGLATSYNASDIAFADNNANRRKLRKVLVHEVLSNVNLEASNAFRRREVRKTIKNVHEIIGNEVDINEIAFSTVLSVLTSIVFGKSMVK

GAKYSNLVAEMRKFVSGVVEIAGELNISDFFPMLARFDFQGVKRRMAQQMKIFDRIFETTIEERTGSTSGIIDQKVKEEGRKDFLQILLELLDQNTGTSITMTQLKALVVDIFLGGTDATSAMVEWAMTEIFRDKKVMKRVQDELAEVVGLHNIVEESHLPKLKYLDAVFKETFRLHPPLPFLLPRAPNKSCTVGGYTVPKGSTIFLNVWAIQRDPRYWENPSDFNPERFLNYKGSEKWDYSGTNLKFFPFGSGRRRCPGVPLGEKMMMHILASLLHSFDWSLPTGEKLDLSDKFGITLKKRKPLIAIPSMRLSDASLYM*

**>Design6444**

MASNELAFSALLVTLVLVLISWYKREISNSRKAGTPPLPPGPKGLPLVGFLPFLGPNLHLDFLTMAHQYGPIFKLYLGSNLHIVVNSADLAKVVTGEQDESFANRAQHIAGLATSYNASDILFADNNANRRKLRKVLVHEVLSNVNLEASNAYRRREVRKTIKNVHEIIGNEVDINEIAFSTVLSVLTSIVFGKSMVKGAKYSNLVADMRKFVSGVVEIAGGLNISDFFPMLARFDFQGVKRKMAQQMKMFDKIFESTVEERVGSTSGIIEEAVKQEGRKDFLQILLELLDQNTETSITMTQMKALVVDIFLGGTDATSAMVEWAMTEIFRDKKVMKRVQDELAEVVGLHNIVEESHLPKLKYLDAVFKETFRLHPPLPFLLPRAPNKSCTVGGYTVPKGSTIFLNVWAIQRDPQYWENPSEFNPERFLNYKGSEKWDYTGTNLKFFPFGSGRRRCPGVPLGEKMMMHILASLLHSFDWSLPNGQKLDLSDKFGITLKKRKPLIAVPSLRLSDASLYV*

**>Design84497**

MASNELAFSALLVTLVLVLISWYKREISNSRKAGTPPLPPGPRGLPLVGYLPFLGPQPHRSLSEISHRYGPIFKLQLGTKLWIVVNSAELAKVIHVEQDESFANRAPHIAGLATSYGGNDIAFAPNNANRRNLRKLLVQEVLSNVNLEASHAYRRHEVRKAVKYVYDRVGMDIDINEISFTTVINVFFNIWRMGFIDDQSNIGNLLEKIQKVICRIVEIAEGLNISDFFPVLARFDLQRVERKMKDQMKQFDKIIETTIKERMNSKSTNVEETVEQEGRKDFLQILLELLDQSTATSITMTQLKALVVDVFLGGTDATSAMVEWAMTEILRNRQVMKKVQDELAQVVGLHNVVEESHLPKLKYLDAVFKETFRLHPPLPFLLPRAPNKSCTVGGYTVPKGSTIFLNVWAIQRDPQHWTNPSEFNPERFLNYKGSEKWDYAGTNSKFFPLGSGRRRCPGIPLGEKMMMHILASLLHSFDWSLPTGQKLDLSDKFGIAMKKKKPLVVVPSLRLSDLSLYS*

**>Design91808**

MASNELAFSALLVTLVLVLISWYKREISNSRKAGTPPLPPGPKGLPIVGYLLFLGTNLHIHFSNLSQSYGPIFKFHLGNKLWVIVNTAELAKTIVVEQDESFANRWPHIAGLATSYGGNDIAFAPNNANRRNLRKVLVQEVLSNVNLEASHNYRRHEVRKAVKYVYDRVGMDIDINEISFTTVLNVFFNIIWRMGFEDDQTNVGNLLQKIQKVICRIVEIAEGLNISDFFPVLARFDVQRVERKMKDQMKQFDKIIETTIKERMNSKSTNVEETVEQKGRKDFLQILLELLDQNNETSITMTQMKALVVDIFLGGTDATSAMVEWAMTEILRNRQVMKKVQDELAEIVGLNNIVEESHLPKLKYLDAVFKETFRLHPPLPFLLPRAPNKTCTVGGYTVPKGSTIFLNVWAIQRDPQYWDNPSEFNPERFLNYKGSEKWDYNGTNLKFFPFGSGRRRCPGIPLGEKMMMHILASLMHSFDWSLPRGEELDLSDKFGIAMKKKKPLVVIPSLRLSDHNLYM*

**>Design49566**

MASNELAFSALLVTLVLVLISWYKREISNSRKAGTPPPLPPGPKGLPLVGSLPFLGPNIHQELTKITHQYGPIFKLYLGSKLHIVVNSAELAKVITAEQDESFANRAPHIAGLATSYGGNDIAFAPNNANRRNLRKVLVQEVLSNVNLEASHAYRRHEVRKAVKYVHERVGMEVDINKIAFSTVLSVLTNIVWAKSVVDDGANYSNEVQNIISRVVEIAGGLNISFFVMAQIDFQGVERRMKKQMKQFDKIIEMTIEERMNSSKNEAKTKEEGRKDFLQILLEQQQNTETSITMTQMKALVVDIFLGGTDATSAMVEWAMTEILRDQQVMKRVQDELEEIVGLNNVEESHLPKLKYLDAVFKETFRLHPPIPFLLPRAPIKSCTVGGYTVPEGATIFVNVWAIQRDPQRWENPSEFNPDRFLNRNGSTGKWDYSGTNLTFLPFGSGRRRCPGIPLGEKMMMHILASLLHSFDWKLPNGQKLELSERFGIALKKKKPLVAVPTKRLDVSLYM*

**>Design12854**

MAALNTHGSWWPAEGNGGKNDGDLPLALLAVITAALLPLLWYKRSISSSQNGAPPLPPGPKGLPVVGYLPFLGPNLHLDYLTMVHQYGPIFKIYLGSNLHIVVNSVDLAKVVTGEQDESFANRAQHIAGLATSYNASDIAFADNNANRRKLRKVLVHEVLSNVNLEASNAYRRREVRKTIKNVHEVIGNEVDINEISFSTVLSVLTSIVFGKSMVKGAKYSNLAADIRKFVSGVVEIAGGLNISDFFPMLARFDFQGVEQRMKTQMKMFDKIFETSVEERINSRSAIKEEAVKEEGRKDFLQILLELLEQNTETSITMTQMKALVVDVFLGGTDATSAMVEWAMTEIFRNRQVMKKVQDELAEIVGLHNIVEESHLPKLKYLDAVFKETFRLHPPLPFLLPRAPNKTCTVGGYTVPKGSTIFLNVWAIQRDPKYWDNPSDFNPERFLNYEGEKWDYNGTNLKFFPFGSGRRRCPGIPLGEKMMMHILASLLHSFNWSLPEGEDHDLSEKFGIAMKKKKPLIAIPSLRLSDHNLYM*

**>Design33105**

MASNELAFSALLVTLVLVLISWYKREISNSRKAGTPPLPPGPRGLPLVGYLPFLGPNLHQELTKMAHRYGPIFKLYLGSKLHIVVNSADLAKVVTGEQDESFANRAPHIAGLATSYNASDIAFADNNANRRKLRKVLVHEVLSNVNLEASHAYRRREVRKTIKNVHEIIGNEVDINEIAFSTVLSVLTSIVWGKSMVKGASNMIVEVRKFVSGVVEIAGELNISDFFPMLARFDFQGVERRMKKQMKLFDKIFESTVEERINSRSIIKEEAASKEENRRKDFLQILLELQEQNNTSITMTQMKALVVDIFLGGTDATSAMIEWAMTEILRNRRVMKKVQDELAEIVGLQNNVEEESHLPKLKYLDAVFKETFRLHPPLPFLLPRAPNKSCTVGGY

TIPKGSTIFLNVWAIQRDPQYWENPSEFNPERFLNYKGSEKWDYAGTNSKFFPLGSGRRRCPGVSLGEKMMMHILASLLHSFDWSLPTGQKLDLSDK
FGIALKKRKPLIAVPSPRLNDASLYM*
**>Design42565**
MASNELAFSALLVTLVLVLISWYKREISNSRKAGTPPLPPGPKGLPLVGYLPFLGPNLHHEFTKVSHRYGPIFKLYLGSKLHIVVNSADLAKVITSEQDE
SFANRAPHIAGLATSYGGNDIAFADNNANRRNLRKVLVHEVLSNVNLEASHAYRRHEVRKTIKSVHDMIGMEVDINEISFSTVLNVLTNIVWGKGLVE
GTKYSNLSEEIRKVVYRIVEIAEGLNISDFFPMLARFDLQGVERKMKKQMKQFDRIFENTISERTNNKNSNHGKQCLQILLELKTKTITTTQLKALVVD
IFLGGTDATSAMVEWAMTEILRNKKVMKRVQDELEEIVGLNNIVEESHIPKLKYLDAVFKETFRLHPPLPFLLPRAPSKSCTVGGYTVPKGATIFLNVW
AIQRDPRHWENPSEFNPDRFLNNNNGSTEKWDYSGTNLTFLPFGSGRRRCPGIPLGEKMMMHILASLMHSFDWSLPNGEEHDLSDKFGIALKKKKPL
VAIPTRRLSDENLYM*
**>Design26159**
MASNELAFSALLVTLVLVLISWYKREISNSRKAGTPPLPPGPYGLPLLGYLPFFLGPSLHHELTKMAHRYGPIFKLYLGSKLHIVVNSADLAKVITSEQD
ESFANRAPHIAGLATSYGANDIAFADNNANRRNLRKILVHEILSNVNLEASHAYRRREVRKTIKSVHDMIGMPVDINEMSFSTVVNVLTSIVWGNSMV
EGTKHSNLGEEIRKVVSEIVDIAEGLNISDFFPKLARFDLQGVEQKMKRKMKQFDWIFETTIEERINLKSTHGEDALKHEGRKDFLQILLELKDKKSIT
MTQLKALVVDIFLGGTDATSAMVEWAMAEILKNQKVMKKVQDELAEIVGLKNMVEESHLPKLKYLNATFKETFRLHTPLPVLLPRTPSKSCMVGGY
LIPRDSTVFLNVWAIQRDPQHWENPSEFNPERFLNYEGSGKWDYSGTNSKYFPFGSGRRRCPGIPLAEKMMLHILASLLHSFDWSLPKGEDHDLFEKF
GIALKKKKPLVAVPSPRLIDLSLYM*
**>Design49566**
MASNELAFSALLVTLVLVLISWYKREISNSRKAGTPPPLPPGPKGLPLVGSLPFLGPNIHQELTKITHQYGPIFKLYLGSKLHIVVNSAELAKVITAEQDE
SFANRAPHIAGLATSYGGNDIAFAPNNANRRNLRKVLVQEVLSNVNLEASHAYRRHEVRKAVKYVHERVGMEVDINKIAFSTVLSVLTNIVWAKSVV
DDGANYSNEVQNIISRVVEIAGGLNISFFVMAQIDFQGVERRMKKQMKQFDKIIEMTIEERMNSSKNEAKTKEEGRKDFLQILLEQQQQNTETSITMT
QMKALVVDIFLGGTDATSAMVEWAMTEILRDQQVMKRVQDELEEIVGLNNVEESHLPKLKYLDAVFKETFRLHPPIPFLLPRAPIKSCTVGGYTVPEG
ATIFVNVWAIQRDPQRWENPSEFNPDRFLNRNGSTGKWDYSGTNLTFLPFGSGRRRCPGIPLGEKMMMHILASLLHSFDWKLPNGQKLELSERFGIAL
KKKKPLVAVPTKRLDVSLYM*
**>Design58**
MASNELAFSALLVTLVLVLISWYKREISNSRKAGTPPLPPGPKGLPLVGYLPFLGPNLHQELTKMAHRYGPIFKLYLGSKLHIVVNSADLAKVVTGEQD
ESFANRAPHIAGLATSYNASDIAFADNNANRRKLRKVLVHEVLSNVNLEASHAYRRREVRKTIKNVHEIIGNEVDINEIAFSTVLSVLTSIVWGKSMVE
GAKYSNLVAEMRKFVSGVVEIAGELNISDFFPMLARFDFQGVERRMKKQMKLFDKIFESTVEERINSRKSVEKEDFLQILLELQEQNNETSITMTQMK
ALVVDIFLGGTDATSAMIEWAMTEILRNRQVMKKVQDELAEIVGLNNIVEESHLPKLKYLDAVFKETFRLHPPLPFLLPRAPNKSCTVGGYTIPKGSTI
FLNVWAIQRDPQYWENPSEFNPERFLNYKGSEKWDYAGTNSKFFPLGSGRRRCPGVSLGEKMMMHILASLLHSFDWSLPTGQKLDLSDKFGIALKK
RKPLIAVPSPRLNDASLYM*
**> Design4779**
MASNELAFSALLVTLVLVLISWYKREISNSRKAGTPPLPPGPRGLPLVGYLPFLGPNLHQELTKMAHRYGPIFKLYLGSKLHIVVNSADLAKVVTGEQD
ESFANRAPHIAGLATSYNASDIAFADNNANRRKLRKVLVHEVLSNVNLEASHAYRRREVRKTIKNVHEIIGNEVDINEIAFSTVLSILTSIVWGKSMVEG
AKYTNLVAEMRKFVSGVVEIAGELNISDFFPMLARFDFQGVERRMKKQMKLFDKIFESTVEERINSSKKEGRKDFLQILLELQEQNNETSITMTQMKA
LVVDIFLGGTDATSAMVEWAMTEILRNRQVMKKVQDELAEIVGLNNIVEESHLPKLKYLDAVFKETFRLHPPLPFLLPRAPNKSCTVGGYTIPKGSTIF
LNVWAIQRDPQYWENPSEFNPERFLNYKGSEKWDYAGTNSKFFPLGSGRRRCPGVSLGEKMMMHILASLLHSFDWSLPTGQKLDLSDKFGIALKKR
KPLIAVPSPRLNDASLYM*
**>Design3644**
MASNELAFSALLVTLVLVLISWYKREISNSRKAGTPPLPPGPRGLPLVGYLPFLGPNLHHELTKMAHRYGPIFKLYLGSKLHIVVNSADLAKVVTGEQD
ESFANRDPHIAGLATSYNANDIAFADNNANRRKLRKVLVHEVLSNVNLEASHAYRRREVRKTIKNVHEIIGNEVDINEIAFSTVLSVLTSIVWGKSMVK
GAKYPNLVAEMRKFVSGVVEIAGELNISDFFPMLARFDFQGVERRMKKQMKLFDKIFESTVEERINSRKKIARKDFLQILLELKEQNNETSITMTQMK
ALVVDIFLGGTDATSAMIEWAMTEILRNRQVMKKVQDELAEIVGLNNIVEESHLPKLKYLDAVFKETFRLHPPLPFLLPRAPNKSCTVGGYTIPKGSTI
FLNVWAIQRDPQYWENPSEFNPERFLNYEGSEKWDYAGTNSKFFPLGSGRRRCPGIPLGEKMMMHILASLMHSFDWSLPEGEKLDLSDKFGIALKKK
KPLIAVPSPRLNDASLYM*

**>Design2205**

MASNELAFSALLVTLVLVLISWYKREISNSRKAGTPPLPPGPRGLPLVGYLPFLGPNLHHELDKMAHRYGPIFKLYLGSKLHIVVNSADLAKVVTGEQ
DESFANRAPHIAGLATSYNASDIAFADNNANRRKLRKVLVHEVLSNVNLEASHAYRRREVRKTIKNVHEIIGNEVDINEIAFSTVLSVLTSIVWGKSMV
KGAKYGNLVAEMRKFVSGVVEIAGELNISDFFPMLARFDFQGVERRMKKQMKLFDKIFETTVEERTNSSKKKVKKDFLQVLLELKEQNNETSINMTQ
MKALVVDIFLGGTDATSAMIEWAMTEILRNKQVMKKVQDELAEVVGLNNIVEESHLPKLKYLDAVFKETFRLHPPLPFLLPRAPNKSCTVGGYTIPK
GSTIFLNVWAIQRDPQYWENPSEFNPERFLNYKGSEKWDYAGTNSKFFPLGSGRRRCPGVSLGEKMMMHILASLLHSFDWSLPEGQKLDLSDKFGIA
LKKKKPLIAIPSPRLNDASLYM*

**>Design4129**

MASNELAFSALLVTLVLVLISWYKREISNSRKAGTPPLPPGPRGLPLVGYLPFLGPNLHQELTKMAHRYGPIFKLYLGSKLHIVVNSADLAKVVTGEQD
ESFANRAPHIAGLATSYNASDIAFADNNANRRKLRKVLVHEVLSNVNLEASHAYRRREVRKTIKNVHEIIGNEVDINEIAFSTVLSVLTSIVLKGSNIAA
EMRKFVSGVVEIAGELNISDFFPMLARFDFQGVERRMKKQMKLFDKIFESTVEERINSRSAIKEEAVKEEGRKDFLQILLELQEQNNETSITMTQMKAL
VVDIFLGGTDATSAMIEWAMTEILRNRQVMKKVQDELAEIVGLNNIVEESHLPKLKYLDAVFKETFRLHPPLPFLLPRAPNKSCTVGGYTIPKGSTIFL
NVWAIQRDPQYWENPSEFNPERFLNYKGSEKWDYAGTNSKFFPLGSGRRRCPGVSLGEKMMMHILASLLHSFDWSLPTGQKLDLSDKFGIALKKRK
PLIAVPSPRLNDASLYM*

### DNA sequences used in this study

#### Natural sequences

**>CYP706X1**

ATGGCATCAAACGAGCTTGCTTTTTCAGCACTATTAGTTACACTTGTGTTAGTTCTTATTTCATGGTACAAAAGAGAAATCTCCAACTCCCGAAAG
GCAGGCACACCTCCATTGCCTCCGGGTCCAAAAGGTCTACCATTAGTTGGATCTCTTCCATTTCTTGGCCCTAATATTCACCAAGAACTAACCAAA
ATATCGCACCAATATGGCCCGATTTTTAAGCTATACCTTGGAAGCAAGCTTCACATTGTGGTGAACTCTGCTGAACTGGCAAAGGTCATAACCGCT
GAGCAAGACGAGAGCTTTGCTAACCGGGCCCCACATATTGCCGGGCTAGCAACAAGTTACGGTGGCAACGATATAGCATTTGCACCAAATAATG
CTAACCGACGTAACCTACGTAAAGTTTTGGTCCAAGAGGTCTTAAGTAATGTCAACCTTGAGGCGTCTCATGCGTATCGTAGACATGAGGTTAGA
AAGGCCGTTAAATATGTCTACGATAGGGTTGGTATGGACGTTGATATCAACGAGATATCGTTCTCAACGGTGTTGAATGTGTTTACAAACATAATAT
GGAGGAAAGGGTTTGTGGATGATGGGGCAAATTATGCTAATCTTAGTGAGAATATACAAAAAGTGATATGTAGAATTGTTGAGATCGCGGAAGGG
CTAAATATCTCGGACTTCTTCCCAATGCTTGCAAGGTTTGATCTTCAAGGAGTTGAACGAAAAATGAAGGATCAAATGAAGCAATTCGACAAGAT
TATAGAGCCTACTATCAAGGAGAGAATGAACTCGAGGTCTACAAACGTTGAAGAAACCGTTGAGCATAAAGGAAGGAAAGATTTTCTACAAATA
TTGTTAGAGCATACAGATCAAAAAAATGGGACATCAATCACCATGACTCAATTAAAAGCGCTTGTTGCGGATATATTTCTAGGAGGAACAGATGC
AACTTCAGCGATGGTGGAATGGGCAATGACAGAGATTTTTAGAGATCAAAAGGTGATGAAAAGGGTACAAGATGAACTAGAAGAAATAGTAGG
TCTAAACAACATCGTTGAAGAATCGCATATTCCAAAATTGAAGTACCTTGAGGCCGTGTGTAAGGAAACATTCCGTTTACATCCTCCAATACCTTT
CCTACTCCCTCGAGCACCAATTAAGTCTTGTACGGTCGGAGGTTACACAGTTCCAGAAGGCGCTACTATCTTTGTAAATGTATGGGCGATACAAA
GGGACCCACGACATTGGGAGAATCCATCTGAGTTCAACCCTGATCGGTTTTTGAACCGTAATGGATCTACCGAGAAATGGGACTATAGTGGTACG
AATCTTACGTTTTTACCATTTGGATCTGGTAGGAGAAGGTGTCCAGGAATTCCTTTAGGTGAGAAGATGATGATGCATATTTTGGCTTCATTAATGC
ACTCTTTTGATTGGAAATTGCCGAATGGCGAAGAACTTGACCTCTCTGAGAGATTTGGTATTGCACTCAAGAAAAAAAAGCCGCTTGTAGCCGT
CCCGACTAAAAGATTAAGTGACCTAAGCCTTTACATGTGA

**>Cnan706X**

ATGGCTGCTTCCTCCCACTCTAGTTTATGGGAAGAAGCCACCAACAACAAGCACGAAGTCGCTTTGGCCGTTTTCGTTGGTACTTTGGTTATTTT
GGCTATCTCTTGGTACAAGAGATCCAACTCTAGAACCGGTAAGGGTACACCTCCATTGCCTCCAGGTCCAAAGGGTTTACCAATCGTCGGTTATC
TACCATTTTTGTCTCCAAACTTGCATCACGAATTTGCTAAGATTGCCAACCAATACGGTCCAATTTTCAAGTTGTACTTGGGTTCTAAATTGCACAT
CGTTGTTAACTCCGCTGACTTGGCTAAGGTCGTTACTGGTGAACAAGACGAATCTTTCGCTAACAGAGACCCGCACATTGCTGGCTTGACCGCAT
CTTACGGTGCCTCCGATGTTGCTTGGCAAAACAATAACTCCAACAGAAGAAACTTGAGAAAGGTGTTGGTCCATGAAGTTTTGTCAAACAAGAA
CTTGGAAGCTTCTCACGCTTACAGAAGATCTGAAGTTCGTAAGACCATCAAGAACGTTCACGACATGATTGGTACCGCTGTTGACATCAACGAA
GTTTCCTTTTCTACTGTTTTGAATATTCTTACTCACATTGTTTGGGGTAACTCTTTCGTCGAAGGTGCTAAATACCCAAATCTAGCCGCTGATATCA

GAAAGGTTGTCTTGGATATTGTTGAAATTGCTGAAGGTTTGAACCTCTCCGATTTCTTCCCAATGTTGGCTAGATTCGATTTCCAAGGTGTTGAAC
GTCGTGTCAAAGCTCAAGTTAAGAAGTTTGACCACATTTTCGAAACCACTATTGAAGAAAGAACCAACTCCAAGTCTAAGGTCTCTGAAGAAGC
TGTCAAGCAAGAAGGTAGAAAGGATTTCTTGCAAATCTTGTTGGAATTGTTAGACCAAAACACTGCTACCTCCATCAACATGACCCAATTGAAG
GCTTTGGTTGTCGATATCTTCTTGGGTGGTACTGATGCTACTGCTGCCATGACTGAATGGGCTATGGCCGAAATCCTACGTAACCCAAAGGTTATG
AAGAAGGTTCAAGATGAATTGGCTGAAGTCGTCGGCTTAAACAACATCGTCGAAGAATCTCATTTGCCAAAGTTGAAGTACTTGGACGCTGTTT
TCAAGGAAACTTTCAGATTGCACACTCCATTGCCATTCTTATTGCCAAGAACTCCAGACAAGTCTTGTGTCGTTGGTGGTTACACCGTTCCAAAG
GGTGCTACTGTCTTCTTGAACGTCTGGGCCATCCAAAGAGACCCACAAAACTGGGAAAACCCTTCTGAATTTAACCCAGAAAGATTCTTGAATA
ACAAAGGTTCCGAAAAGTGGGACTACTCTGGTACCAATTCCACTTACTTCCCATTCGGTTCTGGTAGAAGAAGATGTCCAGGTATTTTGTTGGGT
GAAAAAATGATGATGCATATCCTCGCCTCGTTGATGCACTCCTTCGACTGGTCTTTGCCAAAGGGTGAAGAGTTAGATTTGTCAGACAAGTTCGG
TATTGCTATGAAAAAGAAGATGCCATTAGTTGTCATCCCATCTTTGAGATTAAGTGACTTGTCTTTATACTCC

**>Lsal706X**

atggcttctaacgaattggctttttcagcattgttggttactttggttttggttttgatttcttggtacaagagagaaatctctaactcaagaaaggcaggtacaccaccattaccaccaggtccatacggcttgccactagtcggttatttgccattct
tgggtccttccctacaccacgaattgactaagatggcccacagatacggtccaatcttcaagttgtacttgggttccaaattgcacattgttgttaattctgctgatctggctaaggttatcacttccgaacaagatgaatcctttgctaacagagc
tccacacattgctggtttggccacatcttacggtgctaacgacattgctttcgctgacaacaacgccaacagaagaaacttgagaaagatcttggtccacgaaatcttgtccaacgttaacttggaagcttctcacgcatacagaagaagaga
agttagaaagactatcaagagtgttcatgacatgattggtatgccagttgatattaacgaaatgtctttctccactgttgttaacgtattgacctctattgtctggggtaactccatggttgaaggtaccaagcattctaacctaggtgaagaaataa
gaaaggttgtctctgaaatcgtcgatattgctgaaggtttgaacatctctgacttcttcccaaaattagctagattcgacttgcaaggtgtcgaacaaaagatgaagagaaagatgaagcaatttgactggattttcgaaaccaccattgaagaa
cgtatcaacttgaagtctacccacggtgaagatgccctaaagcacgaaggtcgtaaggatttcttgcaaatcttattggaattgaaggacaagaaatcaatcaccatgactcaattgaaggctctcgtcgttgacatcttcttgggtggtactga
tgctacttctgccatggttgaatgggctatggctgagattttgaagaaccaaaaggttatgaaaaaggtccaagacgaattagccgaaatcgttggtttaaagaacatggtcgaagaatctcatttacctaaattaaagtacttgaatgctacctt
caaggaaactttcagattgcataccccactaccagtcttgttgccaagaactccatctaagagttgtatggtcggtggttacttgatcccaagagattctactgtcttcttaaatgtctgggccattcaaagagacccacaacactgggaaaacc
catccgaattcaacccagaaagatttttgaactacgaaggttcaggtaagtgggactactctggtaccaactccaagtacttcccattcggttccggtcgtcgtagatgtccaggtattccattggctgaaaagatgatgttgcacatcctggct
tccttgttgcactctttcgattggtctttgccgaagggtgaagaccacgatttgtttgaaaagttcggtatcgctttgaaaaagaagaagccattggttgctgttccatctccaagattgattgacttgtctttatacatgtaa

#### Ancestral DNA sequences

**>ancXY**

ATGGCTTCTAACGAATTGGCTTTTTCAGCATTGTTGGTTACTTTGGTTTTGGTTTTGATTTCTTGGTACAAGAGAGAAATCTCTAACTCAAGAAAG
GCAGGTACACCACCATTGCCACCAGGTCCAAGAGGTTTACCAGTTGTTGGTTATTTGCCATTTTTAGGTCCAAATTTGCATCAAGAATTCACTAAG
ATGGCTCATAGATACGGTCCAATTTTTAAGTTGCATTTGGGTTCAAAGTTGCATATCGTTGTTAATTCTGCTGATTTGGCAAAAGTTGTTGCTAGAG
AACAAGATGAAACTTTCGCAAACAGAAATCCACCAGTTGCTGCATTAGCTATTACATATGGTGGTCAAGATATCGCATGGTCTAACAACAACTCA
AACTGGAGAAATTTGAGAAAGGTTTTGGTTCATGAAGTTTTGTCAAATAAGAATTTGGAAGCATCTAGATCTTTTAGAAGAAGAGAAGTTAGAA
AGACTATTAAAAATGTTTACGAAAAGATTGGTACAGAAATTGATATTAATGAAATTGCTTTTTCTACTGAATTGAACGTTTTGACATCAATGGTTTG
GGGTAAATCTTTGGTTGAAGGTGAAAAGTACTCAAATTTGGGTGACGAATTCAGAGAAGTTGTTTCTAAGATCGTTGAAATCTTGGGTGCACCA
AACATCTCTGATTTCTTTCCAATCTTGGCTTGGTTCGATTTGCAAGGTGTTGAAAGAGAAATGAAAAGACAATTGAAGCAATTGGATAGAATCTT
CGAATCAATTATTGAAGAAAGAATTAATTCTAATTCAACTAAATCTGAAGAAGCTGTTGAACATGAAGGTAGAAAGGATTTCTTGCAAATCTTGTT
GGAATTGAAGGATCAAAAGGATGCAACTTCAATTAATATCACACAAATTAAAGCTTTGTTAGTTGATATTTTGTTAGGTGGTACTGATACTACATCT
ACAATGGTTGAATGGGCTATGGCAGAAATCTTGCAAAACCAAAAAGTTATGAAGAAAGTTCAAGATGAATTGGCAGAAATCGTTGGTTTGAACA
ACATCGTTGAAGAATCACATTTGCCAAAGTTGAAGTATTTGGATGCTGTTATTAAAGAAACTTTTAGATTACATCCACCATTGCCATTGTTAATTCC
AAGATCTCCAAATCAATCATGTACTGTTGGTGGTTACACAATCCCAAAGGGTTCTACAGTTTTCTTGAATGTTTGGGCAATTCATAGAGATCCACA
ATACTGGGATAACCCATTGGAATTCAATCCAGAAAGATTTTTGAACAGAGAAGGTACAGATAAATGGGATTACAACGGTAACAATTTGAAGTTCT
TGCCATTTGGTTCAGGTAGAAGAAGATGTCCAGGTATTCCATTGGGTGAAAAGATGTTGATGTACATCTTGGCTTCTTTGTTGCATTCATTCGATT
GGTCTTTGCCAAAGGGTGAAGAACATGATTTGTCTGATAAGTTCGGTATCGCTTTGAAGAAAAGAAAGCCATTGATCGCAATTCCATCACAAAGA
TTACCAGATGCTTCTTTGTACATGTAA

**>ancX**

ATGGCTTCTAACGAATTGGCTTTTTCAGCATTGTTGGTTACTTTGGTTTTGGTTTTGATTTCTTGGTACAAGAGAGAAATCTCTAACTCAAGAAAG
GCAGGTACACCACCATTGCCACCAGGTCCAAGAGGTTTGCCATTAGTTGGTTATTTGCCATTTTTAGGTCCAAATTTGCATCAAGAATTGACTAAG
ATGGCTCATAGATACGGTCCAATTTTTAAGTTGTACTTAGGTTCAAAGTTACATATTGTTGTTAATTCTGCTGATTTGGCAAAAGTTGTTACTGGTG

AACAAGATGAATCATTTGCTAATAGAGCACCACATATTGCTGGTTTAGCAACATCTTATGGTGCAAATGATATCGCTTTCGCAGATAACAACGCTA
ACAGAAGAAATTTGAGAAAGGTTTTGGTTCATGAAGTTTTGTCAAATGTTAATTTGGAAGCTTCTCATGCATACAGAAGAAGAGAAGTTAGAAA
GACAATTAAAAATGTTCATGAAATGATCGGTAACGAAGTTGATATCAACGAAATCGCATTTTCTACTGTTTTGAACGTTTTGACATCAATCGTTTG
GGGTAAATCTATGGTTGAAGGTGCTAAGTACTCAAATTTGGGTGAAGAAATCAGAAAGGTTGTTTCTGGTATTGTTGAAATTGCAGGTGGTTTGA
ACATCTCAGATTTCTTTCCAATGTTGGCTAGATTCGATTTGCAAGGTGTTGAAAGAAAGATGAAGAGACAAATGAAGCAATTCGATAAGATCTTC
GAATCTACTATTGAAGAAAGAATTAATTCTAAGTCAACAAACGTTGAAGAAGCAGTTAAGCATGAAGGTAGAAAGGATTTCTTGCAAATCTTGTT
GGAATTGAAAGATCAAAAGAATGAAACTTCAATCACTATGACACAATTGAAGGCTTTGGTTGTTGATATTTTCTTGGGTGGTACTGATGCTACATC
TGCAATGGTTGAATGGGCAATGACAGAAATCTTGAGAAACCAAAAAGTTATGAAGAAAGTTCAAGATGAATTGGCTGAAATCGTTGGTTTGAAC
AACATCGTTGAAGAATCACATTTGCCAAAGTTGAAGTATTTGGATGCAGTTTTTAAAGAAACTTTTAGATTACATCCACCATTGCCATTTTTGTTAC
CAAGAGCTCCAAATAAGTCATGTACTGTTGGTGGTTACACAATCCCAAAGGGTTCTACAATTTTCTTGAACGTTTGGGCTATTCAAAGAGATCCA
CAATACTGGGAAAATCCATCAGAATTCAATCCAGAAAGATTTTTGAACTACGAAGGTTCTGAAAAATGGGATTACTCTGGTACTAACTCAAAGTT
TTTCCCATTCGGTTCTGGTAGAAGAAGATGTCCAGGTATTCCATTGGGTGAAAAGATGATGATGCATATCTTGGCTTCTTTGTTGCATTCATTCGAT
TGGTCTTTGCCAAAGGGTGAAGAACATGATTTGTCTGATAAGTTCGGTATCGCTTTGAAGAAAAGAAAGCCATTGATTGCAGTTCCATCTCCAAG
ATTATCTGATGCTTCATTGTACATGTAA

**>ancX1**

ATGGCTTCTAACGAATTGGCTTTTTCAGCATTGTTGGTTACTTTGGTTTTGGTTTTGATTTCTTGGTACAAGAGAGAAATCTCTAACTCAAGAAAG
GCAGGTACACCACCATTGCCACCAGGTCCAAGAGGTTTGCCATTAGTTGGTTATTTGCCATTTTTAGGTCCAAATTTGCATCAAGAATTGACTAAG
ATGGCTCATAGATACGGTCCAATTTTTAAGTTGTACTTAGGTTCTAAGTTACATATTGTTGTTAATTCAGCTGATTTGGCAAAAGTTGTTACTGGTG
AACAAGATGAATCTTTTGCTAATAGAGCACCACATATTGCTGGTTTGGCAACATCTTACAACGCATCAGATATCGCTTTCGCAGATAACAACGCTA
ACAGAAGAAAGTTGAGAAAAGTTTTGGTTCATGAAGTTTTATCTAATGTTAATTTGGAAGCTTCACATGCATACAGAAGAAGAGAAGTTAGAAA
GACTATTAAAAATGTTCATGAAATCATTGGTAATGAAGTTGATATTAATGAAATCGCTTTTTCTACTGTTTTGTCAGTTTTGACATCTATCGTTTGGG
GTAAATCTATGGTTAAGGGTGCTAAGTACTCAAATTTGGTTGCAGAAATGAGAAAGTTCGTTTCTGGTGTTGTTGAAATCGCTGGTGAATTGAAC
ATCTCAGATTTCTTTCCAATGTTGGCTAGATTCGATTTCCAAGGTGTCGAAAGAAGAATGAAGAAACAAATGAAGTTGTTCGATAAGATCTTCGA
ATCTACAGTTGAAGAAAGAATTAATTCTAGATCAGCTATTAAAGAAGAAGCAGTTAAGGAAGAAGGTAGAAAGGATTTCTTGCAAATCTTGTTGG
AATTGCAAGAACAAAACAATGAAACTTCTATCACTATGACACAAATGAAGGCTTTGGTTGTTGATATTTTCTTGGGTGGTACTGATGCTACATCAG
CAATGATTGAATGGGCAATGACAGAAATCTTGAGAAACAGACAAGTTATGAAGAAAGTTCAAGATGAATTGGCTGAAATCGTTGGTTTGAACAA
CATCGTTGAAGAATCTCATTTGCCAAAGTTGAAGTATTTGGATGCAGTTTTTAAAGAAACTTTTAGATTACATCCACCATTGCCATTTTTGTTACCA
AGAGCTCCAAATAAGTCTTGTACTGTTGGTGGTTACACAATCCCAAAGGGTTCAACAATTTTCTTGAACGTTTGGGCAATTCAAAGAGATCCACA
ATACTGGGAAAATCCATCTGAATTCAATCCAGAAAGATTTTTAAACTACAAAGGTTCAGAAAAATGGGATTACGCTGGTACTAATTCTAAATTTTT
CCCATTGGGTTCAGGTAGAAGAAGATGTCCAGGTGTTTCTTTGGGTGAAAAGATGATGATGCATATCTTGGCATCATTGTTGCATTCTTTCGATTG
GTCATTGCCAACAGGTCAAAAGTTGGATTTGTCTGATAAGTTCGGTATCGCTTTGAAGAAAAGAAAGCCATTGATTGCAGTTCCATCTCCAAGAT
TAAATGATGCTTCATTGTACATGTAA

**>ancX2**

ATGGCTTCTAACGAATTGGCTTTTTCAGCATTGTTGGTTACTTTGGTTTTGGTTTTGATTTCTTGGTACAAGAGAGAAATCTCTAACTCAAGAAAG
GCAGGTACACCACCATTGCCACCAGGTCCAAGAGGTTTGCCATTAGTTGGTTATTTGCCATTTTTAGGTCCAAATTTGCATCAAGAATTGACTAAG
ATCGCTCATAGATACGGTCCAATTTTTAAGTTGTACTTAGGTTCAAAGTTACATATTGTTGTTAACTCTGCTGATTTGGCAAAAGTTATTACTGGTG
AACAAGATGAATCATTTGCTAATAGAGCACCACATATTGCTGGTTTAGCAACATCTTATGGTGCTAATGATATCGCTTTCGCAGATAACAACGCAA
ACAGAAGAAATTTGAGAAAGGTTTTGGTTCATGAAGTCTTGTCAAATGTTAATTTGGAAGCTTCTCATGCATACAGAAGAAGAGAAGTTAGAAA
GACAATTAAAAATGTTCATGATATGATTGGTATGGAAGTTGATATCAACGAAATCTCTTTTTCAACTGTTTTGAACGTTTTGACATCAATCGTTTGG
GGTAAAGGCATGGTTGAAGGTGCTAAGTACTCAAATTTGGGTGAAGAAATCAGAAAGGTTGTTTCTGGTATCGTTGAAATCGCTGAAGGTTTGA
ACATCTCTGATTTCTTTCCAATGTTGGCAAGATTCGATTTGCAAGGTGTTGAAAGAAAGATGAAGAGACAAATGAAGCAATTCGATAAGATCTTC
GAAACTACAATTGAAGAAAGAATTAATTCTAAGTCAACTAACGTTGAAGAAGCTGTTAAGCATGAAGGTAGAAAGGATTTCTTGCAAATCTTGTT
GGAATTGAAGGATCAAAAGAATGGTACATCAATCACTATGACACAATTGAAGGCATTGGTTGTTGATATTTTCTTGGGTGGTACTGATGCTACATC
TGCAATGGTTGAATGGGCTATGACTGAAATCTTGAGAAACCAAAAAGTTATGAAGAAAGTTCAAGATGAATTGGCAGAAATCGTTGGTTTGAAC

AACATCGTTGAAGAATCTCATTTGCCAAAGTTGAAGTATTTGGATGCTGTTTTTAAAGAAACTTTTAGATTACATCCACCATTGCCATTTTTGTTAC
CAAGAACACCAAATAAGTCATGTACTGTTGGTGGTTACACAATCCCAAAGGGTTCTACTATTTTCTTGAACGTTTGGGCAATTCAAAGAGATCCA
CAACATTGGGAAAATCCATCAGAATTCAATCCAGAAAGATTTTTGAACTACGAAGGTTCTGAAAAATGGGATTACTCTGGTACAAACTCAAAGTT
TTTCCCATTCGGTTCTGGTAGAAGAAGATGTCCAGGTATTCCATTGGGTGAAAAGATGATGATGCATATCTTGGCTTCTTTGTTGCATTCATTCGAT
TGGTCTTTGCCAAAGGGTGAAGAACATGATTTGTCAGAAAAGTTCGGTATCGCTTTGAAGAAAAAGAAACCATTGGTTGCAGTTCCATCTCCAA
GATTGTCTGATTTGTCATTGTACATGTAA

**>ancX3**

ATGGCTTCTAACGAATTGGCTTTTTCAGCATTGTTGGTTACTTTGGTTTTGGTTTTGATTTCTTGGTACAAGAGAGAAATCTCTAACTCAAGAAAG
GCAGGTACACCACCATTGCCACCAGGTCCAAAAGGTTTGCCATTAGTTGGTTCTTTGCCATTTTTAGGTCCAAACATCCATCAAGAATTGACTAA
GATCTCACATCAATACGGTCCAATTTTTAAGTTGTACTTAGGTTCTAAGTTGCATATCGTTGTTAACTCAGCTGAATTGGCAAAAGTTATTACTGCT
GAACAAGATGAATCTTTTGCTAATAGAGCACCACATATTGCTGGTTTAGCAACATCATATGGTGGTAATGATATCGCTTTCGCACCAAACAACGCA
AACAGAAGAAATTTGAGAAAGGTTTTGGTTCAAGAAGTTTTGTCTAACGTTAATTTGGAAGCTTCACATGCATACAGAAGACATGAAGTTAGAA
AGGCTGTTAAGTACGTTTACGATAGAGTTGGTATGGATGTTGATATCAACGAAATCTCTTTTTCAACTGTTTTGAATGTTTTAACAAATATTATTTG
GAGAAAAGGTTTTGTTGATGATGGTGCTAATTATGCAAATTTGTCTGAAAAGATTCAAAAAGTTATTTGTAGAATTGTTGAAATTGCTGAAGGTTT
GAACATCTCAGATTTCTTTCCAATGTTGGCAAGATTCGATTTGCAAGGTGTTGAAAGAAAGATGAAGGATCAAATGAAGCAATTCGATAAGATCA
TCGAAACTACAATTAAAGAAAGAATGAATTCTAAGTCAACTAACGTTGAAGAAACAGTTGAACATAAGGGTAGAAAGGATTTCTTGCAAATCTT
GTTGGAACATACTGATCAAAAGAATGGTACATCTATCACTATGACACAATTGAAGGCTTTGGTTGCAGATATTTTCTTGGGTGGTACTGATGCTAC
ATCAGCAATGGTTGAATGGGCTATGACAGAAATTTTTAGAGATCAAAAAGTTATGAAGAGAGTTCAAGATGAATTGGAAGAAATCGTTGGTTTGA
ACAACATCGTTGAAGAATCTCATATTCCAAAGTTGAAGTATTTGGAAGCTGTTTGTAAGGAAACTTTTAGATTGCATCCACCAATTCCATTTTTGTT
ACCAAGAGCACCAAATAAGTCTTGTACTGTTGGTGGTTACACAGTTCCAGAAGGTGCTACAATCTTCGTTAACGTTTGGGCAATCCAAAGAGATC
CAAGACATTGGGAAAATCCATCTGAATTCAATCCAGATAGATTTTTAAACTGTAATGGTTCTACTGAAAAGTGGGATTACTCAGGTACTAATTTGA
CATTTTTACCATTCGGTTCAGGTAGAAGAAGATGTCCAGGTATTCCATTGGGTGAAAAGATGATGATGCATATCTTGGCTTCTTTGATGCATTCATT
CGATTGGAAGTTACCAAATGGTGAAGAATTGGATTTGTCTGAAAGATTCGGTATCGCTTTGAAGAAAAAGAAACCATTGGTTGCAGTTCCAACA
AAAAGATTGTCTGATTTGTCATTATACATGTAA

**>ancX-16**

ATGGCATCAAACGAATTGGCTTTTTCTGCATTGTTGGTTACTTTGGTTTTGGTTTTGATCTCTTGGTACAAGAGAGAAATCTCTAATTCAAGAAAA
GCTGGTACACCACCATTACCACCAGGTCCAAGAGGTTTGCCAGTTGTTGGTTATTTGCCATTTTTAGGTCCAAATTTGCATCAAGAATTCACTAAG
ATGGCACATAGATACGGTCCAATTTTTAAGTTGCATTTGGGTTCAAAGTTGCATATCGTTGTTAATTCTGCTGATTTGGCAAAAGTTGTTGCTAGAG
AACAAGATGAAACTTTCGCAAACAGAAATCCACCAATTGCTGGTTTAGCAACATCATACGGTGCTAATGATATCGCTTTCGCAAACAACAACTCT
AACTGGAGAAATTTGAGAAAGGTTTTGGTTCATGAAGTTTTGTCAAATAAGAATTTGGAAGCATCTAGATCTTTTAGAAGAAGAGAAGTTAGAA
AGACTATTAAAAATGTTTACGAAAAGATTGGTACAGAAATTGATATTAATGAAATTGCTTTTTCTACTGAATTGAACGTTTTGACATCAATGGTTTG
GGGTAAATCTTTGGTTGAAGGTGAAAAGTACTCAAATTTGGGTGACGAATTCAGAGAAGTTGTTTCTAAGATCGTTGAAATCGCTGGTGCACCA
AACATCTCTGATTTCTTTCCAATCTTGGCATGGTTCGATTTGCAAGGTGTTGAAAGAGAAATGAAAAGACAAATGAAGCAATTCGATAGAATCTT
CGAATCAATTATTGAAGAAAGAATTAATTCTAATTCAACTAAATCTGAAGAAGCTGTTGAACATGAAGGTAGAAAGGATTTCTTGCAAATCTTGTT
GGAATTGAAGGATCAAAAGGATGCAACTTCAATTAATATCACACAAATTAAAGCTTTAGTTGTTGATATTTTCTTGGGTGGTACTGATGCTACATCT
GCAATGGTTGAATGGGCTATGGCAGAAATCTTGCAAAACCAAAAAGTTATGAAGAAAGTTCAAGATGAATTGGCAGAAATCGTTGGTTTGAACA
ACATCGTTGAAGAATCACATTTGCCAAAGTTGAAGTATTTGGATGCTGTTATTAAAGAAACTTTTAGATTACATCCACCATTGCCATTGTTATTGCC
AAGATCTCCAAATCAATCATGTACTGTTGGTGGTTACACAATCCCAAAGGGTTCTACAGTTTTCTTGAATGTTTGGGCTATTCATAGAGATCCACA
ATACTGGGATAACCCATTGGAATTCAATCCAGAAAGATTTTTGAACAGAGAAGGTACAGATAAATGGGATTACAACGGTAACAATTTGAAGTTCT
TGCCATTTGGTTCAGGTAGAAGAAGATGTCCAGGTATTCCATTGGGTGAAAAGATGTTGATGTACATCTTGGCTTCTTTGTTGCATTCATTCGATT
GGTCTTTACCAAAAGGTGAAGAACATGATTTGTCTGATAAGTTCGGTATCGCTTTGAAGAAAAGAAAGCCATTGATCGCAATTCCATCACAAAGA
TTACCAGATGCTTCTTTGTACATGTAA

#### Other subfamily DNA sequences

>CYP73A1 (XM_020846215)

atggatctggtgttggttgagaaggcgttactcggcctcttcgcggcggtgactctcgccatcgtcgtctctaagctcagcgggaagaagctcaagctcccgccagggccccttccggtgccgatcttcggcaactggctccaggtcggc
gacgacctcaatcaccgcaatctcgcatccctggcaaagaagttcggcgatatccttctcctccgaatgggccagcgcaatctcgttgtcgtctcgtcgccggatcatgcccgtgaggtcctccacactcagggagtcgaatttggttcccg
aactcgcaacgtcgtctttgacatcttcactggaaagggtcaggacatggtcttcactgtatacggcgagcattggcggaagatgcgccgcataatgaccgtccccttcttcaccaacaaggtcgtgcagcaataccgctatgggtgggaa
gatgaggccgcaagcgtcgttgaggatgtccgtaaaaaccctaagtcggccaccgagggggttgtgatccgccgccgcctgcaactcatgatgtacaacaatatgtaccgcatcatgttcgaccgaaggtttgagagcgaggacgatcc
ccttttcatgaagctgagggcccttaataacgagcggagtcgcctcgcgcagagcttcgagtacaactatggagattttatccccattctccggccgttcctgaggggatatctcaagatctgcaaggatgtgaaagatgtgcgactcaacct
attcaagaactatttcgttcaggagaggaagaaattgtcgagcacgaagccgatggataacgccggactcaagtgcgccatcgatcatatactggatgcggaaaagaagggcgagattaacgaggataacgttctctacatcgtcgaaaa
catcaacgttgctgcaatcgagacaactctatggtcgatcgagtggggcatcgccgagctagtaaaccacccggaagtccagcgcaagctccgcgaagaactcgacaccgtccttggccgcggcaaccaaatcacagagcccgacac
ccacaagctcccctacctccaggccgtcataaaagaaactctccggcttcgcatggccatccctctcctcgtcccccacatgaatctccacgacgccaagctgggaggctacgaaatccccgccgagagcaagatcctcgtcaacgcct
ggtggctcgccaacaaccccgcccaatggaaaaaccccgatcagttccgccccgagcggttcctcgaagaggagtcaaaagttgaggccaacggcaacgacttccgcttcatcccttcggcgtcggccgccgcagctgccctggca
tcatcctcgccctccctattctcggaatcactctcggccgcctcgtccagaactttgagctcctgccgccgccgggtatggagaagctcgatacttctgagaaaggagggcagttcagtctccatatacttaagcattacaccgtcgtcgcca
agcccagggaattctaa
>CYP706A3 (NCBI Number: NM_123829)
atgacagacatttccagtctctttagaaaccgaagtcgcaaggaccaacttgactatggtttaactgtgatagttatatcaactctttgttggtgtctttggctctacgccaaatgcaaacggcggtctccaccgttgccgccgggaccttgggg
ccttcccattatcggaaaccttccattcctccaaccggagcttcacacctacttccaagggctggctaaaaagcacggtcccattttcaaactctggctcggggccaagctcacaatcgtggtcacctcttctgaagtagcgcaagagattctc
aaaacgaatgacatcatcttcgcgaaccacgatgtccctgctgtgggccctgttaacacgtatggtggtacggagatcatttggtcgccgtatggaccaaagtggcggatgctgaggaagctatgtgttaataggatactgagaaacgcca
tgttggattcctccactgacctccgtcgccgagagactcggcaaaccgtccggtatttggcggaccaggctcgggtcggttcaccagttaacctgggagaacaaatattcttgatgatgttgaatgttgtgacgcagatgctatggggaaca
acggttaaggaagaagagagggaggttgttggagccgagttcttagaggtgattagagagatgaacgacctcctgctagtgcccaatatctccgactttttcccggtattgagccggtttgatcttcagggtttggctaagcgtatgcgaaga
ccggctcaaagaatggatcagatgttcgaccggatcattaaccaacggttggggatggatagggacagtagtgacgggagagctgtggattttctagacgtcttgttgaaagtcaaggatgaagaagctgaaaagacgaagttgaccatg
aacgatgtcaaggccgtactcatggacatggtgctcggtggtacagacacatcactgcacgtaatagaattcgcgatggccgagctattacacaaccccgatatcatgaagagagctcaacaagaggttgacaaagttgtgggaaaaga
aaaagttgtggaagaatctcacatctctaaacttccatacattctcgccattatgaaagaaactctaaggcttcacacggttgctccactccttgttcctcggcgcccgtcacaaaccaccgtggtgggcggcttcaccatccctaaagattcaa
agattttcatcaacgcgtgggcaatccataggaacccgaatgtatgggagaatccactgaagtttgatcccgacaggtttctggatatgtcttatgacttcaaaggaaatgacttcaattatctaccgtttgggtctggtcgtaggatatgcgtgg
gaatggctatgggcgagagggttgttctttacaatctcgctacgtttttgcattcttttgattggaaaattccgcaaggagagagagtggaggtcgaggagaagtttgggatcgtgttggagctaaagaatccacttgttgccacgcccgttcta
aggttgtccgatccaaatctttatctctag
>CYP706B1 (NCBI Number: XM_016849700)
atgttgcaaatagctttcagctcgtattcatggctgttgactgctagccaccagaaagatggaatgttgttcccagtagctttgtcatttttggtagccatattgggcatttcattgtggcacgtatggaccataagaaagccaaagaaagacatc
gccccattaccgccgggtccccgtgggttgccaatagtgggatatcttccatatcttggaactgataatcttcacttggtgtttacagatttggctgcagcttatggtcccatctacaagctttggctcggaaacaaattatgcgtagtcattagct
cggcaccactggcgaaagaagtggttcgtgacaacgacatcacattttcggagagggatcctcccgtttgtgcaaagattattacctttggcctcaatgatattgtatttgattcttacagtagtccagattggagaatgaagagaaaagtgctg
gtacgtgaaatgcttagccatagtagcattaaagcttgttatggtctaaggagggaacaagtgcttaaaggcgtacaaaatgttgctcaaagtgctggcaagccaattgattttggtgaaacggcatttttaacatcaatcaatgcgatgatgag
catgctgtggggtggcaaacagggaggagagcagaaaggggccgacgtttggggccaatttcgagatctcataaccgaactaatggtgattcttggaaaaccaaacgtttctgatattttcccggtgcttgcaaggtttgacatacaggga
ttggagaaagagatgactaaaatcgttaattctttcgataagcttttcaactccatgattgaagaaagagagaactttagcaacaaattgagcaaagaagatggaaacactgaagcaaaagacttcttgcagcttcttttggaactcaagcaga
agaacgatagcggaatatcgataacaatgaatcaagtcaaggccttgctcatggacattgtggtcggtggaactgatacaacatcaaccatgatggaatggacaatggctgaactaattgcaaatcctgaagcaatgaaaaaggtgaagc
aagaaatagacgatgttgtcggttcggatgccgccgtcgatgagactcacttgcctaagttgcgctatctagatgctgcagtaaaggagaccttccgattgcacccaccgatgccactccttgtaccccgttgcccaggtgacttaagcaac
gttggtggctatagcgtaccaaagggcaccagggtcttcttaaacatttggtgtattcagagggatccacagctttgggaaaatcctttagaattcaagcctgagaggttcttgactgatcatcaaaagctcgattatttaggaaacgattcccg
gtacatgccgtttggttccggaaggagaatgtgtgccggagtatctctcggtgaaaagatgttgtattcctccttggcggcaatgatccatgcttatgattggaatttggccgacggtgaagaaaatgacttgattggcttatttggaattatcat
gaagaaaaagaagcctttaattcttgttcctacaccaagaccatcaaatctccagcactatatgaagtaa
>CYP706C55 (NCBI Number: MH974544)
atgtctccgtccatctccctttcccttctccacaatctcgaatcttccctctcatcgaacgatttctcgcttctgagcttactttttctcctctcgggcgctttggcaattggcttttgggcatggcgtcgttccgcagagaagaagaactcgctgccg
ctgccaccgggaccggccggtctccccctcgtcgggaacctgcccttcctcgacccggagctccacacctacttcgcaaccctcgcgatgacgtacggccccatcctgaagctccagcttggcaagaagctcggtatcgtcgtgacgtc
cccagccaccgcccgcgaggtcctcaaggataacgacgtcacgttcgccaaccgcgacgtgcccatcgctgggagggtcgcgttctacggcgggtcagacatcgtgtggaattcatacgggcccgagtggaggatgttccggaaggt
gtgcggcctcaagatgctgagcaaccacgcgctggactccgtctacgagctccggcggagggaggtccgaaggaccgtggggtacttcctgctccaggccgggtcgccggtgaacgttggggagcagatgttcttgacggtgcttaa
cgtgatcacgagcatgttgtggggtgggacggcgcagggcgaggagaaagagagcctgggtgctgatttcaggcaggcggtgtcaagtcttacaaaacttcttgggaagccaaacatttcggacttttatccgagcctggctcgatttga
tttacaagggatagagaggcagatgaaggggttggcaaagcggtttgatggaatatttcagaagatgattgagcagaggttaaagatgcagagagaaaatggaagcgagtccttggacggtggtgaggagaacaaagatttcttgcagtt
tctattgaacttgaaggatgaggaagatgcccagactcagctaaccacgactggcctcaaagctctgctcatggacatgttggttggtggaacagactcatcctccaacacaatcgagtttgccatggctgagatcataaataaaccaaaag
ttctgcaaaacatccagcaagaattggaaactgtggtaggcagaggcaatgtggtagaagaatcgcacatccctaaattgccttacttacaagctgtcatgaaagagtcgctgagattgcacccgccagttcccttgctgatcccacactgc

ccgagtgcgacatgcaccgtcggaggctacactgtcccgaagggctctcgggtcttcatcaatgtgtgggctatacacagggaccctttgatttggagggacccattagagttcgatcctgagaggttcttgcacagtgagggcaattaca
atgagcacaacttcaattacttccctttcgggtctggaagaagaatgtgcgtgggaatattgatggcggagaggatggtgctatactcgctcgccacgctcctgcactcctttgattggaagctgcccaagggggagaagatggatctaaca
gagcaatttgggattgtcatgaagaagaaaaagcctctaatggctgttccgtcgccgaggttctctaattctcggctttatgaatga

>CYP706M1 (NCBI Number: JX518290)

atggacatgagcacaatatggtactactgggtgagtataatcttgggtgttttcatatttctgattgtgggtattcaaaaatggcgatcaaagaaacttcctccggggccttttgcattgcccctgttagggcatcttcatctgcttgagccaaatgt
ccatgaatgcctctcaaagatctctgagaaatttgggcctctcatgtccttcaaatttggcatgaaaacctcaatcatagtctcctcccctgcaatggcgaaggagattctgagagaaaatgaccagatatttgcaaatagaagcattcctgttgt
tgccagatgcattgcatatgatgcatctgatattctgtggagccctaatggacccagatggcgtttactcagaaaaatctgtgttaaggagctttttagccccaagagcactgaggccctgcagcctctgagaagagaggaagtgagaagaa
caatggggaatatttataaggactccattaatggcgtcagtgttgatgttggggcaaaggcctttattacttcactgaatctgattacaaatatgatgtggagtacgagtactgagactggcgaaagagggggggaatttaaggatcttgttgg
ggaacttgttcatgttcttggtgtgcctaatgcttctgatcttttcccctttcttgagagatttgatgttcaggggctttacaggaggatggagaaggtttttgtgagatttgataagatgtttgatgggattattgaggataaattgagtgggaagagt
aaggagaaggatttttttacagtctttgcttgatctggttgaaagaggtgtggatgaacaggatcctgatagcgttcagctcaccatgaaggatgttaaagttctcttgatggacatggtgacaggatcaacagatacaacatccaacacagtgg
aatgggcaatggcagagcttttacagcagccagagataatgaaaagagcccaaaaagaattggaagaagttgtagggcttgacaacatggtagaagaatgccacctgtcccaactcccatacttggacataatagtgaaggaagttctaa
ggctccaccctgcattgcctctgcttgcaccacacagaccagaaagggagtgtgagattggagggtacatcattccaaaggacacccaagtgctgatcaatgtgtggagcattcagaggaacccaaaagtgtggaaggagccactgttg
tttgatccagagaggttttccgattcaaagtgggattacaatggaagggattttgactactttccatttgggtcagggagaagaatctgtgcagggctctccatggcaaaaataatggtgcattattcattggcttctcttctgcattcatttgattgg
tctcttcctgtggcggagaagcttaacatggatgagaagtatggaattgtgcttcgcaaggctgttcctcttgttgccctgcctaagcctcgtttgttgtaccctaatctctatgaatag

### Deep learning DNA sequences

**>Design11361**

ATGGCTTCTAATGAATTGGCCTTCTCCGCTTTGTTGGTTACTTTGGTCTTGGTTCTCATTTCTTGGTACAAGAGAGAAATCTCCAATTCTAGAAAG
GCTGGTACTCCACCATTGCCACCAGGTCCAAAGGGTTTACCTTTGGTCGGTTTCTTGCCATTCTTGGGCCCCAACCTGCATTTGGACTTGTTGAA
GATGGCCAACAACTACGGTCCAATCTTCAAGTTATACTTGGGTTCCAACCTGCACATCGTAGTCAACTCTGCTGATTTGGCCAAGGTTGTTACCG
GTGAACAAGACGAATCCTTCGCTAACCGTGCTCAACACATTGCTGGTTTGGCTACCAGTTACAACGCTTCCGACATCGCCTTCGCCGATAACAAT
GCTAACAGAAGAAAATTGAGAAAGGTTTTGGTCCACGAAGTTTTGTCTAATGTTAACTTGGAAGCTTCCAACGCTTACAGAAGAAGAGAAGTTA
GAAAGACTATCAAGAACGTTCACGAAATTATCGGCAACGAAGTCGATATTAACGAAATTGCCTTCTCTACTGTGTTATCTGTCTTGACATCCATCG
TCTTTGGTAAGTCCATGGTCAAGGGTGCTAAGTATTCTAATTTGGTTGCTGACATGAGAAAGTTCGTCTCCGGTGTTGTTGAGATTGCTGGTGGTT
TGAACATCTCTGACTTCTTCCCAGTTCTTGCCAGATTCGATTTCCAAGGTGTCAAGAGAAAGATGGCAGATCAAATGAAGATGTTCGACAAGATT
TTTGAAACCTCAGTCGAAGAAAGAATCAACTCGCGTTCTGCAATTATTGAAGAAACTGTTAAGCAAGAAGGTCGTAAGGACTTTTTGCAAATCT
TGTTAGAACTGTTGGACCAAAACACTGAAACCTCCATCACCATGACTCAATTGAAAGCTTTGGTGGTTGACATTTTCTTGGGTGGTACCGATGCC
ACCTCCGCCATGGTTGAATGGGCTATGACCGAAATCTTCAGAGATAAAAAGGTCATGAAGAGAGTTCAAGACGAATTAGCTGAAGTTGTCGGTT
TGAACAACATTGTTGAAGAATCTCACTTGCCAAAGTTGAAGTATCTAGATGCTGTCTTCAAGGAAACTTTCAGATTACATCCACCATTGCCTTTTC
TATTGCCAAGAGCTCCAAACAAGTCTTGTACTGTCGGTGGTTACACCGTCCCAAAGGGTTCTACCATCTTTTTGAACGTCTGGGCTATCCAAAGA
GACCCACAATACTGGGAAAACCCATCCGATTTCAACCCAGAAAGATTCTTAAACTACAAGGGTTCTAACAAATGGGACTACGCTGGTACTAACTT
GAAATTCTTTCCATTCGGTTCTGGTCGTAGAAGATGTCCAGGTGTTTCTCTCGGTGAAAAGATGTTGATGCACATATTGGCTTCTTTGTTACACTC
TTTCGACTGGTCTTTGCCAACTGGTCAAAAGCTAGACTTGTCTGATAAGTTCGGTATCACTTTAAAGAAACGTAAGCCATTGATTGCTGTTCCATC
CCCAAGATTGTCTGATGCTTCCTTGTACTTGTAA

**>Design33380**

ATGGCTTCTAACGAACTGGCTTTCTCTGCTTTGTTGGTCACTTTGGTTTTGGTCTTGATCTCTTGGTACAAGAGAGAAATCTCCAACAGTCGTAAG
GCTGGTACCCCACCATTACCTCCATCACCAAAGTCCTTGCCAATTGTCGGTCACTTGCCATTCTTGGGTACTGATATTCACCACGAATTGACTGAA
ATCTCTCACCAATACGGTCCAATCTTCAAGTTCCATCTCGGGTCAAAATTGCATATCATCATTAACTCCGCTGAATTGGCCAAGGTTATCACTGTCG
AACAAGATGAATCTTTCGCTAACAGATGGCCACACATTGCCGGTATCGCCACATCTTATGGTGGTAACGACATTGCTTTTGCTCCAAACAACGCTA
ACTGGAGAAACTTGAGAAAGGTCTTAGTCCAAGAAGTCTTGTCTAACGTTAACTTGGAAGCTTCCCACGCTTACAGAAGACACGAAGTCAGAA
AGGCTGTTAAGTACGTTTACGACAGAGTTGGTATGGATGTTGATATCAATGAAATCTCTTTTTCTACTGTTTTGAACGTCTTCTCCAACATTATCTG
GAGAAAGGGTTTCGTCGACGATGGTACCAACTACGCCAATCTATCCGAAAAGATTCAAAAGGTCATCTGTCGTATTGTCGAAATTGCTGAAGGTT
TGAACATCTCTGACTTCTTCCCAATGTTGGCTAGATTCGATTTGCAAGGTGTTGAACGTAAGATGAAGACCCAAATGAAGCAATTCGACAAGATC
TTCGAAACCACCGTCGATGAAAGAATTAACTCTAAGCCAGCTATTTCTGAAGAAGCTGTCAAGGAAGAAGGTAGAAAAGATTTCTTGCAAATCT
TGTTGGAACTTCTGGACCAAAACACTGCTACCTCCATCACCATGACTCAAATGAAGGCCTTGGTTGTTGACGTTTTTCTAGGTGGTACCGACGCT

ACTAGTGCTATGACTGAATGGGCCATGACCGAGATTTTGAGAAACAGACAAGTTATGAAAAAGGTTCAAGACGAATTGGCTGAAGTTGTCGGTT
TGAACAACATTGTTGAAGAATCCCATCTACCAAAGTTGAAGTATTTGGACGCCGTTTTCAAGGAAACTTTCAGATTGCACCCACCTTTGCCATTC
TTGTTGCCAAGAGCTCCAAACAAGTCTTGTACCGTTGGTGGTTACACTGTTCCAAAGGGTTCCACTATCTTTTTAAATGTCTGGGCTATTCAAAG
AGACCCACAACACTGGACTAACCCATCTGAATTCAACCCAGAAAGATTCTTAAACAAAGGCTCTGAAAAGTGGGACTACAACGGTACCAATTCC
AAGTACTTCCCTTTCGGTTCTGGTCGTAGAAGATGTCCAGGTATCCCATTGGGTGAAAAAATGATGATGCACATTTTAGCCTCTTTGATGCATTCC
TTTGACTGGTCTTTACCAAGAGGTGAAGAACACGATTTATCCGACAAATTCGGTATTGCTATGAAGAAGAAGATGCCATTGGTTTTGATACCATCT
CAAAGATTATCCGATCACAACTTGTACATGTAA

**>Design58683**

ATGGCTTCCAACGAATTGGCTTTCTCTGCTTTGTTGGTCACCTTGGTTTTGGTATTGATCTCTTGGTACAAGAGAGAAATCTCGAATTCCAGAAAG
GCTGGTACTCCACCTTTGCCTCCAGGTCCAAAAGGTTTGCCAGTTGTCGGTTTCTTACCATTTTTAGGCCCAAACTTACATTTGGACTTCCTGACG
TTAGTCCACAAGTACGGTCCAATCTTCAAGTTGTATTTAGGTTCCAACCTACATATCGTCGTTAACTCTGCCGATTTGGCCAAGGTTGTTACTGGT
GAACAAGATGAAAGTTTTGCTAACCGTGCTCAACACATTGCAGGTTTGGCTACCTCTTACAACGCTTCTGATATCGCTTTCGCTGACAACAATGC
CAACAGAAGAAAATTGAGAAAAGTCTTGGTTCATGAGGTTCTGTCCAACGTTAATTTGGAAGCTTCAAATGCGTTCAGACGTCGTGAAGTCAGA
AAGACCATCAAGAACGTTCACGAAATCATTGGTAACGAAGTTGACATCAACGAAATTGCTTTCTCTACCGTCTTGTCCGTTTTGACCTCCATCGT
TTTCGGTAAGTCCATGGTCAAGGGTGCTAAGTACTCCAACTTGGTTGCCGAGATGAGAAAATTCGTTTCTGGCGTGGTTGAAATTGCCGGTGAAT
TGAACATTTCCGATTTCTTTCCAATGTTGGCTAGATTTGATTTCCAAGGTGTTAAGAGAAGAATGGCCCAACAAATGAAGATCTTCGACAGAATTT
TCGAAACAACTATTGAAGAAAGAACCGGTTCTACCTCCGGTATTATTGATCAAAAGGTCAAGGAAGAAGGTAGAAAGGATTTCTTGCAAATCTT
GTTGGAATTATTGGACCAAAACACTGGTACCTCAATCACTATGACCCAATTGAAGGCTTTGGTCGTCGACATTTTTTTGGGTGGTACTGACGCTAC
TTCCGCCATGGTTGAATGGGCTATGACTGAAATATTCAGAGATAAGAAGGTTATGAAAAGGGTTCAAGACGAATTAGCTGAAGTTGTTGGTTTGC
ACAACATTGTCGAAGAATCTCACTTGCCAAAGTTGAAATACTTGGATGCTGTCTTCAAGGAAACCTTCAGATTGCACCCACCATTGCCTTTCTTG
TTGCCAAGAGCTCCAAACAAGTCTTGTACTGTCGGTGGTTACACTGTTCCAAAGGGTTCCACTATCTTTTTGAACGTCTGGGCTATCCAAAGAGA
CCCAAGATACTGGGAAAACCCATCCGACTTCAACCCAGAAAGATTCTTGAACTACAAGGGTTCTGAAAAGTGGGACTACTCTGGTACCAACTTG
AAGTTCTTCCCATTCGGATCTGGTAGACGTAGATGTCCAGGTGTCCCATTGGGTGAAAAGATGATGATGCACATCTTAGCTTCTTTGTTGCACTCT
TTCGACTGGTCTTTGCCAACCGGTGAAAAGTTAGACCTCTCTGACAAGTTCGGTATCACTTTAAAGAAGCGTAAGCCATTGATTGCTATTCCATCT
ATGAGATTATCTGATGCCAGCCTGTACATGTA

**>Design6444**

ATGGCTTCTAATGAACTGGCTTTCTCCGCTTTGTTGGTTACTTTGGTCCTTGTCTTGATTTCTTGGTACAAGAGAGAAATCAGCAACTCTAGAAAG
GCTGGAACTCCACCATTACCTCCAGGTCCTAAGGGTTTACCACTTGTTGGTTTCTTGCCATTCTTAGGTCCAAACTTGCATTTGGACTTTTTAACC
ATGGCTCACCAATACGGTCCAATCTTCAAGCTCTACTTGGGTTCCAACTTGCACATTGTCGTCAACTCCGCTGATTTGGCGAAGGTTGTTACCGGT
GAACAAGACGAATCTTTCGCCAACAGAGCCCAACACATCGCCGGTTTGGCTACCTCTTACAACGCTTCCGATATTTTATTTGCTGACAACAACGC
TAACAGACGTAAATTGCGTAAGGTCTTGGTTCATGAAGTGCTAAGTAATGTTAACTTGGAAGCTTCAAATGCTTACAGAAGAAGAGAAGTCAGA
AAGACTATCAAGAACGTTCACGAGATCATTGGTAACGAAGTCGACATCAACGAAATTGCTTTCTCTACTGTTTTGTCTGTCCTAACTTCCATCGTC
TTCGGTAAGTCCATGGTCAAGGGTGCTAAGTACTCTAACTTGGTTGCCGACATGAGAAAATTCGTTTCCGGTGTTGTTGAAATTGCTGGTGGTTT
GAACATCTCTGATTTTTTCCCAATGTTGGCTAGATTCGATTTCCAAGGTGTTAAGAGAAAAATGGCTCAACAAATGAAGATGTTCGATAAGATCTT
CGAATCTACCGTCGAAGAAAGAGTTGGTTCTACCTCTGGTATTATTGAAGAAGCTGTCAAGCAAGAAGGTAGAAAGGACTTCTTGCAAATCTTG
TTAGAATTGTTGGACCAAAACACTGAAACCTCCATCACTATGACGCAAATGAAGGCATTGGTCGTTGACATTTTCTTGGGTGGTACCGATGCTAC
TTCTGCCATGGTTGAATGGGCCATGACTGAAATCTTCAGAGACAAGAAGGTTATGAAACGTGTCCAAGATGAATTGGCTGAAGTTGTCGGTTTG
CACAACATTGTTGAAGAATCCCATTTACCAAAGTTGAAGTATTTGGATGCCGTTTTCAAAGAAACTTTCAGATTGCACCCACCATTGCCATTCTTA
TTGCCAAGAGCTCCAAACAAGTCTTGTACTGTTGGTGGTTACACCGTTCCAAAGGGTTCCACCATTTTTTTGAACGTCTGGGCTATCCAAAGAGA
TCCACAATACTGGGAAAACCCATCTGAATTCAACCCAGAAAGATTCTTAAACTACAAGGGATCTGAAAAGTGGGACTATACTGGTACTAACTTGA
AGTTCTTCCCATTTGGTTCCGGTAGACGTAGGTGTCCAGGTGTCCCTTTGGGTGAAAAGATGATGATGCACATATTGGCCTCCCTTTTGCACTCAT
TTGACTGGTCTTTGCCAAACGGTCAAAAGTTGGATTTGTCTGACAAGTTCGGTATTACCCTAAAGAAGAGAAAGCCATTGATCGCTGTTCCATCC
TTGAGACTGTCTGACGCTTCTTTGTACGTTTAA

**>Design84497**

ATGGCTTCTAACGAATTAGCCTTTTCTGCCTTGCTAGTGACATTAGTCCTAGTTTTGATTTCTTGGTACAAGAGAGAAATCTCCAACTCCAGAAAG
GCCGGTACTCCCCCATTACCTCCAGGTCCAAGAGGTTTGCCTCTTGTCGGTTACTTGCCATTCTTGGGTCCACAACCACACCGTTCTTTGTCTGAA
ATCTCCCACAGATATGGTCCAATCTTCAAGCTTCAATTAGGTACCAAGTTGTGGATTGTTGTCAACTCTGCTGAATTGGCTAAGGTTATTCACGTT
GAGCAAGATGAATCTTTCGCCAACAGAGCCCCACATATCGCCGGTTTAGCCACTTCTTACGGTGGTAACGATATTGCTTTCGCTCCAAACAACGC
TAACCGTCGTAACTTGAGAAAATTGTTGGTTCAAGAAGTTTTATCTAACGTTAACTTGGAAGCTTCCCACGCTTACAGAAGACACGAAGTCAGA
AAGGCTGTCAAGTACGTTTACGATAGAGTCGGTATGGACATTGATATCAATGAAATTAGCTTCACCACTGTCATCAACGTTTTCTTCAACATCTGG
AGAATGGGTTTCATCGATGACCAATCCAATATTGGTAACTTGTTGGAAAAGATTCAAAAGGTTATCTGTAGAATCGTCGAAATTGCTGAAGGTTT
GAACATTTCTGACTTCTTCCCAGTTTTGGCTAGATTCGATTTGCAAAGAGTTGAACGTAAGATGAAAGACCAAATGAAGCAATTCGACAAAATCA
TTGAAACTACCATCAAGGAAAGAATGAACTCCAAGTCCACTAACGTAGAAGAAACCGTCGAACAAGAAGGTAGAAAGGATTTCTTGCAAATCT
TGTTGGAATTGTTGGATCAATCTACTGCAACCTCTATTACCATGACCCAATTAAAGGCTTTGGTCGTTGATGTTTTCTTGGGCGGTACCGACGCTA
CTTCCGCCATGGTTGAATGGGCTATGACTGAAATTTTGAGAAACAGACAGGTTATGAAGAAGGTCCAAGACGAATTGGCTCAAGTCGTCGGTTT
GCACAACGTTGTTGAAGAATCCCATTTGCCAAAGTTGAAATATTTGGACGCTGTTTTCAAGGAAACTTTCAGACTTCATCCACCATTGCCATTTCT
ATTACCAAGAGCTCCAAACAAGTCTTGTACTGTTGGTGGTTACACTGTTCCAAAGGGTTCCACCATCTTTTTGAACGTCTGGGCTATCCAAAGAG
ACCCACAACACTGGACCAATCCTTCTGAATTCAATCCAGAAAGATTTTTGAACTACAAGGGTTCTGAAAAATGGGACTACGCTGGTACCAACTCT
AAGTTTTTCCCATTGGGCTCCGGTCGTAGAAGATGTCCAGGTATCCCATTGGGTGAAAAGATGATGATGCACATTTTGGCTTCCTTGTTGCACAGT
TTCGACTGGTCATTGCCAACTGGTCAAAAGTTGGACTTGTCTGACAAGTTCGGTATCGCTATGAAGAAAAAGAAGCCATTAGTTGTTGTCCCATC
TTTGCGTTTGTCAGATTTGTCCTTGTACTCTTAA

**>Design91808**

ATGGCTTCCAACGAATTAGCTTTCTCTGCTTTGTTGGTCACTTTGGTTTTGGTTTTGATTTCTTGGTACAAGAGAGAAATCTCAAACTCCAGAAAG
GCTGGTACTCCACCTCTACCACCAGGTCCAAAGGGATTGCCAATTGTCGGTTACTTGTTGTTCTTGGGCACCAACTTGCACATTCACTTCTCCAA
CTTATCTCAATCTTACGGTCCGATCTTCAAGTTTCACTTGGGTAACAAGTTGTGGGTTATCGTCAACACTGCTGAATTGGCTAAGACCATTGTCGT
CGAGCAAGATGAATCTTTCGCTAACAGATGGCCACACATCGCTGGTTTGGCTACCTCCTACGGTGGTAACGACATTGCATTTGCTCCAAACAACG
CCAACAGAAGAAACTTGAGAAAAGTATTGGTCCAAGAAGTTTTATCCAACGTTAACTTGGAAGCTTCTCATAACTACAGAAGACACGAAGTCAG
AAAGGCTGTTAAGTACGTCTACGACAGAGTCGGTATGGATATTGACATCAACGAAATCTCCTTCACCACCGTTTTGAACGTTTTCTTCAACATCAT
CTGGAGAATGGGTTTCGAAGATGACCAAACCAATGTTGGTAACTTGTTGCAAAAGATTCAAAAGGTCATCTGTAGGATCGTTGAAATTGCTGAA
GGTTTAAATATCTCTGACTTCTTCCCAGTCTTGGCCAGATTTGATGTTCAAAGAGTTGAACGTAAGATGAAGGACCAAATGAAGCAATTCGATAA
GATTATTGAAACAACTATCAAGGAACGTATGAACTCTAAGTCTACTAATGTCGAAGAAACTGTTGAACAAAAAGGTCGTAAAGATTTCCTCCAAA
TCTTGCTTGAATTGTTGGACCAAAACAATGAAACCAGTATCACCATGACTCAAATGAAGGCCTTGGTTGTTGATATCTTCTTGGGTGGTACCGATG
CTACTTCTGCCATGGTTGAATGGGCTATGACTGAAATTTTGCGTAACAGACAAGTTATGAAAAAGGTTCAAGACGAACTCGCCGAAATTGTTGGT
TTGAACAACATAGTCGAAGAATCCCACTTACCAAAGTTGAAGTATTTAGACGCTGTCTTTAAGGAAACTTTCAGATTGCATCCACCACTACCTTT
CTTACTTCCAAGAGCTCCAAACAAGACTTGTACTGTCGGTGGTTACACCGTTCCAAAGGGTTCCACTATCTTCTTGAACGTCTGGGCCATTCAAA
GAGACCCACAATACTGGGATAACCCTTCCGAATTCAACCCAGAAAGATTCCTCAACTATAAGGGTTCTGAAAAGTGGGACTACAACGGTACCAA
CTTGAAATTCTTCCCATTCGGTTCTGGTAGAAGAAGATGTCCAGGTATCCCATTGGGTGAAAAAATGATGATGCATATCTTAGCCTCCTTGATGCA
CTCTTTCGACTGGTCTTTGCCAAGAGGTGAAGAATTGGATTTGAGCGACAAGTTTGGGATTGCTATGAAGAAGAAGAAGCCATTGGTTGTCATTC
CATCTTTGCGTTTATCGGACCACAACTTGTACATGTAA

**>Design49566**

atggcttctaacgaattggctttttcagcattgttggttactttggttttggttttgatttcttggtacaagagagaaatctctaactcaagaaaggcaggtacaccaccacccctgcctccaggtccaaagggtttaccattggttggttccttgcca
ttcttgggtccaaacatccatcaagaattgaccaagatcactcaccaatacggtccaattttcaagttatacttgggttctaaattgcacatcgttgtcaactccgctgaattggctaaggttatcaccgctgagcaagacgaatcttttgctaata
gagctccacatatcgccggtttggctacttcttacggtggtaacgacattgctttcgccccaaacaacgctaacagaagaaacttgagaaaggttttagttcaagaagttttatccaacgttaatttggaagcttctcacgcttacagaagacac
gaagtcagaaaggctgtcaagtacgttcacgaaagagtcggtatggaagtcgatatcaacaaaattgccttttccacagtcttgtccgttttgactaacattgtctgggctaagagcgtcgtcgatgatggtgccaactactctaatgaagttca
aaacatcatttctagagttgtcgaaattgctggtggtttgaacatttccttcttcgtcatggctcaaattgatttccaaggtgttgaaagaagaatgaagaaacaaatgaagcaattcgacaagataatcgaaatgaccattgaagaacgtatgaa
ctcttctaagaacgaagccaagactaaggaagaaggtagaaaggatttttgcaaatcttgttggaacaacaacaacaaaacactgaaacctccatcaccatgactcaaatgaaagctttggtcgttgacatctttttgggtggtactgatgct
acctctgctatggttgaatgggccatgactgaaatcttgagagaccaacaagttatgaagcgtgtccaagatgaattagaagaaattgttggcttaaacaacgtcgaagaatcccacttgccaaagttgaaatatttggacgccgtcttcaag
gaaactttcagactccatccaccaattccattcttgctaccaagagctccaatcaagtcttgtaccgtgggtggttacactgttccagaaggtgctaccatcttcgttaacgtttgggctattcaacgtgacccacaaagatgggaaaacccatc

tgaattcaacccagaccgtttcttgaacagaaatggatctactggtaagtgggactactccggtaccaacttgaccttcttacctttcggttctggtcgtagaagatgtccaggtattccactaggtgaaaagatgatgatgcacattttggcctc
acttttgcactccttcgactggaagttgccaaacggtcaaaagttggaattgtctgaaaggttcggtatcgctttgaagaagaagaagccattggtagctgttccaaccaagagattagatgttagtttgtacatgtaa

**>Design12854**

ATGGCTGCTTTGAACACCCACGGTAGTTGGTGGCCAGCTGAAGGTAACGGTGGTAAGAACGACGGTGACCTCCCATTAGCTTTGTTAGCTGTTAT
CACAGCTGCCTTATTGCCATTGTTATGGTACAAGAGATCCATCTCCTCTTCTCAAAACGGTGCCCCACCACTGCCGCCAGGTCCAAAGGGTTTAC
CAGTTGTCGGTTACCTTCCTTTCTTGGGCCCTAACTTGCACTTGGACTACTTGACCATGGTGCACCAATACGGTCCAATTTTCAAGATTTACTTGG
GCTCAAACTTACATATCGTGGTCAACTCTGTCGATTTGGCCAAGGTTGTCACTGGTGAACAAGATGAATCCTTTGCTAACAGAGCTCAACACATC
GCCGGTTTGGCTACATCTTACAACGCTTCCGACATCGCTTTCGCTGATAACAACGCCAACAGAAGAAAGTTGAGAAAGGTTTTGGTTCACGAAG
TCTTGTCCAACGTTAACTTGGAAGCTTCTAATGCTTACAGACGTAGAGAAGTTAGAAAGACTATCAAGAACGTTCACGAAGTCATCGGTAACGA
AGTCGATATCAATGAAATTTCTTTCTCTACTGTTTTGTCCGTTTTGACTTCTATCGTTTTCGGTAAGTCCATGGTTAAGGGTGCTAAGTACTCTAAC
CTCGCCGCTGACATTCGTAAATTTGTTTCTGGTGTCGTTGAAATTGCTGGTGGTCTAAACATCTCTGACTTCTTTCCAATGTTAGCTAGATTCGATT
TCCAAGGTGTTGAACAAAGAATGAAGACCCAAATGAAGATGTTCGACAAGATTTTCGAAACCTCTGTCGAAGAAAGAATCAACTCCAGATCTG
CTATCAAAGAAGAAGCCGTCAAGGAAGAAGGTAGAAAGGACTTCTTGCAAATCTTGTTGGAATTGTTGGAACAAAATACTGAGACCAGTATTAC
TATGACTCAAATGAAGGCTCTGGTCGTTGACGTCTTTTTGGGTGGTACCGATGCTACCAGTGCTATGGTTGAATGGGCCATGACCGAAATCTTCA
GAAACAGACAAGTTATGAAAAAGGTCCAAGACGAATTGGCCGAAATCGTCGGATTGCACAACATTGTCGAAGAATCTCATTTGCCAAAGTTAAA
ATATTTGGACGCTGTCTTCAAGGAAACTTTCAGATTGCACCCACCATTGCCATTCTTATTGCCAAGAGCTCCAAACAAGACTTGTACCGTTGGTG
GTTACACTGTTCCAAAAGGTTCAACCATTTTCTTGAACGTTTGGGCTATTCAAAGAGACCCAAAGTACTGGGACAACCCATCCGATTTCAACCCA
GAACGTTTTTTGAACTACGAAGGTGAAAAATGGGATTATAATGGTACTAACCTAAAGTTCTTCCCATTCGGTTCCGGTAGAAGAAGATGTCCTGG
TATCCCATTGGGTGAAAAGATGATGATGCACATTTTGGCTTCCTTGTTGCACTCTTTCAACTGGTCTCTACCAGAAGGTGAAGACCACGATTTGTC
TGAAAAGTTCGGTATTGCTATGAAGAAGAAGAAGCCATTGATTGCCATTCCATCTTTGCGTTTGAGCGATCATAACTTGTACATGTAA

**>Design61021**

atggcttctaacgaattggctttttcagcattgttggttactttggttttggttttgatttcttggtacaagagagaaatctctaactcaagaaaggcaggtacaccaccattaccaccaggtccatacggcttgccactagtcggttatttgccattct
tgggtccttccctacaccacgaattgactaagatggcccacagatacggtccaatcttcaagttgtacttgggttccaaattgcacattgttgttaattctgctgatctggctaaggttatcacttccgaacaagatgaatcctttgctaacagagc
tccacacattgctggtttggccacatcttacggtgctaacgacattgctttcgctgacaacaacgccaacagaagaaacttgagaaagatcttggtccacgaaatcttgtccaacgttaacttggaagcttctcacgcatacagaagaagaga
agttagaaagactatcaagagtgttcatgacatgattggtatgccagttgatattaacgaaatgtctttctccactgttgttaacgtattgacctctattgtctggggtaactccatggttgaaggtaccaagcattctaacctaggtgaagaaataa
gaaaggttgtctctgaaatcgtcgatattgctgaaggtttgaacatctctgacttcttcccaaaattagctagattcgacttgcaaggtgtcgaacaaaagatgaagagaaagatgaagcaatttgactggattttcgaaaccaccattgaagaa
cgtatcaacttgaagtctacccacggtgaagatgccctaaagcacgaaggtcgtaaggatttcttgcaaatcttattggaattgaaggacaagaaatcaatcaccatgactcaattgaaggctctcgtcgttgacatcttcttgggtggtactga
tgctacttctgccatggttgaatgggctatggctgagattttgaagaaccaaaaggttatgaaaaaggtccaagacgaattagccgaaatcgttggtttaaagaacatggtcgaagaatctcatttacctaaattaaagtacttgaatgctacctt
caaggaaactttcagattgcataccccactaccagtcttgttgccaagaactccatctaagagttgtatggtcggtggttacttgatcccaagagattctactgtcttcttaaatgtctgggccattcaaagagacccacaacactgggaaaacc
catccgaattcaacccagaaagatttttgaactacgaaggttcaggtaagtgggactactctggtaccaactccaagtacttcccattcggttccggtcgtcgtagatgtccaggtattccattggctgaaaagatgatgttgcacatcctggct
tccttgttgcactctttcgattggtctttgccgaagggtgaagaccacgatttgtttgaaaagttcggtatcgctttgaaaaagaagaagccattggttgctgttccatctccaagattgattgacttgtctttatacatgtaa

**>Design33105**

atggcttctaacgaattggctttttcagcattgttggttactttggttttggttttgatttcttggtacaagagagaaatctctaactcaagaaaggcaggtacaccaccattgccaccaggtccaagaggtttacctttggttggttatttgccattctt
aggcccaaacttgcaccaagaattgaccaagatggctcacagatacggtccaattttcaagttgtacttgggttctaaactacacatcgttgttaattctgctgatttggctaaggttgttaccggtgaacaagacgaatctttcgcgaaccgtgc
tccacatattgccggtttggccacttcttacaacgcttccgatatcgccttcgctgacaacaatgccaacagaagaaaattgcgtaaggtcttggtccacgaagttttgtctaatgtgaacttggaagcttctcacgcttacagaagaagagaag
ttagaaagactatcaagaacgttcatgaaatcatcggtaacgaagttgacataaatgaaattgctttctcaaccgtcttgtccgtccttacctctatcgtctggggtaagtccatggttaagggtgcttccaacatgattgttgaagtcagaaagtt
cgtttccggtgttgtcgagattgctggggaattaaacatctctgatttttttcccaatgttggctagattcgatttccaaggtgtcgaaagaagaatgaagaagcaaatgaaattgtttgacaagattttcgaatccactgttgaagaacgtatcaac
agcagatctatcattaaggaagaagctgcttccaaagaagaaaaccgtagaaaggactttttgcaaatcttgttagaattacaagaacaaaacaacacttccatcaccatgactcaaatgaaggctttggttgtcgacatcttcctgggtggta
ccgatgctacctccgccatgattgaatgggctatgaccgaaattttgagaaaccgtagagttatgaaaaaggtccaagatgaattggctgaaattgtcggtttgcaaaacaacgttgaagaagaaagtcatctcccaaagttaaagtacttgg
acgctgtcttcaaagaaactttcagattgcacccacctctacctttcttgttgccaagagccccaaacaagtcttgtactgtcggtggttacaccatcccaaagggttctactatctttttgaacgtctgggctattcaaagagacccacaatactg
ggaaaacccatccgaattcaacccagaacgtttcttgaactacaagggttctgaaaagtgggactacgctggtactaactctaagtttttcccattgggttctggtagaagaagatgtccaggtgtttctttaggtgaaaagatgatgatgcaca
tcttggcctctttgttgcactccttcgactggtccttgccaactggtcaaaagttggatctatctgacaaattcggtatcgctttgaagaagagaaagccattgattgctgttccatctccaagattgaacgatgcctccttatacatgtaa

**>Design42565**

atggcttctaacgaattggctttttcagcattgttggttactttggttttggttttgatttcttggtacaagagagaaatctctaactcaagaaaggcaggtacaccaccattgccaccaggcccaaagggtctaccattggttggttacttgcctttct

tgggtccaaatttacatcacgaattcactaaggtttctcacagatacggtccaattttcaagttgtatttgggttccaagctccacattgttgttaacagcgctgatttggccaaggtcatcacttctgaacaagacgaatctttcgccaaccgtgcc
ccacacattgctggtttggccacctcttacggtggtaacgatattgctttcgctgacaacaacgctaacagaagaaacttaaggaaggtcttggttcatgaagttttgtcaaacgtcaacttggaagcttcccacgcttacagacgtcacgaag
tcagaaagactattaagtccgtccacgacatgatcggtatggaagttgatatcaacgaaatttccttctctactgttttgaacgttttgaccaacattgtttggggtaagggtttagttgaaggtaccaaatactccaacttgtctgaagaaatccg
gaaggttgtctacagaattgtcgaaattgctgaaggattgaacatttctgatttttcccaatgttggccagatttgatttacaaggtgtcgaaagaaagatgaaaaaacaaatgaagcaattcgatagaatcttcgaaaacactatctctgaaag
aactaacaacaagaactctaaccacggtaagcaatgtttgcaaatcttgttggaattgaagaccaagaccatcaccaccacacaattgaaggctttggtcgttgacatcttcttaggtggtaccgatgctacttccgccatggttgaatgggct
atgactgaaattttgagaaacaagaaggtaatgaagagagttcaagatgaattggaagaaatcgttggtttgaacaatatcgtcgaggaatcccatatcccaaagttgaagtacttggacgctgtcttcaaggaaactttcagattgcacccac
ctctgcctttttttattgccaagagctccatctaagtcttgtactgtcggtggttacaccgttccaaaaggtgctaccatcttcctaaatgtctgggctatccaaagagacccaagacactgggaaaatccatctgaattcaacccagacagattcc
ttaacaacaacaacggttcaactgaaaaatgggactactctggtactaacttgactttttgccattcggttccggtagacgtagatgtccaggtatcccattaggtgaaaagatgatgatgcatatattagcttctttgatgcactccttcgactgg
agtttgccaaacggtgaagaacacgatttgtccgacaagttcggtattgcattgaaaaagaagaagccattggttgctattccaaccagaagattgtctgacgaaaatttgtacatgtaa

**>Design26159**

atggcttctaacgaattggctttttcagcattgttggttactttggttttggttttgatttcttggtacaagagagaaatctctaactcaagaaaggcaggtacaccaccattgccaccaggaccatacggtctgccattgttgggttatttgcctttttt
cttgggcccttccttgcaccacgaattgaccaagatggctcacagatacggtccaatcttcaaactgtatctaggttctaagttacacattgtcgtcaactctgctgatttggctaaggtcatcactagtgaacaagacgaatcttttgctaacag
agctccacacatcgctggtctagccacttcttacggtgctaacgacattgctttcgctgacaacaacgccaaccgtagaaacttgagaaaaattttggtccatgaaattctatctaacgttaacttggaagcttcccacgcttacagaagaaga
gaagtgcgtaagaccatcaagtccgttcacgacatgattggtatgccagttgacattaacgaaatgtctttctctactgtcgtaaatgttttgacttccatagtctggggtaactccatggttgaaggtaccaagcattccaacttgggtgaagaa
atcagaaaggttgtctccgaaattgttgatattgccgaaggtttgaacatctctgatttcttcccaaaattagctagattcgatttgcaaggtgttgaacaaaagatgaagagaaagatgaagcaattcgactggattttcgaaaccaccattgaa
gaaagaatcaacctgaagtctactcacggtgaagatgctctcaagcacgaaggtcgtaaggatttcttacaaatcttattggaattgaaagacaagaagtccattaccatgactcaattgaaggctttggttgtcgacatctttttaggtggtact
gacgctacctccgccatggtcgaatgggccatggccgaaattcttaagaaccaaaaggttatgaaaaaagttcaagatgaattggctgaaatcgttggtttgaagaacatggttgaagaatcccatttgccaaagttgaagtacttaaacgct
accttcaaggagactttcagattgcacactccattgccagtcttgttgccaagaactcctagcaagtcttgtatggtcggtggttacttaattccaagagattccactgttttcttgaacgtctgggctatccaaagagacccacaacactgggaa
aacccatctgaattcaatccagaaagattcttaaactacgaaggttccggtaagtgggactacagtggtacaaattcaaagtacttcccattcggttctggtagaagaagatgtccaggtatcccgttggcagaaaagatgatgttgcatatctt
ggcctctttgttgcactctttcgactggtctttgccaaagggtgaagaccacgatttgtttgaaaagttcggtattgctttgaaaaagaagaagccattagttgctgttccatctccacgtttgatcgatttgtctttgtacatgtaa

**>Design49566**

atggcttctaacgaattggctttttcagcattgttggttactttggttttggttttgatttcttggtacaagagagaaatctctaactcaagaaaggcaggtacaccaccacccctgcctccaggtccaaagggtttaccattggttggttccttgcca
ttcttgggtccaaacatccatcaagaattgaccaagatcactcaccaatacggtccaattttcaagttatacttgggttctaaattgcacatcgttgtcaactccgctgaattggctaaggttatcaccgctgagcaagacgaatcttttgctaata
gagctccacatatcgccggtttggctacttcttacggtggtaacgacattgctttcgccccaaacaacgctaacagaagaaacttgagaaaggttttagttcaagaagttttatccaacgttaatttggaagcttctcacgcttacagaagacac
gaagtcagaaaggctgtcaagtacgttcacgaaagagtcggtatggaagtcgatatcaacaaaattgccttttccacagtcttgtccgttttgactaacattgtctgggctaagagcgtcgtcgatgatggtgccaactactctaatgaagttca
aaacatcatttctagagttgtcgaaattgctggtggtttgaacatttccttcttcgtcatggctcaaattgatttccaaggtgttgaaagaagaatgaagaaacaaatgaagcaattcgacaagataatcgaaatgaccattgaagaacgtatgaa
ctcttctaagaacgaagccaagactaaggaagaaggtagaaaggatttttttgcaaatcttgttggaacaacaacaacaaaacactgaaacctccatcaccatgactcaaatgaaagctttggtcgttgacatctttttgggtggtactgatgct
acctctgctatggttgaatgggccatgactgaaatcttgagagaccaacaagttatgaagcgtgtccaagatgaattagaagaaattgttggcttaaacaacgtcgaagaatcccacttgccaaagttgaaatatttggacgccgtcttcaag
gaaactttcagactccatccaccaattccattcttgctaccaagagctccaatcaagtcttgtaccgtgggtggttacactgttccagaaggtgctaccatcttcgttaacgtttgggctattcaacgtgacccacaaagatgggaaaacccatc
tgaattcaacccagaccgtttcttgaacagaaatggatctactggtaagtgggactactccggtaccaacttgaccttcttacctttcggttctggtcgtagaagatgtccaggtattccactaggtgaaaagatgatgatgcacattttggcctc
acttttgcactccttcgactggaagttgccaaacggtcaaaagttggaattgtctgaaaggttcggtatcgctttgaagaagaagaagccattggtagctgttccaaccaagagattagatgttagtttgtacatgtaa

**>Design58**

atggcttctaacgaattggctttttcagcattgttggttactttggttttggttttgatttcttggtacaagagagaaatctctaactcaagaaaggcaggtacaccaccacttcctccaggtccaaagggtttgccattggttggttacttgccattctt
gggtccaaacttgcaccaagaattaaccaagatggctcacagatacggtccaatcttcaagttgtatttgggttccaagttgcacatcgttgtcaactcagctgatttggccaaggttgttactggtgaacaagacgaaagtttcgctaacaga
gctccacatattgctggtttagctacctcttacaacgcttctgacattgctttcgcggacaacaacgccaacagacgtaagttgagaaaggtcctcgtccatgaagttttgtccaacgttaacttggaagctagtcacgcttacagaagaagag
aagtcagaaagaccatcaagaacgtccacgaaatcatcggtaacgaagtcgacattaacgaaatcgccttctctactgtcttgtccgtcttgacctctattgtttggggtaagtccatggttgaaggtgctaagtactctaacttggttgctgaaa
tgagaaaattcgtttctggtgttgtcgaaatcgccggtgagttgaacatctccgacttcttcccaatgttggctagattcgatttccaaggtgttgaaaggagaatgaagaagcaaatgaaattgtttgacaagatctttgaatctaccgttgaaga
aagaatcaactcccgtaaatctgtcgaaaaggaagatttttttgcaaatcttgttggaattgcaagaacaaaacaatgaaacctccatcactatgactcaaatgaaggctttggttgtcgatatcttcttgggtggtactgatgctacttctgctatg
attgaatgggccatgactgaaattttaagaaaccgtcaagttatgaagaaagttcaagatgaattggcagaaattgtcggtttgaataatattgtggaagaatcccacctaccaaaattgaagtacttagacgctgttttcaaggaaactttcaga
ttacacccaccattgcctttcttgctcccaagagccccaaacaagtcttgtactgttggtggttacactattccaaagggttctaccattttcttgaatgtctgggctatccaaagagacccacaatactgggaaaacccatccgaattcaaccca
gaacgtttttttaaactacaagggctctgaaaagtgggactacgctggtaccaactctaagtttttcccattaggttccggtagaagaagatgtccaggtgtttcactaggtgaaaagatgatgatgcacattttagcttccttgttgcattctttcga
ctggtctttgccaaccggtcaaaagttggacttgtctgataagttcggtatcgctttgaaaaagagaaagccattgattgctgtcccatctccaagattgaacgatgcctctttatacatgtaa

**> Design4779**

atggcctctaatgaattggctttctccgctttgttagttactctggtcttagtcttgatttcttggtacaagagagaaatctccaactctcgtaaggccggaactccaccattgccaccaggtccaagaggtttgccattggttggttacttgccattc

ttaggtccaaacttgcaccaagaattgaccaagatggctcacagatacggtccaatcttcaagttgtacttgggttctaaattacatatcgttgttaactctgctgacttggccaaagtcgtcactggtgaacaagacgaatcttttgctaaccgtg
cccctcacattgctggcttggctacttcttacaacgcttccgatatcgccttcgctgacaacaacgcaaacagaagaaaattgcgtaaggtccttgtccacgaagtcttgtcaaacgtcaacttggaagcttctcacgcttacagaagaagag
aggttagaaagaccatcaagaacgttcacgaaattattggtaacgaagttgacattaacgaaattgctttctcaaccgttttgagcattctcacttccattgtttggggtaagtccatggttgaaggtgctaagtacactaatttggttgctgaaatg
agaaagttcgtttccggtgttgtggaaatagccggtgaattaaacatttccgatttttcccaatgttggccagattcgatttccaaggtgttgaaagaagaatgaagaagcaaatgaaattattcgacaagattttcgaatccactgttgaagaaa
gaatcaactcctccaagaaggaaggtagaaaggacttcctacaaatcttgttggaattgcaagaacaaaacaatgaaacctccatcaccatgactcaaatgaaggctttggttgtcgacatctttttgggtggtacagatgctacctctgctat
ggtcgaatgggctatgactgaaatcttgagaaacagacaagttatgaagaaagttcaagatgaattagctgaaattgtcggtttgaacaacatcgtcgaagaatctcatttgccaaagttgaaatatttggacgctgttttcaaggaaactttca
gattacatccacctctgccattcttgttaccaagagctcctaacaagtcttgtaccgtaggtggttacaccattccaaagggttctactattttttgaatgtctgggctatccaaagagacccacaatactgggaaaacccatctgaattcaaccc
agaaaggttcttgaactacaagggttctgaaaagtgggattatgctggtaccaactccaagttcttcccattgggttccggtcggcgtagatgtccaggtgtctctttgggtgaaaagatgatgatgcacatccttgcttccttgctacactctttc
gactggtctttgccaaccggtcaaaagttggatttgtctgacaaattcggtatcgccttgaagaagagaaagccattgatcgctgtcccatccccaagattgaacgatgcctctttgtacatgtaa
**>Design3644**
atggcttctaacgaattggctttttcagcattgttggttactttggttttggttttgatttcttggtacaagagagaaatctctaactcaagaaaggcaggtacaccaccactcccaccaggtcctagaggtttgccactggttggttacttgccattct
tgggtccaaacttgcaccacgaattgaccaagatggctcacagatatggtccaatcttcaagttatatcttggttctaagctacatatcgttgttaacagcgctgatttggctaaggttgtcaccggtgaacaagacgaatcttttgccaacagag
atccacacatcgctggtttagctacctcatacaacgctaacgatattgcattcgctgacaacaacgccaaccggagaaaattgagaaaggttttggtccacgaagtcttatccaacgtcaacttggaagcttctcacgcttacagacgtagag
aagttcgtaagacaattaagaacgttcatgaaatcatcggtaacgaagtcgacattaatgaaatcgctttctctactgttttgtcagtcttgacttccattgtttggggtaagtccatggttaagggtgctaagtacccaaacttagtcgctgaaatg
agaaagttcgtttccggtgtcgttgaaatcgccggtgaattgaacatttctgacttcttcccaatgttggccagatttgatttccaaggtgttgaaagaagaatgaagaagcaaatgaaattgttcgataagatcttcgaatccactgttgaagaa
agaattaactctagaaagaagattgctcgtaaggactttttgcaaatcttgttggaattgaaagaacaaaacaacgaaacctctatcaccatgacccaaatgaaggccctggtcgttgacatcttcttgggaggtaccgatgccacttctgctat
gattgaatgggctatgactgaaattctcagaaacagacaagttatgaagaaggttcaagatgaattagctgaaattgtcggcttgaataacatcgtcgaagaatctcatttgccaaagttaaagtacttggatgctgttttcaaggaaactttcag
attacacccacctttgcctttttgttaccaagagccccaaacaagtcttgtaccgtcggtggttacactatcccaaagggttcgactattttcttgaatgtctgggctatccaaagagacccacaatactgggaaaacccatctgaattcaaccc
agaaagattcttgaactacgaaggttccgagaagtgggactacgctggtactaactccaagttcttcccattgggttccggtagacgtagatgtccaggtatcccattaggtgaaaagatgatgatgcacatcttggcttccttgatgcactctt
tcgactggtctttgccagaaggtgaaaaattggacttgtccgataaattcggtattgccttgaagaagaaaaagccattgattgctgttccatctccaagattgaacgacgcttctttgtacatgtaa
**>Design2205**
atggcttctaacgaattggctttttcagcattgttggttactttggttttggttttgatttcttggtacaagagagaaatctctaactcaagaaaggcaggtacaccaccattgccaccaggtccaagaggtttaccactcgtaggatacttgcctttc
ttgggtccaaacctacatcacgaattggacaagatggctcacagatacggtccaatcttcaagttgtatttgggttctaagttgcacatcgttgtcaacagcgccgacttggcaaaggttgtcactggtgaacaagacgaatccttcgctaatc
gtgctccacacattgctggtttggctacctcttacaacgcttcagacattgcgtttgctgacaacaacgccaacagaagaaagttgagaaaggtgttggttcacgaagttctatctaacgttaacttggaagcttcccacgcttacagaaggag
agaagtcagaaagactattaagaacgttcatgaaattatcggtaatgaagttgacattaatgaaattgctttctctactgttttgtctgttttgacttccattgtctggggtaagtccatggttaagggtgctaagtacggtaacttggttgctgaaatg
agaaaattcgtttctggtgttgttgaaatcgccggtgaattgaacatctctgatttttcccaatgttggctagatttgatttccaaggtgtcgaacgtagaatgaagaaacaaatgaagttgttcgataagatattcgaaaccaccgtcgaagaaa
gaaccaactcctccaagaagaaggtcaagaaggacttcttgcaagtcttgttggaattaaaggaacaaaacaacgaaacatccatcaacatgactcaaatgaaggctttggtcgtcgatatctttttgggtggtaccgacgctacttctgctat
gattgaatgggccatgaccgaaatcctaagaaacaagcaagttatgaaaaaagtccaagatgaattggctgaagttgttggtttgaacaacattgttgaagagtctcacttgccaaaattgaaatacttagatgccgtcttcaaggaaactttca
gattgcaccctccattgccattcctgttgccacgtgcccctaacaagtcttgtactgtcggtggttacaccatcccaaagggttccaccatcttcttaaacgtctgggctatccaaagagacccacaatactgggaaaacccaagtgaattcaa
cccagaaagattcttgaactacaagggttccgaaaagtgggactacgctggtactaactccaaattcttcccattgggctctggtcgtagaagatgtccaggtgtttccttgggtgaaaagatgatgatgcatatcttagcttctttattacactct
tttgactggtctttgccagaaggtcaaaagcttgatttgtctgacaagttcggtattgccttgaagaagaagaagccattgatcgctattccatctccaagattgaacgatgcctctttgtacatgtaa
**>Design4129**
atggcttctaacgaattggctttttcagcattgttggttactttggttttggttttgatttcttggtacaagagagaaatctctaactcaagaaaggcaggtacaccaccattgccaccaggtccaagaggtttgccattagttggttacttgccattct
tgggtccaaacttgcaccaagaattgaccaagatggctcacagatacggtccaatcttcaagttgtatttgggctccaagttgcatattgttgtcaactctgctgatttggccaaggtggtcaccggtgaacaagatgaatcttttgctaaccgtg
ctccacacattgccggtttggctacttcttacaacgcttcagacattgctttcgctgacaacaatgccaacagacgtaaattaagaaaagtcttggtccatgaagtcttgtccaacgttaacttggaagcctcccacgcttacagaagaagaga
agttcgtaagaccattaagaacgttcacgaaatcatcggtaacgaagttgatattaacgaaatcgctttctccactgtcttatccgttctaacaagtatcgttttgaagggttctaacattgctgctgaaatgagaaagttcgtttccggtgtcgttg
aaattgctggtgaattgaatatctctgatttttcccaatgttggctagatttgatttccaaggtgttgaaagaagaatgaagaagcaaatgaaattgttcgataagattttcgaatcaactgttgaagaaagaatcaactccagatctgctattaag
gaggaagctgtcaaggaagaaggtagaaaggacttcttgcaaatcttattggaattgcaagaacaaaacaacgaaacctcgataaccatgacccaaatgaaggctctagttgttgacatcttcttaggtggtactgatgctacctctgccatg
attgaatgggctatgactgaaattttgagaaacagacaagttatgaagaaggttcaagacgaattggcagaaattgtcggtttaaacaacattgtcgaagaatcccatttgccaaaactaaagtacttggacgccgttttcaaagaaactttcag
attgcacccacctttgcctttttgttaccaagagccccaaacaagtcttgtactgtcggtggttacactatcccaaagggttccactatcttcttgaatgtctgggccatccaaagagacccacaatactgggaaaacccatctgagttcaaccc
agaaagattcttaaactacaagggttccgaaaagtgggactacgctggtaccaacagcaaattcttcccattgggttctggtcggagaagatgtccaggtgtctctttgggtgaaaagatgatgatgcacatcttggcttccctattgcactctt
tcgactggtctttgcctactggccaaaagttggacttgtctgacaagttcggtatcgctttgaagaagaggaagccattgatcgctgttccatctccacgtttgaacgatgcttctctttacatgtaa

### The Rosetta scripts and options for RosettaLigand

**ligand_dock.options**

```
-in:file:s ancX_HEM_AGI.pdb
-in:file:extra_res_fa inputs_CPD1/HEM.params inputs_CPD1/AGI.params
-run::preserve_header
-packing
	-ex1
	-ex2aro
	-ex2
	-no_optH false
	-flip_HNQ true
	-ignore_ligand_chi true
-enzdes
	-cstfile inputs_CPD1/HEM.cst
-parser
	-protocol inputs_CPD1/ligand_dock.xml

-out
	-path:all outputs_CPD1
	-nstruct 50
	#-overwrite
```

**ligand_dock.xml**

```
<ROSETTASCRIPTS>
	<SCOREFXNS>
		<ligand_soft_rep weights="ligand_soft_rep">
		</ligand_soft_rep>
		<hard_rep weights="ligand">
		</hard_rep>
	</SCOREFXNS>

	<LIGAND_AREAS>
		<docking_sidechain_X chain="X" cutoff="6.0" add_nbr_radius="true" all_atom_mode="true"
minimize_ligand="10"/>
		<final_sidechain_X chain="X" cutoff="6.0" add_nbr_radius="true" all_atom_mode="true"/>
		<final_backbone_X chain="X" cutoff="7.0" add_nbr_radius="false" all_atom_mode="true"
Calpha_restraints="0.3"/>

		<docking_sidechain_F chain="F" cutoff="6.0" add_nbr_radius="true" all_atom_mode="true"
minimize_ligand="10"/>
		<final_sidechain_F chain="F" cutoff="6.0" add_nbr_radius="true" all_atom_mode="true"/>
		<final_backbone_F chain="F" cutoff="7.0" add_nbr_radius="false" all_atom_mode="true"
Calpha_restraints="0.3"/>
```

```
	</LIGAND_AREAS>
	<INTERFACE_BUILDERS>
		<side_chain_for_docking ligand_areas="docking_sidechain_X,docking_sidechain_F"/>
		<side_chain_for_final ligand_areas="final_sidechain_X,final_sidechain_F"/>
		<backbone ligand_areas="final_backbone_X,final_backbone_F" extension_window="3"/>
	</INTERFACE_BUILDERS>
	<MOVEMAP_BUILDERS>
		<docking sc_interface="side_chain_for_docking" minimize_water="true"/>
		<final sc_interface="side_chain_for_final" bb_interface="backbone" minimize_water="true"/>
	</MOVEMAP_BUILDERS>
	<SCORINGGRIDS ligand_chain="F" width="15">
		<classic grid_type="ClassicGrid" weight="1.0"/>
	</SCORINGGRIDS>
	<SCORINGGRIDS ligand_chain="X" width="15">
		<classic grid_type="ClassicGrid" weight="1.0"/>
	</SCORINGGRIDS>
	<MOVERS>
	single movers_X
		<AddOrRemoveMatchCsts name="cstadd" cst_instruction="add_new"/> add catalytic constraints
		<Transform name="transform_F" chain="F" box_size="7.0" move_distance="0.2" angle="20" cycles="700" repeats="1" temperature="5"/>
		<Transform name="transform_X" chain="X" box_size="8.0" move_distance="0.2" angle="20" cycles="700" repeats="1" temperature="5"/>
		<AddOrRemoveMatchCsts name="cstrem" cst_instruction="remove" keep_covalent="1"/> remove constraints
		<HighResDocker name="high_res_docker" cycles="6" repack_every_Nth="3" scorefxn="ligand_soft_rep" movemap_builder="docking"/>
		<FinalMinimizer name="final" scorefxn="hard_rep" movemap_builder="final"/>
		<AddOrRemoveMatchCsts name="cstfinadd" cst_instruction="add_pregenerated"/>
		<InterfaceScoreCalculator name="add_scores" chains="X,F" scorefxn="hard_rep"/>

	compound movers
		<ParsedProtocol name="low_res_dock">
			<Add mover_name="cstadd"/>
			<Add mover_name="transform_F"/>
			<Add mover_name="transform_X"/>
			<Add mover_name="cstrem"/>
		</ParsedProtocol>
		<ParsedProtocol name="high_res_dock">
			<Add mover_name="high_res_docker"/>
			<Add mover_name="final"/>
			<Add mover_name="cstfinadd"/>
		</ParsedProtocol>
	</MOVERS>
```

```
	<PROTOCOLS>
		<Add mover_name="low_res_dock"/>
		<Add mover_name="high_res_dock"/>
		<Add mover_name="add_scores"/>
	</PROTOCOLS>
</ROSETTASCRIPTS>
```

**heme.cst**

```
#block 1 for covalent bond for CYS and CPD1

CST::BEGIN
  TEMPLATE::   ATOM_MAP: 1 atom_name: FE1 N2 N1
  TEMPLATE::   ATOM_MAP: 1 residue3: HEM

  TEMPLATE::   ATOM_MAP: 2 atom_type: S     ,
  TEMPLATE::   ATOM_MAP: 2 residue1: C

  CONSTRAINT:: distanceAB:    2.30   0.20 180.00   1
  CONSTRAINT::    angle_A:   90.00   3.00 100.00  360.00
  CONSTRAINT::    angle_B:  110.00  10.00  50.00  360.00
  CONSTRAINT::  torsion_A:  -80.00   5.00  50.00  360.00
  CONSTRAINT::  torsion_B:   90.00   5.00  25.00  360.00
  CONSTRAINT:: torsion_AB:  -80.00   5.00   5.00  360.00
CST::END
```
